# Joint dimension reduction and clustering analysis for single-cell RNA-seq and spatial transcriptomics data

**DOI:** 10.1101/2021.12.25.474153

**Authors:** Wei Liu, Xu Liao, Yi Yang, Huazhen Lin, Joe Yeong, Xiang Zhou, Xingjie Shi, Jin Liu

**Affiliations:** Academy of Statistics and Interdisciplinary Sciences, East China Normal University, Shanghai, 200062, China; Centre for Quantitative Medicine, Health Services & Systems Research, Duke-NUS Medical School, Singapore 169857, Singapore; Center of Statistical Research and School of Statistics, Southwestern University of Finance and Economics, Chengdu, 611130, China; Institute of Molecular and Cell Biology(IMCB), Agency of Science, Technology and Research(A*STAR), Singapore, 138673, Singapore; Department of Biostatistics, University of Michigan, Ann Arbor, 48109, USA; Key Laboratory of Advanced Theory and Application in Statistics and Data Science-MOE, School of Statistics, East China Normal University, Shanghai, 200062, China

**Keywords:** dimension reduction, clustering, expectation-maximization algorithm hidden Markov random field, spatial transcriptomics, scRNA-Seq

## Abstract

Dimension reduction and (spatial) clustering is usually performed sequentially; however, the low-dimensional embeddings estimated in the dimension-reduction step may not be relevant to the class labels inferred in the clustering step. We therefore developed a computation method, Dimension-Reduction Spatial-Clustering (DR-SC), that can simultaneously perform dimension reduction and (spatial) clustering within a unified framework. Joint analysis by DR-SC produces accurate (spatial) clustering results and ensures the effective extraction of biologically informative low-dimensional features. DR-SC is applicable to spatial clustering in spatial transcriptomics that characterizes the spatial organization of the tissue by segregating it into multiple tissue structures. Here, DR-SC relies on a latent hidden Markov random field model to encourage the spatial smoothness of the detected spatial cluster boundaries. Underlying DR-SC is an efficient expectation-maximization algorithm based on an iterative conditional mode. As such, DR-SC is scalable to large sample sizes and can optimize the spatial smoothness parameter in a data-driven manner. With comprehensive simulations and real data applications, we show that DR-SC outperforms existing clustering and spatial clustering methods: it extracts more biologically relevant features than conventional dimension reduction methods, improves clustering performance, and offers improved trajectory inference and visualization for downstream trajectory inference analyses.

## 1 Introduction

Single-cell RNA sequencing (scRNA-seq) studies encompass a set of widely applied technologies that profile the transcriptome of individual cells on a large scale and can reveal cell subpopulations within a tissue [1, 2]. Spatial transcriptomics studies, on the other hand, involve a series of recently developed technologies that allow for the simultaneous characterization of the expression profiles of multiple tissue locations while retaining their locational information. scRNA-seq technologies include full-length transcript sequencing approaches (e.g., Smart-seq2 [3] and MATQ-seq [4]) and 3’/5’-end transcript sequencing technologies (e.g., Drop-seq [5] and STRT-seq [6]). While spatial transcriptomics technologies include earlier fluorescence in situ hybridization (FISH)-based approaches (e.g., seqFISH [7] and MERFISH [8]) and sequencing-based techniques (e.g., 10x Visium [9] and Slide-seq [10]) among others. Both scRNA-seq and spatial transcriptomic technologies have provided unprecedented new opportunities to characterize the cell type heterogeneity within a tissue, investigate the spatial gene expression patterns [11, 12], explore the transcriptomic landscape of the tissue, identify spatial trajectories on the tissue [13], and characterize the spatial distribution of cell types within tissues and across multiple tissue types [14–16].

In the analysis of both scRNA-seq and spatial transcriptomics datasets, dimension reduction and (spatial) clustering are two key analytical steps that are critical for many downstream analyses such as cell lineage analysis and differential expression analysis. Specifically, due to the curse of dimensionality, dimension-reduction methods are usually applied to the transformation of the original noisy expression matrix in either scRNA-seq or spatial transcriptomics into a low-dimensional representation before performing (spatial) clustering analysis [13, 17–19]. The existing literature describes many dimension-reduction methods that have been developed and common methods include principal component analysis (PCA), weighted PCA (WPCA) [20], t-distributed stochastic neighbor embedding (tSNE) [21], uniform manifold approximation and projection (UMAP) [22], etc. PCA is a well-recognized approach that is routinely used in many software packages used for both scRNA-seq and spatial transcriptomics analyses [17, 23] and has many desirable features such as simplicity, computational efficiency, and relative accuracy. For example, Seurat [24], SpaGCN [25], BayesSpace [26] and SC-MEB [27] all first extract the top principal components (PCs) from the high-dimensional expression matrix and then perform (spatial) clustering analysis. WPCA is a variation of PCA that imposes different weights on different genes to upweight the potentially informative genes [20] in the presence of heteroscedastic noises. SpatialPCA is another variation of PCA that incorporates spatial localization information to encourage neighborhood similarity in the PC space [13]. While fitting PCA, WPCA and SpatialPCA is generally automatic and does not rely on parameter tuning, the other two widely used nonlinear dimension-reduction methods tSNE and UMAP rely relatively heavily on the manual tuning of parameters for optimized performance [19, 23]. In addition to these generic methods, several dimension-reduction methods have been developed that account for either the count nature and/or dropout events of scRNA-seq data, e.g., zero-inflated factor analysis (ZIFA) [28] zero-inflated negative binomialbased wanted variation extraction (ZINB-WaVE) [29], and single-cell variational inference tools (scVI) [30].

After obtaining a low-dimensional representation with dimension reduction, (spatial) clustering analyses are then carried out. Clustering of scRNA-seq data aims to identify cell types and cluster cells into the distinct cell categories. Spatial clustering in spatial transcriptomics aims to use spatial transcriptomic information to cluster tissue locations into multiple spatial clusters, effectively segmenting the entire tissue into multiple tissue structures or domains. Cell-type clustering facilitates our understanding of the cellular composition of tissues with potentially heterogeneous cell types, whereas spatial clustering facilitates the characterization of the tissue structure and is a key step towards understanding the spatial and functional organization of tissue. Common clustering methods for scRNAseq analysis include *k*-means [31] and the Gaussian mixture model (GMM) [32]. Common spatial clustering methods for spatial transcriptomics analysis include the graph convolutional network (GCN)-based approach SpaGCN [25], the hidden Markov random field model (HMRF) implemented in the Giotto package [33], BayesSpace [26], SC-MEB [27], and SpatialPCA [13], all of which promote the smoothness of cluster assignments in neighboring tissue locations. By performing dimension reduction and (spatial) clustering sequentially, the estimated low-dimensional embeddings and class labels can be used for many types of downstream analyses, such as cell lineage analysis [34–37], spatial trajectory inference on the tissue [13], and differential gene expression (DGE) analysis [38].

The majority of existing methods for dimension reduction and (spatial) clustering have been used in a tandem analysis by first performing dimension reduction on expression matrix followed by (spatial) clustering analysis of the estimated low-dimensional embeddings [19], as shown in Fig. 1a. Performing dimension reduction and (spatial) clustering as two sequential analytical steps is not ideal for two important reasons. First, these tandem methods optimize distinct loss functions for dimension reduction and (spatial) clustering separately, and the two loss functions may not be consistent with each other when aiming to achieve optimal (spatial) cluster allocation [39]. PCA aims to retain as much variance as possible in as few PCs as possible, whereas spatial clustering aims to either minimize within-cluster variances or maximize between-cluster variances. Second, the dimension-reduction step in the tandem methods does not consider uncertainty in obtaining low-dimensional features. Consequently, the extracted low-dimensional components are effectively treated as error-free in the spatial clustering analysis, which is not desirable. To address these two drawbacks of tandem analysis, several recent methods have been developed in other research areas to perform joint dimension reduction and clustering analysis. For example, an ad-hoc remedy would perform two analytical steps iteratively: estimating the low-dimensional embeddings by applying supervised dimension reduction together with the inferred latent class labels (the Dimension-Reduction step;DR), and inferring class labels using estimated embeddings, and using, if necessary, spatial information (the Spatial-Clustering step;SC). These simple procedures echo some recent explorations of self-supervised learning [40, 41], in which deep neural networks were combined with simple classifiers to perform unsupervised clustering of image data. To some extent, joint methods employ self-learning to classify all spots and obtain latent features iteratively. However, it is still challenging to unify the existing methods and combine both DR and SC steps in a self-learning manner.

**Figure 1:**
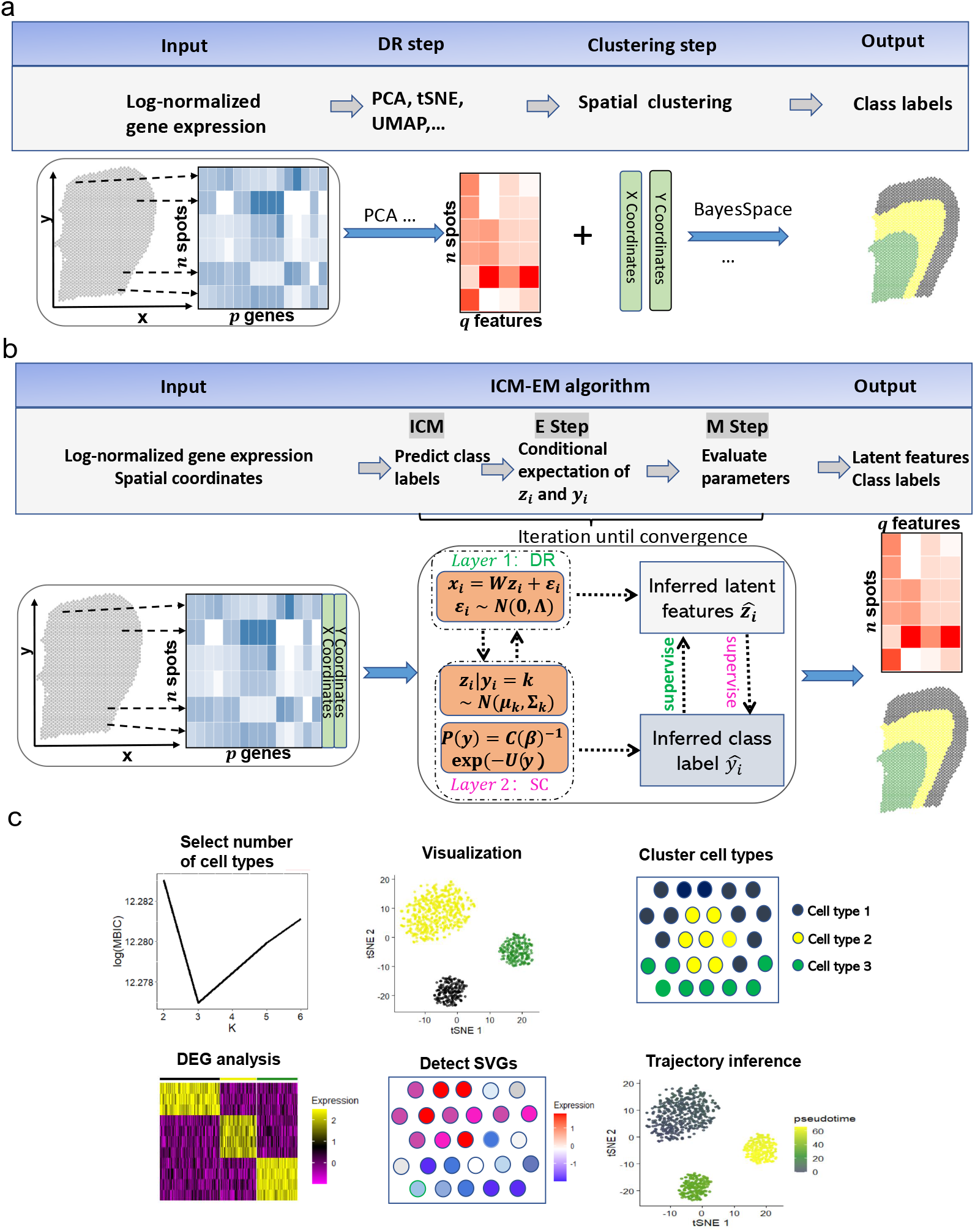
Workflows for both tandem analysis (a) and DR-SC (b) and potential applications of DR-SC in downstream analysis (c). a & b. Compared with tandem analysis, DR-SC iteratively performed dimension reduction and (spatial) clustering with improved estimation for both clustering and low-dimensional embeddings. c. DR-SC can be used to cluster cell types, with the number of clusters selected in a data-driven manner. The estimated cell types can be used to perform differential gene expression analysis. The estimated low-dimensional embeddings from DR-SC can be used for visualization, trajectory inference, and detection of gene expression with spatial variations by controlling cell-type-relevant covariates.

Here, we propose a unified and principled method to both estimate low-dimensional embeddings relevant to latent class labels and, in the case of spatial transcriptomics analysis, further leverage these embeddings with spatial information to perform spatial clustering using an HMRF. The proposed method was built on a hierarchical model with two layers, as shown in Fig. 1b: the first layer relates gene expression to low-dimensional embeddings and represents the DR step; while the second layer relates the latent embeddings to the cluster labels, and, if necessary, spatial information and thus represents the SC step. These two layers are unified in DR-SC such that the relevant features are estimated while simultaneously performing spatial clustering. We developed an efficient expectation-maximization (EM) algorithm based on an iterative conditional mode (ICM) [42, 43]. DR-SC is not only computationally efficient and scalable to large sample sizes but is also capable of optimizing the smoothness parameter in the spatial clustering component. Importantly, when the smoothness parameter is set to zero, DR-SC directly performs clustering for scRNA-seq data with no spatial information. Unlike existing spatial clustering approaches, DC-SR can determine the number of clusters in an automatic fashion using modified Bayesian information criteria (MBIC) [44]. Using 16 benchmark scRNA-seq datasets, we demonstrated that the low-dimensional embeddings and the class labels estimated from DR-SC lead to better performance in the downstream lineage analysis using Slingshot [36]. We further illustrated, using both CITE-seq and spatial transcriptomics (10x Visium and Slide-seqV2) datasets, that DR-SC achieves higher (spatial) clustering accuracy and resolves low-dimensional representations with improved visualization. To exemplify the utility of the estimated low-dimensional embeddings from DR-SC, we performed analysis to infer cell lineages using a seqFISH mouse embryonic dataset. The R package DR.SC is available on CRAN (https://CRAN.R-project.org/package=DR.SC), with functions implemented for standalone analysis and Seurat based [45] pipeline analyses.

## 2 Materials and Methods

### 2.1 Model Specification

We proposed the use of DR-SC to estimate low-dimensional latent features while improving clustering performance via a unified statistically principled method. DR-SC relates to a two-layer hierarchical model that simultaneously performs dimension reduction via a probabilistic PCA model and promotes spatial clustering using an HMRF based on empirical Bayes. With spatial transcriptomics datasets, we observe a *p*-dimensional log-normalized expression vector *x_i_* = (*x*_*i*1_,⋯, *x_ip_*)^*T*^ for each spot, *s_i_* ∈ ℝ^2^, on square or hexagonal lattices, while its class label, *y_i_* ∈ {1,⋯, *K*}, and *q*-dimensional embeddings, **z**_*i*_’s, are unavailable. Without loss of generality, we assume that **x**_*i*_ is centered and DR-SC models the centered log-normalized expression vector **x**_*i*_, with its latent low-dimensional feature, **z**_*i*_, and class label, *y_i_*, as

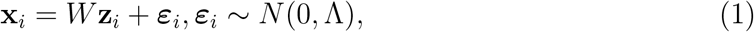

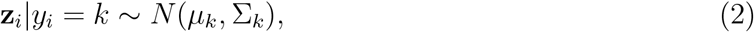

where Λ = diag(λ_1_,⋯, λ_*p*_) is a diagonal matrix for residual variance, *W* ∈ ℝ^*p*×*q*^ is a loading matrix that transforms the *p*-dimensional expression vector into *q*-dimensional embeddings, and *μ_k_* ∈ ℝ^*q*×1^ and Σ_*k*_ ∈ ℝ^*q*×*q*^ are the mean vector and covariance matrix for the *k*th class, respectively. Eqn. (1) relates to the high-dimensional expression vector (**x**_*i*_) in *p* genes with a low-dimensional feature (**z**_*i*_) via a probabilistic PCA model while Eqn. (2) is a GMM for this latent feature among all *n* spots. When spatial coordinates (*s_i_*) are available, we assume each latent class label, *y_i_*, is interconnected with the class labels of its neighborhoods via a Markov random field. To promote spatial smoothness within spot neighborhoods, we assume that the hidden Markov random field **y** = (*y*_1_, ⋯,*y_n_*)^*T*^ takes the following Potts model [46],

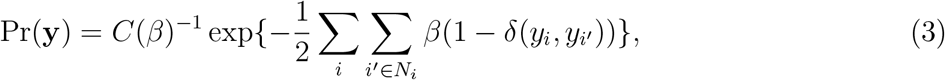

where *δ* is a Dirac function, *C*(*β*) is a normalization constant that does not have a closed form, *N_i_* is the neighborhood of spot *i*, and *β* is the smoothing parameter that controls the similarity among the neighboring labels, in other words, the degree of spatial smoothness. When this smoothing parameter *β* goes to zero, the spatial-clustering step in DR-SC, Eqn. (2) and (3), is reduced to a latent GMM with no spatial information.

DR-SC unifies both models for dimension reduction and (spatial) clustering (Fig. 1b). By combining the latent GMM in Eqn. (2) and the Potts model in Eqn. (3), DR-SC performs the spatial clustering on low-dimensional embeddings obtained from the probabilistic PCA model in Eqn. (1). Conventionally, the embeddings obtained using unsupervised dimension reduction methods, such as PCA, UMAP, or tSNE, reflect variations caused by different sources, including batch effects and microenvironments among the observed cells/spots, etc., other than cell-type differences. Thus, embeddings from unsupervised dimension reduction analyses may distort the downstream clustering used for cell typing [47]. In contrast, DR-SC performs dimension reduction in a self-learning manner, where the embeddings, **z**_*i*_’s, are estimated under the supervision of the estimated latent labels for each spot (Fig. 1b). Thus, the obtained embeddings capture information with regards to biological differences, e.g., cell-type or cellstate differences, which in turn improve the spatial clustering for cell typing. When no spatial information is available, as in scRNA-seq, we can simply apply a latent GMM (2) without considering the Potts model (3).

### 2.2 Compared methods

We conducted comprehensive simulations and real data analysis by comparing the dimension reduction and clustering performance of DR-SC with those of existing methods. In detail, we considered the following eight dimension-reduction methods to compare the dimensionreduction performance: (1) PCA implemented in the R package *stats*; (2) WPCA [48] implemented in the R package *DR.SC*; (3) factorial *k*-means (FKM) [39] implemented in the R package *clustrd*; (4) tSNE; (5) UMAP, in which tSNE and UMAP were implemented in the R package *scater*; (6) ZIFA implemented in the Python module *ZIFA*; (7) ZINB-WaVE implemented in the R package *zinbwave*; and (8) scVI implemented in the Python module *scvi.* As the last three methods, ZIFA, ZINB-WaVE, and scVI, can be applied to only raw count data, we compared their performance with that of DR-SC in Simulation 2 with the count matrix for expression levels and real datasets.

We considered the following 10 clustering methods when comparing clustering performances. (1) BayesSpace [49] implemented in the R package *BayesSpace*; (2) Giotto [33] implemented in the R package *Giotto*; (3) SC-MEB [27] implemented in the R package *SC.MEB*; (4) SpaGCN [25] implemented in the Python module *SpaGCN*; (5) Louvain [50] implemented in the R package *igraph*; (6) Leiden [51] implemented in the R package *leiden*; (7) GMM implemented in the R package *mclust*; (8) *k*-means implemented in the R package *stats*; (9) FKM [39] implemented in the R package *clustrd*; and (10) subspace clustering based on arbitrarily oriented projected cluster generation (ORCLUS) [52] implemented in the R package *orclus*. In tandem analysis, we used BayesSpace, Giotto, SC-MEB, and SpaGCN, which were recently developed to perform spatial clustering, and Louvain, Ledien, GMM, and *k*-means, which are conventional non-spatial clustering algorithms. We also applied FKM and ORCLUS to perform joint dimension reduction and clustering analysis.

### 2.3 Simulations

We performed two sets of simulations as follows. Simulation 1 involved log-normalized gene expression data. In this simulation, we generated non-spatial/spatial log-normalized gene expressions. In detail, we generated the class label, *y_i_*, for each *i* = 1,⋯, *n* in a rectangular 70 × 70 lattice from a *K*-state (*K*=7) Potts model with smoothing parameter *β* = 0 or 1 using function `sampler.mrf` in R package *GiRaF.* Then we generated latent low-dimensional features, **z**_*i*_;, from the conditional Gaussian, such that **z**_*i*_|*y_i_* = *k* ∼ (*μ_k_*, Σ_*k*_), where **z**_*i*_ ∈ *R^q^* with *q* = 10 and structures for *μ_k_* and Σ_*k*_ are shown in Supplementary Table S1. Next, we generated 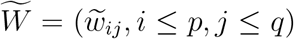 with each 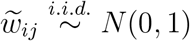, to perform a QR decomposition on 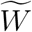 such that 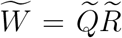, and assigned 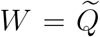, i.e., a column orthogonal matrix. Finally, we generated a high-dimensional expression matrix using **x**_*i*_ = *W***z**_*i*_ + ***ε***_*i*_, ***ε***_*i*_ ~ *N*(**0**, Λ), where Λ = diag(λ_*j*_), *j* = 1,…,*p*. In the case of homoscedasticity, λ_*j*_ = ∀*j*, while in the case of heteroscedasticity, 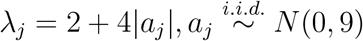.

Simulation 2 involved raw gene expression data. In this simulation, we generated non-spatial/spatial raw gene expressions. The method used to generate the class label, *y_i_*, loading matrix, *w*, and latent features, **z**_*i*_’s, was the same as in Simulation 1, except that *μ_k_* had a different value (see Supplementary Table S1). The difference involved the generation of log-normalized gene expression, **x**_*i*_, using **x**_*i*_ = *W***z**_*i*_ + *τ* + ***ε***_*i*_, *τ_j_* ~ *N*(0,1), ***ε***_*i*_ ~ *N*(**0**, Λ) and raw gene expression, 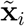, using 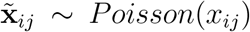, where *τ_j_* is the *j*-th element of *τ*, Λ = diag(λ_*j*_, *j* = 1, ⋯, *p*. To ensure a proper signal, we set λ_*j*_ = 1, ∀_*j*_, in the case of homoscedasticity and 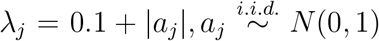 in the case of heteroscedasticity. In this simulation, we only observed raw gene expression 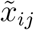 of gene *j* and cell *i* for non-spatial settings and the raw gene expression 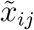 of gene *j* and spot *i* and spatial coordinates *s_i_* for spot *i*.

### 2.4 Real datasets

#### 2.4.1 Human dorsolateral prefrontal cortex datasets

We downloaded spatial transcriptomics obtained on the 10x Visium platform for human dorsolateral prefrontal cortex (DLPFC) from https://github.com/LieberInstitute/spatialLIBD. This dataset is a collection of data from 12 human postmortem DLPFC tissue sections from three independent neurotypical adult donors and the raw data for each sample includes 33,538 genes. We first selected genes with spatial variation using SPARK [11] without adjusting for any covariates. In detail, we selected spatially variable genes (SVGs), either those with adjusted *p*-values of less than 0.05 or the top 2,000 SVGs (Supplementary Table S2), then we performed log-normalization using library size. Detailed information on the 12 samples is given in Supplementary Table S2. Taking the manual annotations based on cytoarchitecture as benchmarks, we were able to evaluate the clustering performance of DR-SC and other methods. In the tandem analysis, we first obtained the top 15 PCs from either PCA or WPCA, then applied other clustering methods using the top 15 PCs [26]. We further performed spatial variation analysis (SVA) to identify SVGs adjusted for cell-type-relevant covariates using SPARK and compared them with SVGs that were not adjusted for these covariates. We then performed DGE analysis using the function *FindAllMarkers* in the R package *Seurat* to identify differentially expressed genes based on cell type labels estimated using DR-SC. Finally, we performed functional enrichment analysis using g:profiler [53] based on the SVGs with adjustment.

#### 2.4.2 Mouse olfactory bulb and mouse E15 neocortex data

We downloaded mouse olfactory bulb or mouse E15 neocortex data from https://singlecell.broadinstitute.org/single_cell/data/public/SCP815/sensitive-spatial-genome-wide-expression-profiling-at-cellular-resolution#study-summary. We first selected the top 2,000 genes with spatial variation using SPARK [11] without adjusting for any covariates. Then we performed log-normalization of these SVGs using library size and obtained the top 15 PCs based on PCA. Because BayesSpace and SC-MEB are both based on tandem analysis, the top PCs obtained from PCA were used as inputs. SpaGCN is also based on tandem analysis but it used its internally embedded PCA algorithm.

For the joint method, DR-SC was applied to the 2,000 SVGs, and we clustered all spots from the mouse olfactory bulb data into 12 clusters and all spots from the mouse E15 neocortex data into 15 clusters. Using the estimated class labels for the clusters from the DR-SC, we performed DGE analysis using the function *FindAllMarkers* in the R package *Seurat* to identify the marker genes for each cluster. Next, we performed cell typing using the PanglaoDB database [54]. Finally, we performed trajectory inference by Slingshot method based on the extracted features and cell classes estimated by DR-SC and detected the differentially expressed genes along the inferred cell pseudotime using the function *testPseudotime* in the R package *TSCAN.*

#### 2.4.3 Mouse embryo datasets

We downloaded a mouse embryo dataset [16] from https://content.cruk.cam.ac.uk/jmlab/SpatialMouseAtlas2020/ that was collated using a sequential fluorescence in situ hybridization (seqFISH) platform. This dataset contained 23,194 cells, 351 genes, and twodimensional spatial coordinates. Cell types were annotated based on their nearest neighbors on an existing scRNA-seq atlas (Gastrulation atlas) [16]. Taking these manual annotations as the benchmark, we compared the clustering performance of DR-SC with that of other spatial clustering methods. We performed log-normalization with library size of the gene-expression matrix. In the tandem analysis, we first obtained the top 15 PCs [26] from either PCA or WPCA and then applied other spatial clustering methods using these topc 15 PCs. We further restricted our analysis to the cells manually annotated as “forebrain/midbrain/hindbrain” and performed downstream trajectory analysis. By applying DR-SC, we obtained six subclusters for the brain region. Then, we performed DGE analysis using the function *FindAllMarkers* in the R package *Seurat* to identify differentially expressed genes among the estimated clusters and further mapped six clusters to either four cortical regions or four cell types using the PanglaoDB database [54]. To visualize the clustering results, we applied tSNE to reduce the 15-dimensional embeddings obtained via different methods to a 2-dimensional representation. Finally, we applied Slingshot to conduct trajectory inference based on the features and clusters from DR-SC and detected differentially expressed genes along the inferred cell pseudotime using the function *testPseudotime* in the R package *TSCAN*.

#### 2.4.4 Benchmark datasets in trajectory inference

We downloaded 16 benchmark datasets with linear trajectory information from the website https://zenodo.org/record/1443566#.XNV25Y5KhaR [55]. These datasets consisted of single-cell gene-expression measurements in the form of raw read counts. Detailed information on these datasets is given in Supplementary Table S3, including the species, number of cells, number of genes, platform, etc. We first pre-processed the raw count data using *Seruat,* which included selecting the top 2,000 most variable genes and log normalized the data using library size [56] for methods based on normalized expression, except ZINB-WaVE and scVI. After normalization, we estimated the low-dimensional embeddings and class labels using both joint and tandem methods. For the joint analysis, we considered the proposed DR-SC and FKM while in tandem analysis, we performed dimension reduction using other methods followed by clustering analysis using the GMM. The number of clusters was chosen using modified BIC and regular BIC by default for DR-SC and GMM, respectively. Because FKM does not select the number of clusters automatically, the number of clusters selected for DR-SC was used for FKM. To further perform lineage development analysis, we applied Slingshot implemented in the R package *slingshot* with default values for parameters using the estimated embeddings and class labels as input. Analysis details and results are deferred to the Supplementary Text (Supplementary Fig. S1 - S3).

#### 2.4.5 Cord blood mononuclear cell datasets

We obtained a cord blood mononuclear cell (CBMC) dataset from NCBI https://www.ncbi.nlm.nih.gov/geo/query/acc.cgi?acc=GSE100866 under the accession number GSE100866. This dataset contains 8,167 CBMCs measured using CITE-Seq technology [57] from two species (human and mouse). In addition to genome-wide expression measurements of 20,511 genes in the form of read counts, this dataset also contains data on the protein levels of 13 cellsurface markers. We first performed preprocessing to retrieve the top 2,000 most variable genes and lognormalized these based on library size [56]. Following the vignette at https://satijalab.org/seurat/archive/v3.1/multimodal_vignette.html [57], we ignored three cell-surface markers and performed clustering analysis with the remaining 10 markers using the *FindClusters* function in the R package *Seurat.* Taking the class labels from the nine clusters estimated using these 10 surface markers as the benchmark, we evaluated the clustering performance of DR-SC and other methods using the adjusted Rand index (ARI) values. We used DR-SC and FKM to simultaneously estimate the embeddings and class labels, while other methods were applied for tandem analyses. Using the estimated class labels for the 11 clusters from DR-SC, we performed DGE analysis using the R package *BPSC* [38] on human cells. Next, we performed cell typing using the PanglaoDB database [54]. For each cell type, we further performed functional enrichment analysis by selecting the significant genes with adjusted *p*-values of less than 0.05 and a log fold-change greater than 0.5. Analysis details and results are deferred to the Supplementary Text (Supplementary Fig. S4 - S8 and Table S4).

### 2.5 Evaluation metric for dimension reduction and (spatial) clustering

We evaluated the performance of DR-SC from four aspects: feature extraction, clustering performance, selection of the number of clusters, and computational efficiency. Here, we briefly present the evaluation metrics for the feature extraction and clustering performance. For details on the other two aspects, please refer to the Supplementary Text.

In the simulations, we used two metrics to assess the performance of the feature extraction including both the canonical correlations between the estimated features and the underlying true ones and the conditional correlation between gene expression, **x**_*i*_, and cell type label, *y_i_*, given the estimated latent features. Canonical correlation measures the similarity between two sets of random variables. Thus, a larger canonical correlation coefficient value suggests a better estimation of **z**_*i*_. For optimal performance, we aimed to obtain the estimated features, 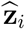, that contain all information on cell types, in other words, 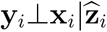 with the smaller the conditional correlation the better.

To compare clustering performance, we evaluated both ARI [58] and normalized mutual information (NMI) [59]. ARI [58] is a corrected version of the Rand index (RI) [60] that avoids some of its drawbacks [58]. The ARI is used to measure the similarity between two different partitions and lies between −1 and 1. The larger the value of ARI, the higher the similarity between two partitions. When the two partitions are equal up to a permutation, the ARI takes a value of 1. NMI is a way of correcting the mutual information (MI) so the NMI value falls between zero and one. MI quantifies the “amount of information” obtained on one random variable in units such as Shannons (bits) by observing another random variable. MI intuitively measures the information that two random variables *x* and yx share. If *x* and *y* do not share information and are independent, then *MI*(*x,y*) = 0. At the other extreme, if *y* = *x*, then *MI*(*x,y*) = *H*(*x*), where *H*(*x*) is the marginal entropy of *x*. This indicates that MI does not take values between zero and one. Thus, some normalized versions have been proposed and we used one of these versions (NMI).

## 3 Results

### 3.1 DR-SC method overview

Here, we provide a brief overview of DR-SC, and further details are available in the Supplementary Text. The proposed method involves simultaneous dimension reduction and (spatial) clustering built on a hierarchical model with two layers, as shown in Fig. 1b. The first layer, the DR step, relates the gene expression to the latent low-dimensional embeddings, while the second layer, the SC step, relates the latent embeddings along with spatial coordinates if necessary to the cluster labels. Unifying the DR and SC steps not only produces relevant low-dimensional embeddings, improving the visualization of the clusters on the tSNE plots, but also enhances the (spatial) clustering performance.

In later sections, we show the improved clustering performance with spatial transcriptomics datasets from different platforms. In the Supplementary Text, we show how DR-SC improved the clustering performance for single-cell datasets. Aside from improving the (spatial) clustering performance, DR-SC estimates low-dimensional embeddings that can also be used in different types of downstream analyses (Fig. 1c). First, the estimated embeddings can be used to better visualize clustering among cells/spots. Second, the performance of the trajectory inference can be improved, as the reduced dimensional space from DR-SC possesses more relevant information on cell clusters. Third, by taking these estimated embeddings as covariates, we can perform hypothesis testing to identify genes with pure spatial variation but not due to cell-type differences. These gene expression differences may be related to a specific cell morphology or tissue type rather than being related to cell type. In the Supplementary Text, we compare the accuracy of the downstream lineage inference using the estimated embeddings and cell-type labels from DR-SC with those from other unsupervised dimension-reduction methods applied to 16 benchmark scRNA-seq datasets. In later sections and in the Supplementary Text, we also show the improvements to clustering performance and cluster visualization provided by DR-SC for both non-spatial (CITE-seq) and spatial transcriptomics (10x Visium, Slide-seqV2, and seqFISH) datasets. The basic information (number of spots/cell/genes and platforms) on the selected spatial transcriptomics datasets are shown in Supplementary Table S5. By applying DR-SC to several spatial transcriptomics datasets, we further show the utility of using the low-dimensional embeddings obtained from DR-SC to identify genes related to cell morphology and tissue type.

### 3.2 DR-SC improves clustering and estimation of low-dimensional features in simulations

We conducted simulation studies to evaluate the performance of DR-SC in comparison with existing dimension reduction and clustering methods. First, we simulated data with both non-spatial (*β* = 0) and spatial (*β* = 1) patterns and with both homogeneous and heterogeneous residual variance *λ_j_* (see Materials and Methods). Two simulation settings were considered. In Simulation 1, log-normalized and centered gene expression data were generated from Eqn. (1) – (3). In Simulation 2, we first generated a count matrix using Poisson distribution with over-dispersion, which more effectively mimics the count nature of scRNA-seq and 10x Visium datasets (see Methods). Then, we log-transformed the raw count matrix using library size [56]. In all simulations, we set *p* = 1,000 and ran 50 replicates. The details of the simulation settings are provided in the Materials and Methods.

We compared the clustering performance of DR-SC with two groups of spatial/non-spatial clustering methods. The first group used tandem analysis using PCs from either PCA or WPCA in the DR step, and using SpaGCN [25], BayesSpace [49], SC-MEB [27], Giotto [33], Louvain [50], Leiden [51], GMM, and *k*-means in the clustering step. Among these, SpaGCN software used the log-normalized expression matrix as the input and its internally embedded PCA algorithm to obtain PCs, and could only be applied to spatial clustering. The second group was a joint analysis with ORCLUS [52] and FKM [39]. By setting the smoothing parameter to zero, BayesSpace, SC-MEB, and Giotto could be applied to cluster non-spatial data. On the other hand, to evaluate the estimation accuracy of low-dimensional embeddings, we compared DR-SC with eight dimension-reduction methods in all simulation settings, including PCA, WPCA [20], FKM [39], tSNE [21], UMAP [22], ZIFA [28], ZINB-WaVE [29], and scVI [30].

We first compared the clustering performances of each method. For tandem analysis in Simulation 1, we applied both PCA and WPCA to obtain low-dimensional embeddings, denoted as suffix -O and -W, respectively in Fig. 2 and Supplementary Fig. S9. In Simulation 2 (Supplementary Fig. S10), besides PCA and WPCA, we also applied ZINB-WaVE to obtain low-dimensional embeddings as the input for different clustering methods in the tandem analysis. Since Giotto, *k*-means, FKM, and ORCLUS do not provide a data-driven way to select the number of clusters, *K*, we evaluated their clustering performances using the true cluster number. DR-SC achieved the best clustering performance and was the most robust to both homogeneous and heterogeneous residual variances among the methods that used the true cluster number (Fig. 2a and Supplementary Fig. S9a and S10a). To select the number of clusters, *K*, DR-SC and SC-MEB used MBIC [44, 61], GMM used BIC, BayesSpace adopted the average loglikelihood-maximization-based method in early iterations, Leiden and Louvain used a community-modularity-maximzing rule [50], and SpaGCN applied Louvain initialization [25]. DR-SC also achieved the best clustering performance among the methods that selected the number of clusters automatically (Fig. 2b, Supplementary Fig. S9b and S10b). Meanwhile, DR-SC selected the true number of clusters consistently (Fig. 2c and Supplementary Fig. S11b). Conventional PCA is unable to recover the underlying features in the presence of heteroscedastic noise, while WPCA can give less informative genes less weight [20]. Thus, when heterogeneous errors appeared, the clustering performance of the tandem analysis using conventional PCA was worse than that using WPCA. In all settings, the clustering performance of DR-SC was robust in both homogeneous and heterogeneous cases. Importantly, DR-SC achieved the highest ARI values among all the methods trialed. Moreover, we observed that only DR-SC correctly chose the number of clusters. In constrast, BayesSpace tended to overestimate the cluster number in non-spatial cases and underestimate them in spatial cases; this is because BayesSpace fixed the smoothing parameter rather than updating it in a data-driven manner. Thus, the selection of the number of clusters for BayesSpace was sensitive to the choice of smoothing parameter (Supplementary Fig. S11c). The other methods tended to show similar patterns across both non-spatial and spatial cases.

**Figure 2:**
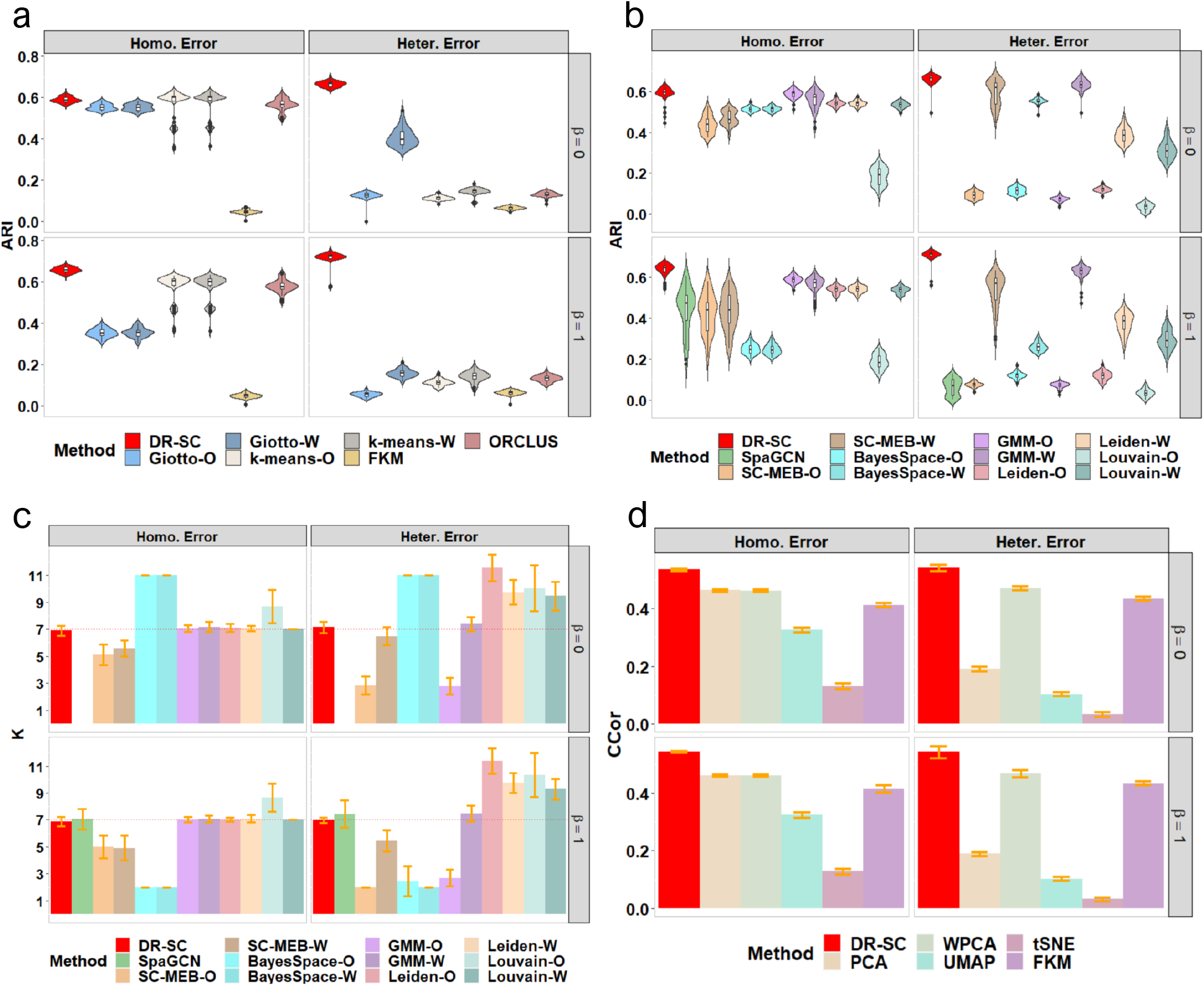
Comparisons of log-normalized gene expressions in Simulation 1. For tandem analysis, we used either PCA or WPCA to obtain PCs and named the analysis as method-O or method-W. a & b. Comparison of the clustering performance using the true number of clusters and with automatically selected cluster number, respectively. The clustering performance is evaluated using ARI. c. Comparison of the cluster-number selection performance of 12 methods that can choose the number of clusters. d. Comparison of the dimension-reduction performance of six methods using the average canonical correlation coefficients.

Next, we evaluated the performance of DR-SC in estimating the low-dimensional embeddings. For the average canonical correlation between the estimated embeddings, 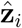, and the true latent features, **z**_*i*_, DR-SC had the highest canonical correlation coefficients (Fig. 2d and Supplementary Fig. S11a), suggesting that the estimated embeddings were more accurate. Pearson correlation coefficients between observed expressions, **x**_*i*_, and the estimated cell-type labels, *ŷ_i_*, conditioned on embeddings from DR-SC were much smaller than those from other methods (Supplementary Fig. S9c and Fig. S11a), suggesting that DR-SC captures more relevant information regarding cell types and, thus, facilitates the downstream analysis.

In addition, we evaluated the corresponding computational time for each method in all simulation settings, as shown in Supplementary Fig. S9a, S9b and Fig. S10 (bottom panel). Louvain and SpaGCN were computational the fastest, while BayesSpace was the slowest. Moreover, DR-SC was computationally efficient and scalable to large sample sizes, only taking around 15 mins to analyze a data with 1,000 genes and 100,000 spots (Supplementary Fig. S9d).

### 3.3 Human dorsolateral prefrontal cortex data

As an emerging spatial transcriptomics technology, the 10x Visium assay represents improvements in both resolution and the time needed to run the protocol [62]. Maynard et al. [14] used this technology to generate spatial maps of gene expression matrices for the six-layered DLPFC of the adult human brain and manually annotated Visium spots based on the cytoarchitecture. In this dataset, there were 12 tissue sections from three adult donors with a median depth of 291 million reads for each sample, a median of 3,844 spots per tissue section and a mean of 1,734 genes per spot. The raw gene expression count matrices were log-transformed and normalized using library size [56].

In this analysis, we considered both joint and tandem methods for dimension reduction and clustering. To apply the joint methods, we took the log-transformed raw count matrix using the library size as input, and in the tandem analysis, we obtained the top 15 PCs from either PCA or WPCA as input for the different clustering methods. As Giotto, *k*-means, FKM, and ORCLUS cannot choose the number of clusters, *K*, we fixed the number of clusters using manual annotations to make comparisons with DR-SC. When the cluster number, *K*, was fixed, the methods with spatial clustering, i.e., DR-SC and Giotto, outperformed those that did not consider spatial information, and DR-SC performed much better than Giotto (Fig. 3a). For all methods that were capable of selecting the number of clusters, we also observed that the spatial clustering methods, such as DR-SC, SpaGCN, SC-MEB, and BayesSpace, outperformed the non-spatial ones such as GMM, Leiden, and Louvain (Fig. 3b.). Note, there were only minor differences in the DR-SC when we used either the fixed *K* or the chosen *K*. We also evaluated the clustering performance using the NMI (Supplementary Fig. S12a) and similar patterns were observed. A heatmap of cell types from the manual annotations, and heatmaps of cluster assignments across spatial and non-spatial clustering methods for sample ID151510 are provided in Fig. 3c. The results for other 11 samples are provided in Supplementary Fig. S13a - S23a. In addition, Fig. 3d and Supplementary Fig. S13b - S23b show the tSNE plots for DR-SC and the other three dimension reduction methods (PCA, WPCA, UMAP), for which tSNE PCs were obtained from the estimated 15-dimensional features of each method with the class labels estimated in DR-SC. We observed better separation of tSNE PCs with DR-SC. Moreover, we evaluated the computational efficiency of DR-SC and compared it with that of other methods (Supplementary Fig. S12b) and found that DR-SC was about 10 times faster than FKM, ORCLUS, and BayesSpace.

**Figure 3:**
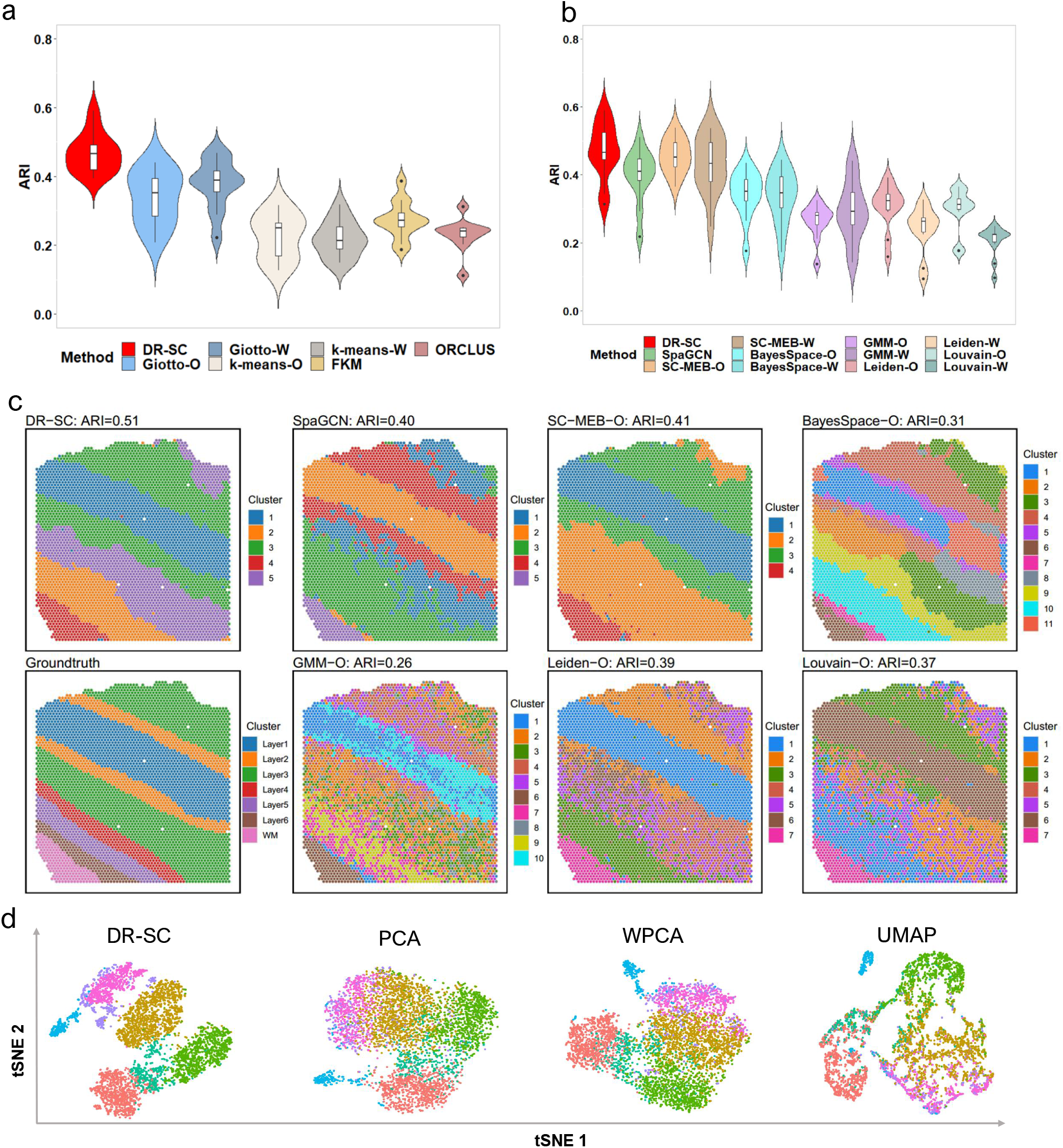
Analysis of human dorsolateral prefrontal cortex data. a. Violin plot of ARI values across 12 samples for DR-SC and other methods that cannot choose the number of clusters. The number of clusters in the analysis was fixed by manual annotations. b. Violin plot of ARI values across 12 samples for DR-SC and other methods that can choose tide number of clusters. c. Spatial heatmaps of cluster assignments for sample ID151510 using DR-SC and other spatial and non-spatial clustering methods. Left bottom corner denotes cell assignment from manual annotation; upper panel corresponds to the cell assignment from spatial clustering methods, and the rest of lower panel corresponds to the cell assignment from non-spatial clustering methods. d. Visualization of the cluster labels for sample ID151510 from DR-SC given the annotated number of clusters based on two-dimensional tSNE embeddings from four different DR methods including DR-SC, PCA, WPCA and UMAP.

We further performed conditional analysis to investigate the roles of SVGs beyond simple cell-type differences. Using SPARK [11], we performed spatial variation analysis (SVA) with the embeddings estimated by DR-SC as covariates. The detailed gene list identified at a false discovery rate (FDR) of 1% is given in Supplementary Table S6. Compared with the list of 1,583 SVGs identified by SVA without using covariates, the number of SVGs dramatically decreased to 113 at an FDR of 1% on average over 12 tissues after adjusting for covariates. Without adjusting for cell-type-relevant covariates, the gene expression variations identified by SVA could simply reflect cell-type differences. A Venn diagram of the links between SVGs obtained without adjusting for cell-type-relevant covariates and differentially expressed genes in different cell types (Supplementary Fig. S24a) showed that the majority of genes differentially expressed according to cell type were also identified as SVGs without adjusting for covaraites. Supplementary Fig. S24b and c show bar plots for the proportion of differentially expressed genes overlapping with SVGs without or with adjusting for covariates. The proportion of overlap was substantially reduced after we performed conditional spatial variation analysis, suggesting these genes were genuinely spatially expressed and not merely the result of variations between cell types.

Next, we performed functional enrichment analysis of SVGs adjusted for cell-type-relevant covariates. A total of 82 terms from Gene Ontology (GO), Kyoto Encyclopedia of Genes and Genomes (KEGG), and Human Protein Atlas (HPA) were enriched with adjusted *p*-values of less than 0.05 in at least three DLPFC tissue sections. Supplementary Fig. S25 shows the top five pathways among all 12 DLPFC tissue sections, in which many common terms could be identified after controlling for cell-type-relevant covariates. These results suggested that SVGs adjusted for cell-type-relevant covariates shared common spatial patterns in the brain tissue. For example, the same set of highly significant HPA terms were identified in 8 out of 12 tissue samples, including processes in white matter, processes in granular layer, and cytoplasm/membrane (Supplementary Fig. S25). Nearly all the most significant KEGG pathways were identified in all 12 samples, including Huntington’s disease, Alzheimer’s disease, and Parkinson’s disease, etc. Several studies [63, 64] reported the distribution of abnormal proteins across the brain causes damage in Alzheimer’s disease, Parkinson’s disease, Huntington’s disease, and other neurodegenerative diseases. Additionally, many common significant GO terms were identified in all 12 samples, such as electron transfer activity, structural molecule activity, structural constituents of the cytoskeleton, oxidative phosphorylation, endocytic vesicle lumen, and respiratory chain complex. Details of the top five pathways for all 12 tissue samples are presented in Supplementary Table S7.

### 3.4 Mouse olfactory bulb data

Slide-seq is a spatial transcriptomics technology that simultaneously decouples imaging from molecular sampling and quantifies expression across the genome with 10-*μm* spatial resolution [10]. To further improve the sensitivity magnitude and enable the more efficient recovery of gene expression, Slide-seqV2 was recently introduced [65] to generate two datasets each from mouse olfactory bulb and mouse cortex. We present our analysis of the mouse olfactory bulb dataset in this section and that of the mouse cortex dataset in the next one. The olfactory bulb contained 21,041 spots and 37,329 genes with a median of 494 unique molecular identifiers per bead. The raw gene expression count matrices were log-transformed and normalized using library size.

In the analysis, we applied the spatial clustering methods BayesSpace, SC-MEB, SpaGCN and DR-SC, all of which, except for DR-SC, were based on tandem analysis. Thus, BayesSpace and SC-MEB took the top 15 PCs from the normalized expression matrix of SVGs as input (see Materials and Methods), while the SpaGCN package took the normalized expression matrix as input and used its internally embedded PCA algorithm to obtain the PCs. In a joint method, DR-SC used the normalized expression matrix of SVGs as input. A spatial heatmap of the cluster assignments across the four methods is provided in Fig. 4a, while the tSNE plots for these four methods is shown in Fig. 4b, where the tSNE PCs of DR-SC were obtained from its estimated 15-dimensional latent features while tSNE PCs of BayesSpace-O, SC-MEB-O, and SpaGCN were based on 15 PCs using the ordinary PCA, denoted as suffix -O. We obtained a clearer visualization of the cell types using tSNE PCs from DR-SC. We also compared the running time of these methods (Supplementary Fig. S26a) and found DR-SC and BayesSpace respectively took 1,042 and 13,193 secs to complete the analysis for all 21,041 spots. To further compare the visualizations of the different dimension-reduction methods, we first applied the other eight dimension-reduction methods to extract latent features and obtained two-dimensional tSNE PCs based on each estimated latent one. We visualized the twodimensional tSNE PCs derived from the different dimension-reduction methods with cluster labels estimated in DR-SC (Fig. 4c and Supplementary Fig. S26b), and the tSNE PCs from DR-SC were more distinguishable than those from other methods.

**Figure 4:**
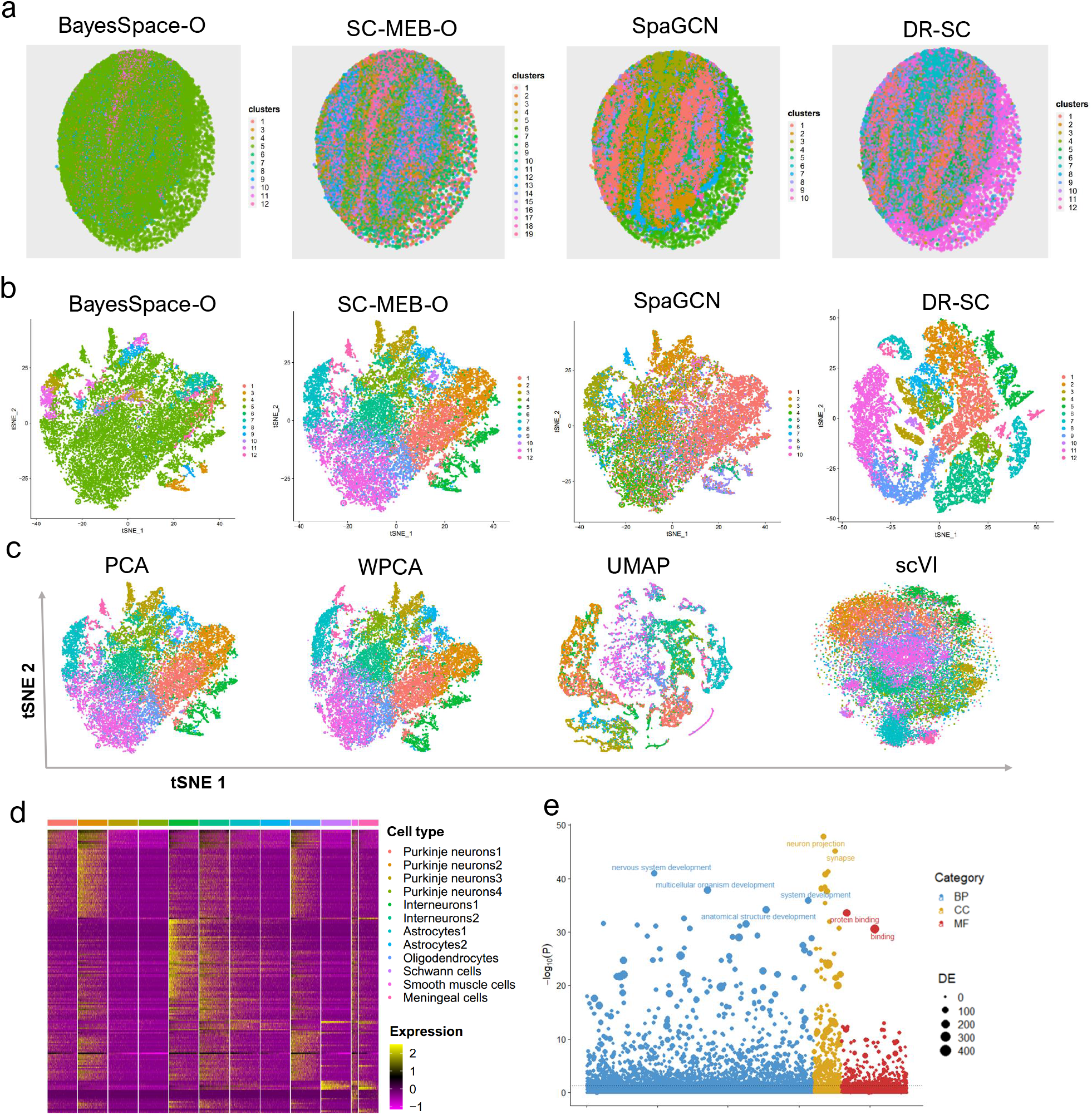
Analysis of mouse olfactory bulb data. a. Spatial heatmap of clusters from four spatial-clustering methods. b. tSNE plots for these four methods, where tSNE PCs of DR-SC were obtained based on the extracted 15-dimensional features, while tSNE PCs of BayesSpace-O, SC-MEB-O and SpaGCN were based on 15 PCs from PCA. c. Visualization of the cluster labels from DR-SC based on two-dimensional tSNE embeddings from four different DR methods, PCA, WPCA, UMAP, and scVI. d. Heatmap of differentially expressed genes for each cell type identified by DR-SC. e. Bubble plot of –log10(*p*–values) for pathway enrichment analysis of 518 SVGs with adjusted *p*-values of less than 0.05. Dashed line represents a *p*-value cutoff of 0.05. Gene sets are colored by category: GO biological process (BP, blue), and GO cellular component (CC, yellow), GO molecular function (MF, brown).

Using the cluster labels estimated in DR-SC, we performed DGE analysis to identify the marker genes for each cluster. The heatmap of differentially expressed genes for each cell type (Fig. 4d) showed good separation across the different cell clusters. By checking PanglaoDB [54] for the identified marker genes, we were able to identify seven cell types in the heatmap (Fig. 4d), including two major neuron cell types: Purkinje neurons and interneurons consisting of 52% and 10% spots, respectively. The primary output signal for Purkinje cells was the modulated discharge of simple spikes while interneurons potentially contributed to the modulation of simple spikes [66].

We then performed spatial variational analysis using SPARK by controlling for the 15-dimensional embeddings estimated by DR-SC. In total, 518 SVGs were identified at an FDR of 1%, and the identified genes are listed in Supplementary Table S8. Next, we performed functional enrichment analysis of these SVGs and 385 GO terms were found to be enriched with adjusted *p*-values of less than 0.05. A bubble plot for this functional enrichment (Fig. 4e) showed that the nervous-system-development-related pathways were enriched in the olfactory bulb.

We additionally applied Slingshot [36] to perform cell lineage analysis using low-dimensional embeddings and the cluster labels estimated by DR-SC. Srivatsan et al. [67] reported that the neuron cells differentiate after glia cells. To check this, we focused on studying neuron cells (Purkinje neurons and interneurons) and glia cells (astrocytes and oligodendrocytes) to infer their differentiation trajectory. The inferred trajectory shown in Supplementary Fig. S26c led us to conclude that Purkinje neurons differentiated after oligodendrocytes, while interneurons differentiated after astrocytes. Supplementary Fig. S26c also provides a heatmap of the expression levels of the top 20 most significant genes presenting dynamic expression patterns over pseudotime. In this analysis, some genes presented interesting dynamic expression patterns, varying from high to low levels and back to high levels, such as *Camk2b* and *Malat1.*

### 3.5 Mouse E15 neocortex data

The mouse E15 neocortex data from the Slide-seqV2 platform contains 33,611 spots and 22,683 genes resolved spatially according to their expression in E15 embryo section. Similarly, we performed spatial clustering by DR-SC and compared it with the BayesSpace, SC-MEB, and SpaGCN results. For BayesSpace and SC-MEB, the top 15 PCs from the normalized expression matrix of SVGs were used as input (see Materials and Methods), while for SpaGCN and DR-SC, the normalized expression matrix was the input. SC-MEB-O and DR-SC shared similar spatial patterns, whereas BayesSpace-O assigned a large proportion of spots (78%) to a single cluster (Fig. 5a). We also gained a better visualization of clusters using tSNE PCs from DR-SC compared to PCA (Fig. 5b). We also compared the running time of these methods (Supplementary Fig. S27a), and DR-SC and BayesSpace respectively took 1,218 and 18,740 secs to analyze all 33,611 spots. To further compare the visualizations obtained via the different dimension-reduction methods, we first applied the other eight dimension reduction methods to extract latent features and obtained two-dimensional tSNE PCs based on each estimated latent one. We visualized the two-dimensional tSNE PCs from the different dimension reductionmethods with cluster labels estimated in DR-SC (Fig 5c and Supplementary Fig. S27b), which indicated DR-SC provided the best visualization.

**Figure 5:**
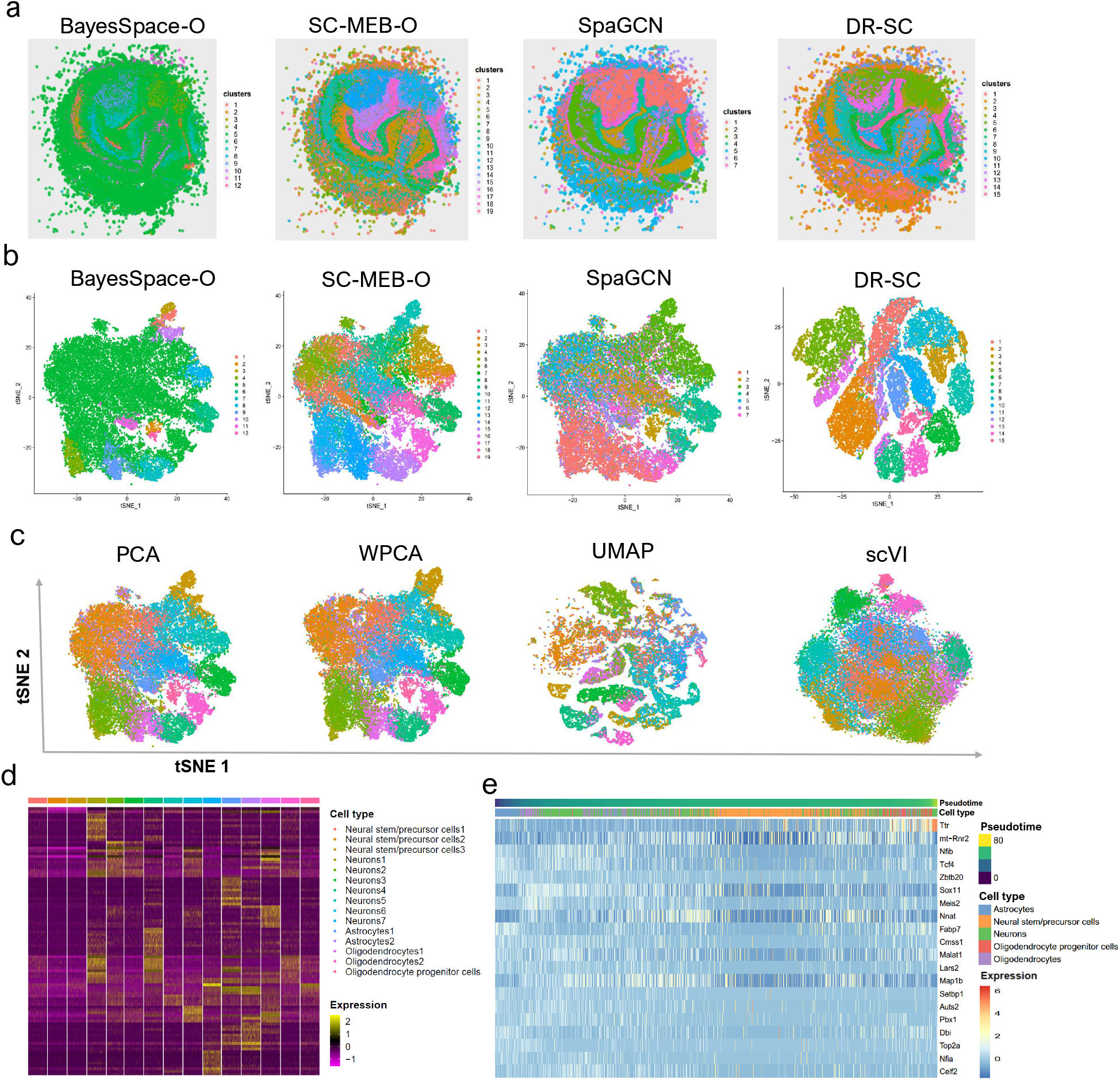
Analysis of mouse E15 neocortex data. a. Spatial heatmap of clusters from four spatial-clustering methods: BayesSpace-O, SC-MEB-O, SpaGCN, and DR-SC. b. tSNE plots for these four methods, where tSNE PCs of DR-SC were obtained based on the extracted 15-dimensional features while tSNE PCs of BayesSpace-O, SC-MEB-O, and SpaGCN were based on 15 PCs from PCA. c. Visualization of the cluster labels estimated by DR-SC based on two-dimensional tSNE embeddings from four different DR methods: PCA, WPCA, UMAP, and scVI. d. Heatmap of differentially expressed genes for each cell type identified by DR-SC. e. Heatmap of gene expression levels of the top 20 genes with significant changes with respect to the Slingshot pseudotime. Each column represents a spot that is mapped to this path and is ordered by its pseudotime value. Each row denotes the most significantly changed gene expression.

Based on the cluster labels estimated in DR-SC, we performed DGE analysis to identify marker genes for each cluster. A heatmap of the findings showed good separation of the differentially expressed genes across different cell types (Fig. 5d). By checking PanglaoDB [54] for the identified marker genes, we were able to identify five cell types, including two major neuron-related cell types: neurons and neural stem/precursor cells consisting of 40% and 29% spots, respectively.

Next, we applied Slingshot [36] to infer the differentiation lineages of E15 neocortex cells based on the low-dimensional features and cluster labels estimated by DR-SC. The inferred development trajectory of different types of cells and a heatmap of the top 20 significant dynamic expressed genes along the trajectory were plotted (Fig. 5e). We also observed the differentiation of neuron cells (neurons and neural stem/precursor cells) after glia cells (astrocytes, oligodendrocytes and oligodendrocyte progenitor cells). For example, a portion of the neurons and all neural stem/precursor cells differentiated after the astrocytes, and the remainder of the neurons differentiated after oligodendrocytes and oligodendrocyte progenitor cells. According to the heatmap, some genes presented interesting dynamic patterns of expression. For instance, the gene *Ttr* had low expression levels at the early stage before substantially increasing later. In contrast, the expression levels of *Nfib, Sox11, Nnat* and *Map1b* changed from low to high, then back to low. Steele-Perkins et al. [68] reported that the transcription factor gene *Nfib* is essential for mouse brain development. Jankowski et al. [69] found that expression of the transcription factor gene *Sox11* modulates peripheral nerve regeneration in mice. *Nnat* was reported the spatial expression pattern during mouse eye development [70]. *Map1b* is required for axon guidance and is involved in the development of the central and peripheral nervous systems [71].

### 3.6 Mouse embryo data

We applied DR-SC to analyze a large seqFISH dataset of mouse organogenesis [16] that contained 23,194 cells. In this dataset, the expression of a panel of 351 genes was resolved spatially within multiple 8- to 12-somite-stage mouse embryo sections using the seqFISH platform. Cell labels were accurately annotated across the embryo [16] based on their nearest neighbors within an existing scRNA-seq atlas (Gastrulation atlas) [72].

We first performed clustering analysis using DR-SC and other existing spatial-clustering methods, SpaGCN, BayesSpace, SC-MEB and Giotto. Taking the above manually annotated cell types as reference, we compared the clustering performance of DR-SC with that of the other methods. As the other methods involved tandem analyses, we obtained the top 15 PCs [26] using either PCA or WPCA from all 351 genes for use as input in these, except for SpaGCN. DR-SC provided better clustering performance than the other clustering methods in terms of the ARI values (Supplementary Fig. S28a). A heatmap of cell types according to the mannual annotations is provided in Supplementary Fig. S28b, while heatmaps of the cell types inferred by DR-SC, SC-MEB, and BayesSpace are provided in Supplementary Fig. S28c. The cell labels estimated by DR-SC and SC-MEB, but not BayesSpace, were in agreement with those in the manual annotations. BayesSpace incorrectly clustered many of the cells as ”low quality” cells.

To refine the analysis of the brain regions, we first collected cells manually annotated as “forebrain, midbrain, or hindbrain”, then we applied DR-SC to estimate low-dimensional embeddings and provide labels for cells in the three brain regions (Fig. 6a). DR-SC identified a total of six clusters. By checking PanglaoDB [54] for the marker genes identified via DGE analysis, we were able to identify four cell types (astrocytes, microglia cells, neurons 1/2, and ependymal cells 1/2; Fig. 6b) and four cortical regions (forebrain, hindbrain 1/2/3, midbrain, and microglia; Fig. 6c). Details of the cell typing are provided in Supplementary Table S9. Note that neuron cells were found in both forebrain and hindbrain regions, while glia (astrocytes, microglia) cells were from both the midbrain and microglia regions. A recent study [67] reported that neurons and glia cells can be distributed over different brain regions. The tSNE plot for the regions and cell types in Fig. 6d shows that DR-SC effectively separated the different clusters.

**Figure 6:**
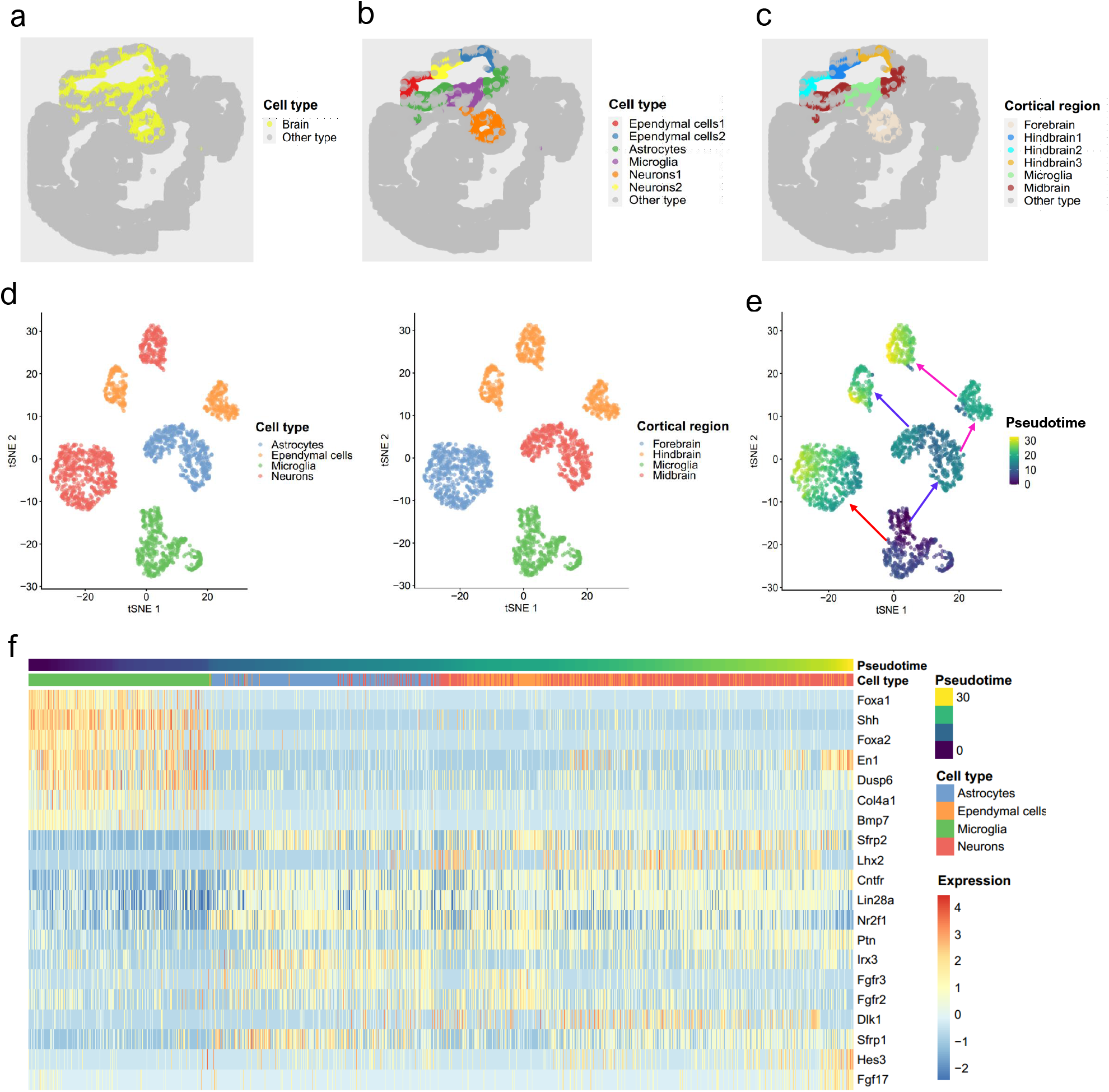
Analysis of mouse embryo data. a. Spatial heatmap of brain regions and other areas. b. Spatial heatmap of cell types based on clusters identified by DR-SC. c. Spatial heatmap of cortical region identified by DR-SC. d. tSNE plot of cell types and corresponding cortical regions, where the tSNE projection was evaluated based on estimated low-dimensional embeddings using DR-SC. Note, the cortical regions and cell types are well separated. e. tSNE plot of inferred pseudotime using Slingshot based on the estimated low-dimensional embeddings and cluster labels for the cortical region from DR-SC. f. Heatmap of gene expression levels for the top 20 genes with significant changes with respect to the Slingshot pseudotime. Each column represents a spot that is mapped to this path and is ordered by its pseudotime value. Each row denotes the most significantly changed gene expression.

To further investigate the development and differentiation of these brain cells, we calculated the pseudotime using Slingshot based on the 15-dimensional embeddings and cluster labels estimated using DR-SC. We identified three lineages that were consistent with the findings in a previous study [67]. A plot of the inferred lineages with pseudotime (Fig. 6e) provided an ilustration of the dynamic trajectory from glia cells to neurons. Following Srivatsan et al. [67], we used Allen Brain Reference Atlases (http://atlas.brain-map.org/) as guides to check how these trajectories were distributed over brain segments. The cells and trajectories from each cluster overwhelmingly occupied different brain regions (Fig. 6d and e). While combing pseudotime and region spatial information, we observed that cells in the early differentiation stage were clustered in the microglia and midbrain regions. Later, cells with differentiated transcriptomes emerged in more distant regions, i.e., the hindbrain and forebrain. According to the inferred pseudotime, we identified differentially expressed genes along cell pseudotime using the method described by Ji et al. [34]. The heatmap of the expression of the top 20 most significant genes (Fig. 6f) suggested the occurance of some interesting dynamic expression patterns over pseudotime. The genes *Foxa1, Shh,* and *Foxa2* had higher expression levels at the early stage, but later their expression levels decreased substantially. In contrast, the expression levels of the genes *Fgfr2* and *Fgfr3* changed from low to high and then returned to low. We observed the pattern of expression of *En1* changed from high to low then to high along the inferred trajectory. In humans, the *En1* gene codes for the homeobox protein engrailed (EN) family of transcription factors. A recent study [73] reported that *En1* shows a transcriptional dependency in triple negative breast cancer associated with brain metastasis. Carratala-Marco et al. [74] found that EN plays an important role in the regionalization of the neural tube and EN’s distribution regulates cerebellum and midbrain morphogenesis, as well as retinotectal synaptogenesis.

## 4 Discussion

In this paper, we have proposed a joint DR-SC for analyzing high-dimensional scRNA-seq and spatial transcriptomics data using a hierarchical model. In contrast to most existing methods that perform dimensional reduction and (spatial) clustering sequentially, DR-SC unifies lowdimensional feature extraction with (spatial) clustering in the same, joint modeling framework, and provides an improved estimation for cell-type-relevant low-dimensional embeddings and enhanced clustering performance for both scRNA-seq and spatial transcriptomic data from different platforms. With simulation studies and benchmark dataset analyses, we demonstrated that DR-SC can improve clustering performance while effectively estimating low-dimensional embeddings.

DR-SC relies on a hidden Markov random field model with a smoothing parameter to perform spatial clustering. The probabilistic framework of DR-SC allows us to adaptively update the spatial smoothing parameter that promotes similar cluster assignments for neighboring tissue locations in a data-driven manner. When the smoothness parameter is set to zero, DR-SC performs clustering for scRNA-seq data without spatial information. We developed an efficient EM algorithm based on iterative conditional mode and expectation-maximization (ICM-EM), making DR-SC computationally efficient and scalable to large sample sizes.

In-depth analyses using scRNA-seq and spatial transcriptomic data from different platforms showed that the estimated clusters and embeddings from DR-SC effectively facilitated the downstream analysis. First, we applied DR-SC to a 10x Visium dataset for DLPFC to demonstrate the improved spatial clustering performance of DR-SC and further carried out conditional SVA to identify genes with pure spatial variations but not cell-type differences. The majority of genes identified in SVA without adjustment for cell-type-relevant covariates simply reflected cell-type differences. Functional enrichment analysis showed that the genes identified in SVA with adjustment for covariates were enriched in pathways related to the DLPFC tissue. For example, the most significant KEGG pathways including Huntington’s disease and Alzheimer’s disease were identified in all 12 LIBD samples. Second, we applied DR-SC to analyze two Slide-seqV2 datasets and found it outperformed both existing dimension reduction methods in terms of visualization and existing spatial clustering methods in terms of separation, as well as its usefulness in cell trajectory inference. In the mouse olfactory bulb data, we identified some genes with interesting dynamic expression patterns, such as *Camk2b* and *Malat1.* Küry et al. [75] reported that *Camk2b* was important for learning and synaptic plasticity in mice, while Zhang et al. [76] reported a potential cis-regulatory role of *Malat1* gene transcription in mice. Third, we applied DR-SC to analyze a seqFISH dataset and showcased its ability to infer cell lineages based on the reduced-dimensionality space estimated by DR-SC. Among the identified genes with interesting dynamic patterns over pseudotime, transcription factors *Foxa1* and *Foxa2* are crucial in maintaining key cellular and functional features of dopaminergic neurons in the brain [77] while *Fgfr2* and *Fgfr3* play important roles during early neural development [78, 79]. Fourth, we demonstrated both the higher Kendall’s and Spearman’s rank correlation coefficients between the true and inferred pseudotime using DR-SC among 16 benchmark scRNA-seq datasets (Supplementary Text). Finally, using a CITE-seq dataset for CBMC, we demonstrated that analysis using DR-SC can improve clustering performance while facilitating the identification of differentially expressed genes among the different cell types in the analysis of scRNA-seq data (Supplementary Text).

There are several potential extensions that can be applied to DR-SC. First, in the current study, we considered single transcriptional profiles. The framework of DR-SC could be naturally extended to perform joint-clustering analyses of multiple samples by properly removing their batch effects. Second, the fast-evolving technology of single-cell omics provides opportunities to integrate omics profiles from different modalities for the same individuals. Extending DR-SC by integrating multiple different omics techniques, such as through the canonical correlation analysis framework [80] which is a nature extension of PCA towards multiple modality analysis, will also likely achieve higher statistical performance. Third, DR-SC essentially performs unsupervised clustering, but with the availability of labels for some cells/spots, it would be interesting to perform semi-supervised clustering of those data. We will investigate these issues in future work.

## Availability of data and materials

All codes in this paper are publicly available at https://github.com/feiyoung/DR-SC.Analysis. The source code is released under the GNU general public license. The 16 benchmark datasets with linear trajectory information are available at https://zenodo.org/record/1443566#.XNV25Y5KhaR. The cord blood mononuclear cells datasets are available at https://www.ncbi.nlm.nih.gov/geo/query/acc.cgi?acc=GSE100866 via the accession number GSE100866. The human dorsolateral prefrontal cortex datasets on the 10x Visium platform are accessible at https://github.com/LieberInstitute/spatialLIBD. The mouse olfactory bulb data and mouse E15 neocortex data on the Slide-seqV2 platform are available at https://singlecell.broadinstitute.org/single_cell/data/public/SCP815. The mouse embryo dataset on the seqFISH platform is accessible at https://content.cruk.cam.ac.uk/jmlab/SpatialMouseAtlas2020/.

## Authors’ contributions

J.L., X.S., and X.Z. initiated and designed the study, W.L., X.L., and Y.Y. implemented the model and performed simulation studies and benchmarking evaluation; J.L., X.S., and X.Z. wrote the manuscript, and all authors edited and revised the manuscript.

## Acknowledgements

This work was supported by grant AcRF Tier 2 (MOE2018-T2-2-006 and T2EP20220-0021) from the Ministry of Education, Singapore, grants from the National Natural Science Foundation of China (NSF: 11931014 and NSF: 12171229), and grant 22ZR1420500 from the Shanghai Committee of Science and Technology. The computational work for this article was partially performed using resources from the National Supercomputing Centre, Singapore (https://www.nscc.sg).

