## Supplementary Figure for "Joint dimension reduction and clustering analysis for single-cell RNA-seq and spatial transcriptomics data"

#### Contents

**1 Supplementary Figures in Supplementary Text** **3**

1.1 Results in Lineage Analysis . . . . . . . . . . . . . . . . . . . . . 3

1.2 Results for Cord blood mononuclear cells data . . . . . . . . . . . . . . . . . . . 6

**2 Supplementary Figures in Main Text** 11

### 1 Supplementary Figures in Supplementary Text

#### 1.1 Results in Lineage Analysis

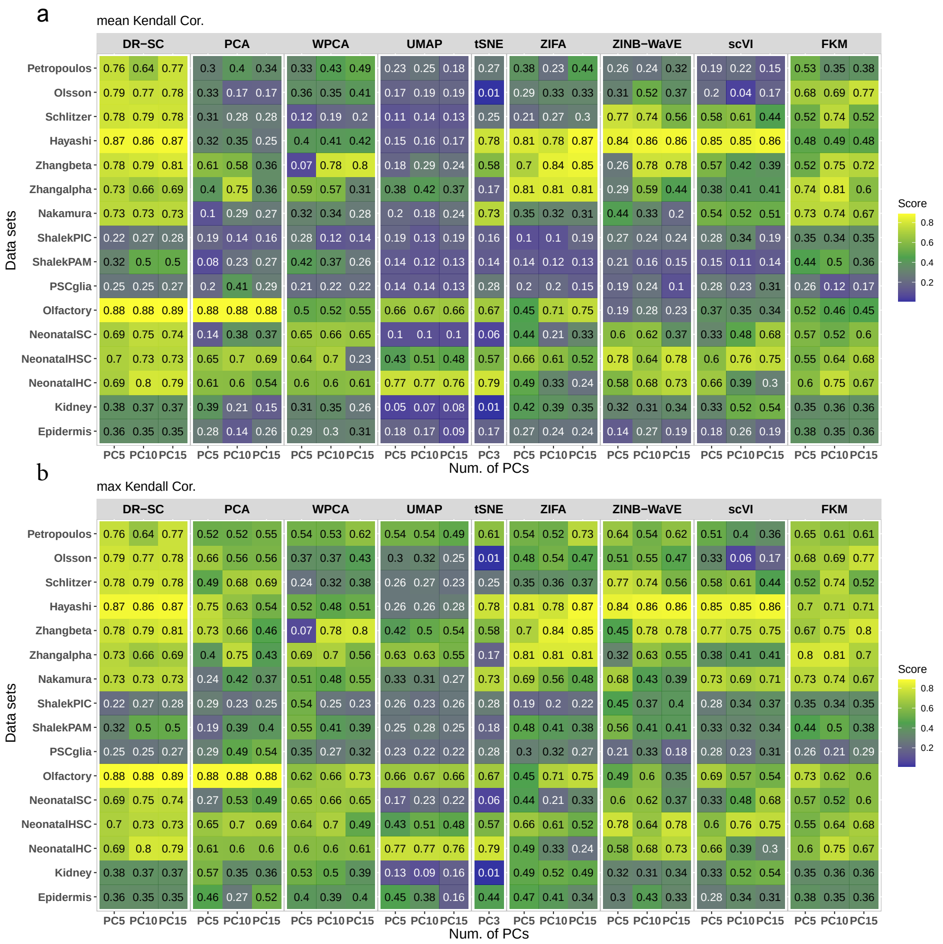

Figure S1: Dimension-reduction performance of DR-SC and other methods in downstream trajectory inference using Slingshot for 16 benchmarking datasets. We compared DR-SC with eight dimension reduction methods (columns), including principal component analysis (PCA), weighted principal component analysis (WPCA), uniform manifold approximation and projection (UMAP),t-distributed stochastic neighbor embedding (tSNE), zero-inflated factor analysis (ZIFA), zero-inflated negative binomial-based wanted variation extraction (ZINB-WaVE), deep-learning-based scVI and factorial *k*-means (FKM). We evaluated their performance on 16 real scRNA-seq data sets (rows) in terms of lineage inference accuracy using mean/maximum Kendall correlation coefficients (a/b). We used Slingshot for DR-SC and FKM with their own clusters but for the other dimension-reduction methods with clusters from the Gaussian mixture model as the initial step for lineage inference. The performance of each dimension-reduction method was measured by mean/maximum Kendall correlation between the true lineage and inferred lineage based on different methods. For each dataset, we compared three different numbers of low-dimensional components (5, 10, and 15). Note that, for tSNE, we only extracted three low-dimensional components due to the limitation of the tSNE software.

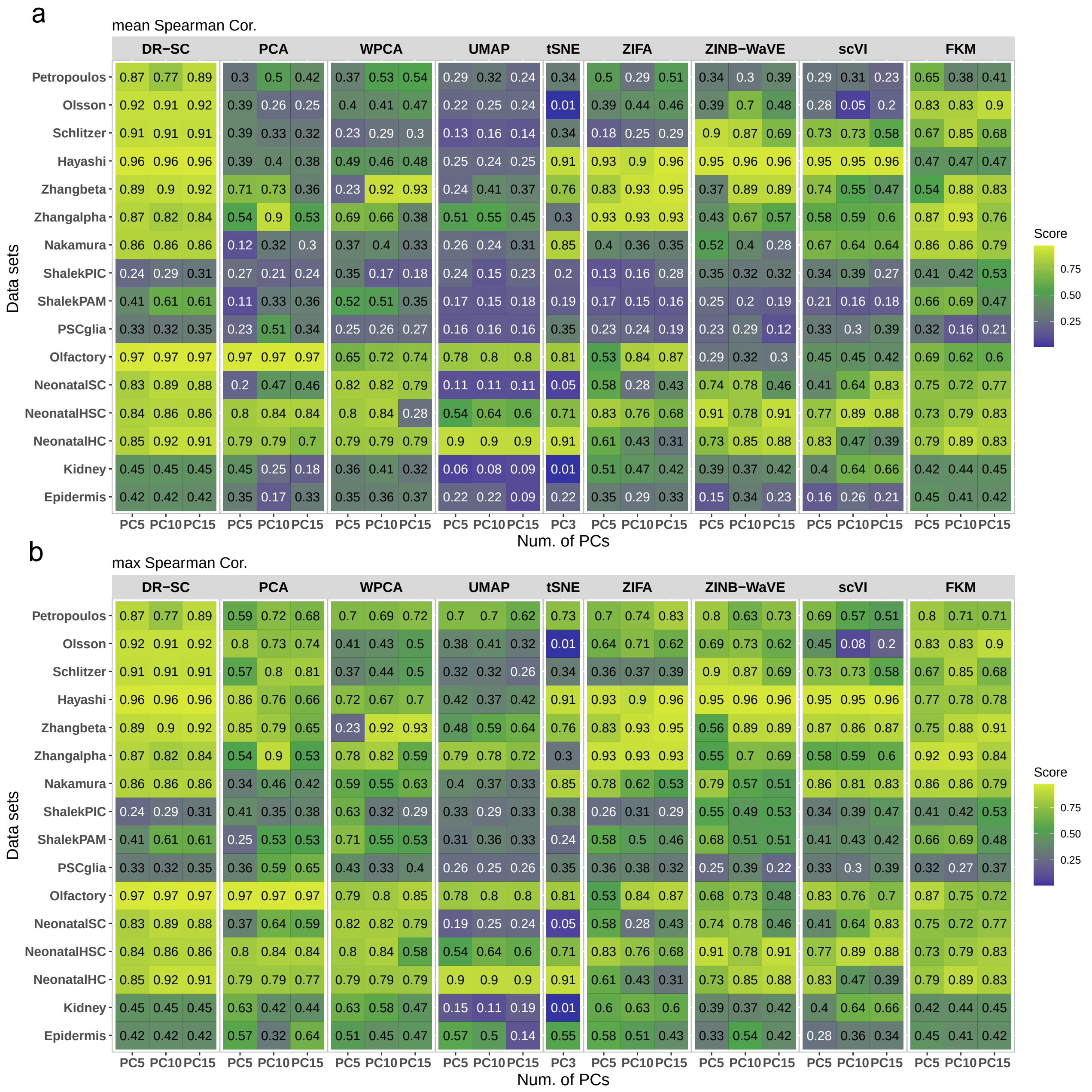

Figure S2: Dimension-reduction method performance evaluated by mean/maximum Spearman correlation in the trajectory inference analysis. We compared DR-SC with eight dimension-reduction methods (columns), including principal component analysis (PCA), weighted principal component analysis (WPCA), uniform manifold approximation and projection (UMAP), t-distributed stochastic neighbor embedding (tSNE), zero-inflated factor analysis (ZIFA), zero-inflated negative binomial-based wanted variation extraction (ZINB-WaVE), deep-learning-based scVI, and factorial $k$-means (FKM). We evaluated their performance on 16 real scRNA-seq data sets (rows) in terms of lineage inference accuracy. We used Slingshot for DR-SC and FKM with their own clusters but for the other dimension reduction methods with clusters from GMM as the initial step for lineage inference. The performance of each dimensionality reduction method is measured by mean/maximum Spearman correlation (a/b). For each dataset, we compared three different numbers of low-dimensional components (5, 10, and 15; three sub-columns under each column). Note that, for tSNE, we only extracted three low-dimensional components due to the limitation of the tSNE software.

Num. of lineages

| Data sets | DR−SC | | | PCA | | | WPCA | | | UMAP | | | tSNE | ZIFA | | | ZINB−WaVE | | | scVI | | | FKM | | |
|---|---|---|---|---|---|---|---|---|---|---|---|---|---|---|---|---|---|---|---|---|---|---|---|---|---|
| Num. of PCs | PC5 | PC10 | PC15 | PC5 | PC10 | PC15 | PC5 | PC10 | PC15 | PC5 | PC10 | PC15 | PC3 | PC5 | PC10 | PC15 | PC5 | PC10 | PC15 | PC5 | PC10 | PC15 | PC5 | PC10 | PC15 |
| Petropoulos | 1 | 1 | 1 | 3 | 4 | 6 | 5 | 5 | 5 | 7 | 7 | 7 | 6 | 3 | 4 | 3 | 5 | 8 | 7 | 9 | 8 | 9 | 2 | 2 | 2 |
| Olsson | 1 | 1 | 1 | 4 | 7 | 7 | 2 | 2 | 2 | 3 | 3 | 3 | 1 | 5 | 6 | 7 | 3 | 3 | 3 | 2 | 2 | 2 | 1 | 1 | 1 |
| Schlitzer | 1 | 1 | 1 | 3 | 4 | 4 | 2 | 2 | 2 | 5 | 5 | 8 | 1 | 2 | 2 | 2 | 1 | 1 | 1 | 1 | 1 | 1 | 1 | 1 | 1 |
| Hayashi | 1 | 1 | 1 | 4 | 2 | 3 | 3 | 3 | 3 | 3 | 3 | 3 | 1 | 1 | 1 | 1 | 1 | 1 | 1 | 1 | 1 | 1 | 2 | 2 | 2 |
| Zhangbeta | 1 | 1 | 1 | 3 | 5 | 4 | 1 | 1 | 1 | 5 | 4 | 3 | 1 | 1 | 1 | 1 | 3 | 1 | 1 | 2 | 2 | 2 | 3 | 1 | 2 |
| Zhangalpha | 1 | 1 | 1 | 1 | 1 | 2 | 3 | 4 | 3 | 2 | 2 | 3 | 1 | 1 | 1 | 1 | 3 | 2 | 3 | 1 | 1 | 1 | 2 | 1 | 2 |
| Nakamura | 1 | 1 | 1 | 7 | 8 | 8 | 4 | 4 | 4 | 3 | 3 | 3 | 1 | 4 | 6 | 8 | 2 | 2 | 2 | 2 | 2 | 2 | 1 | 1 | 1 |
| ShalekPIC | 1 | 1 | 1 | 3 | 3 | 3 | 3 | 3 | 3 | 2 | 2 | 2 | 2 | 2 | 2 | 2 | 3 | 3 | 3 | 1 | 1 | 2 | 1 | 1 | 1 |
| ShalekPAM | 1 | 1 | 1 | 3 | 2 | 3 | 3 | 3 | 3 | 6 | 4 | 4 | 3 | 7 | 8 | 8 | 8 | 8 | 8 | 5 | 5 | 5 | 1 | 1 | 2 |
| PSCglia | 1 | 1 | 1 | 3 | 2 | 3 | 3 | 3 | 3 | 6 | 5 | 5 | 1 | 4 | 6 | 6 | 4 | 3 | 3 | 1 | 1 | 1 | 1 | 2 | 2 |
| Olfactory | 1 | 1 | 1 | 1 | 1 | 1 | 2 | 2 | 2 | 1 | 1 | 1 | 1 | 1 | 1 | 1 | 4 | 4 | 5 | 3 | 3 | 3 | 2 | 2 | 2 |
| NeonatalSC | 1 | 1 | 1 | 2 | 3 | 3 | 1 | 1 | 1 | 8 | 9 | 10 | 1 | 1 | 1 | 1 | 1 | 1 | 1 | 1 | 1 | 1 | 1 | 1 | 1 |
| NeonatalHSC | 1 | 1 | 1 | 1 | 1 | 1 | 1 | 1 | 3 | 1 | 1 | 1 | 1 | 1 | 1 | 1 | 1 | 1 | 1 | 1 | 1 | 1 | 1 | 1 | 1 |
| NeonatalHC | 1 | 1 | 1 | 1 | 1 | 2 | 1 | 1 | 1 | 1 | 1 | 1 | 1 | 1 | 1 | 1 | 1 | 1 | 1 | 1 | 1 | 1 | 1 | 1 | 1 |
| Kidney | 1 | 1 | 1 | 3 | 3 | 3 | 4 | 3 | 3 | 3 | 4 | 3 | 1 | 2 | 2 | 2 | 1 | 1 | 1 | 1 | 1 | 1 | 1 | 1 | 1 |
| Epidermis | 1 | 1 | 1 | 4 | 3 | 4 | 6 | 5 | 4 | 4 | 4 | 3 | 4 | 4 | 5 | 4 | 4 | 4 | 5 | 3 | 3 | 5 | 1 | 1 | 1 |

Figure S3: Estimated number of lineages for 16 benchmark datasets with linear linage. We compared DR-SC with eight dimension-reduction methods (columns), including principal component analysis (PCA), weighted principal component analysis (WPCA), uniform manifold approximation and projection (UMAP), t-distributed stochastic neighbor embedding (tSNE), zero-inflated factor analysis (ZIFA), zero-inflated negative binomial-based wanted variation extraction (ZINB-WaVE), deep-learning-based scVI and factorial $k$-means (FKM). We evaluated their performance on 16 real scRNA-seq data sets (rows) in terms of estimation of the number of lineages. We used Slingshot for DR-SC and FKM with their own clusters but for the other dimension-reduction methods with clusters from GMM as the initial step for lineage inference. For each dataset, we compared three different numbers of low-dimensional components (5, 10, and 15; three sub-columns under each column). Note, for tSNE, we only extracted three low-dimensional components due to the limitation of the tSNE software.

#### 1.2 Results for Cord blood mononuclear cells data

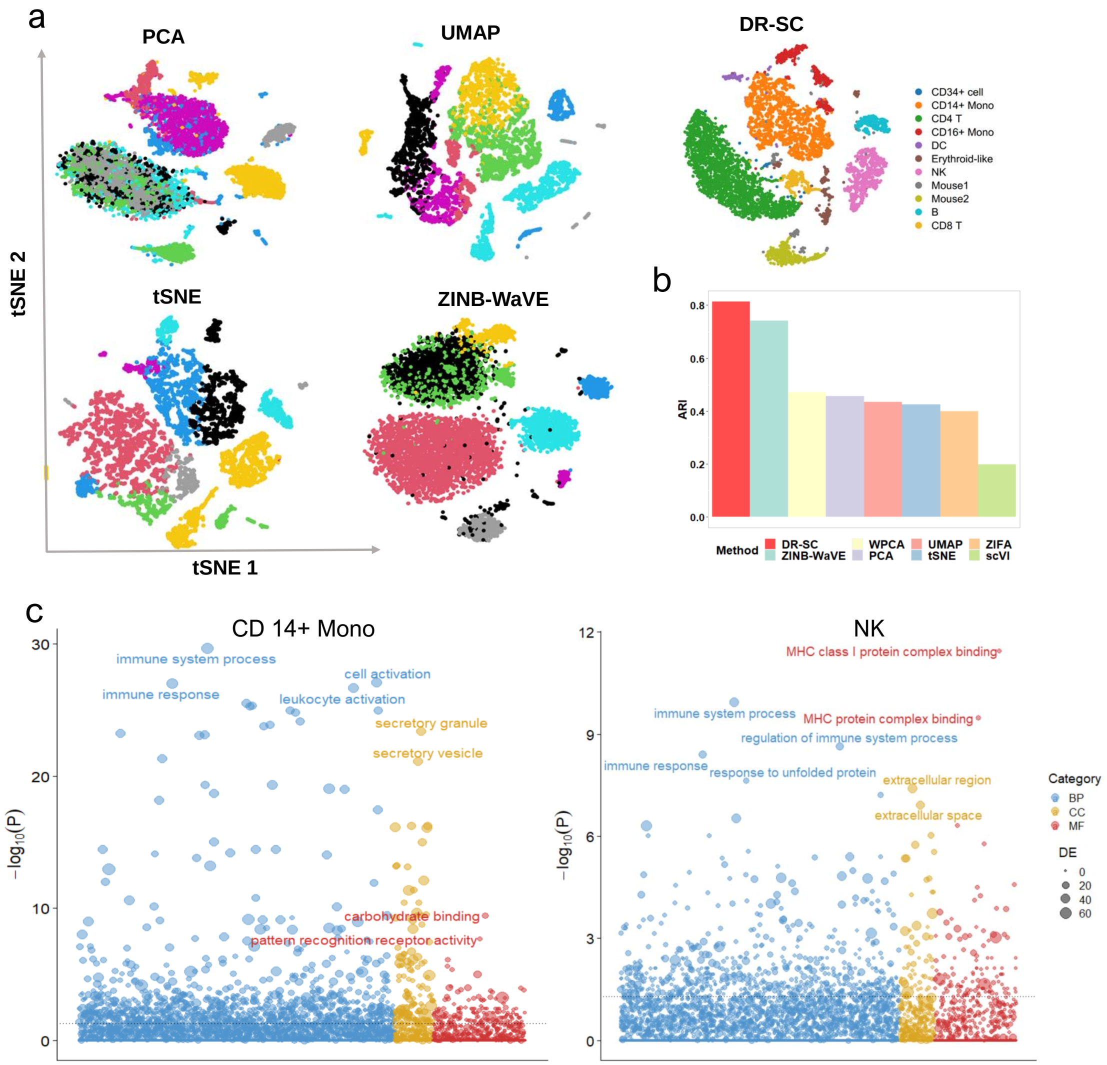

Figure S4: Analysis of human cord blood mononuclear cell data. a. tSNE plots of two-dimensional representations for visualization of cell-type clustering based on different dimension-reduction methods, DR-SC, PCA, UMAP, tSNE, and ZINB-WaVE. First, we obtained 25-dimensional embeddings by applying all methods, except for tSNE, then applied tSNE to obtain two-dimensional embeddings. DR-SC estimated low-dimensional embeddings and class labels simultaneously while further clustering analysis using GMM was required for all other dimension-reduction methods. b. Barplot of ARI values for DR-SC and GMM for the other seven dimension-reduction methods. c. Bubble plot of $-\log 10(p-$values) for pathway enrichment analysis of 171 and 102 DE genes of CD14+ Mono and natural killer (NK) cells, respectively, obtained by applying beta-poisson model for single-cell RNA-seq data (BPSC) with two cutoffs: 0.05 for the adjusted $p$-value, and 0.5 for the log fold-change. Dashed line represents a $p$-value cutoff of 0.05. Gene sets are colored by category: GO biological process (BP, blue), GO cellular component (CC, yellow), and GO molecular function (MF, brown).

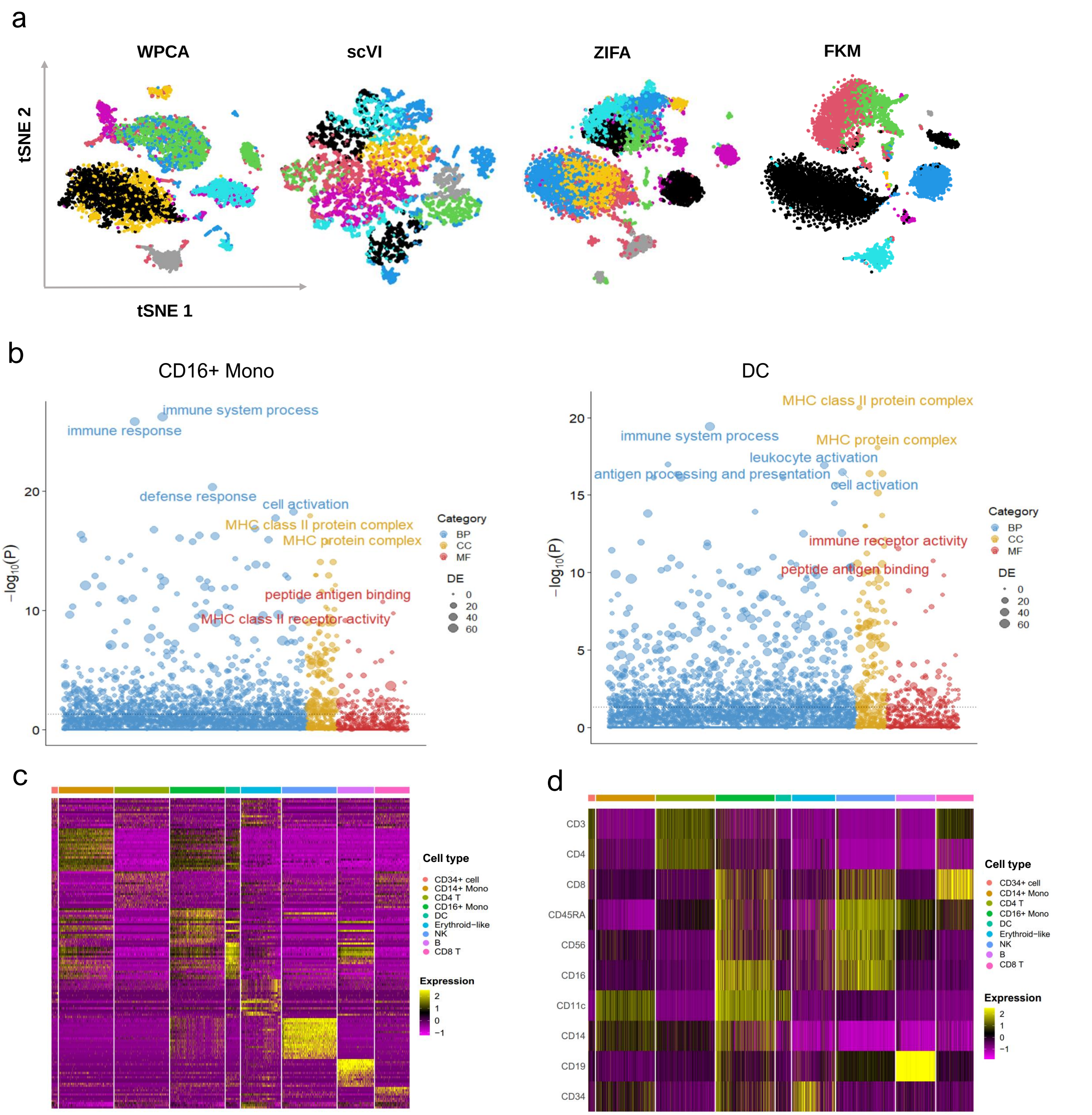

Figure S5: Results for cord blood mononuclear cell data: a. Scatter plot of tSNE based on 25-dimension latent features extracted by four dimension-reduction methods; b. Bubble plot of $-\log 10(p-\text{values})$ for pathway enrichment analysis of 87 and 105 DE genes of CD16+ monocytes (CD16+ Mono) and dendritic cells (DC), respectively, obtained by applying BPSC with two cutoffs: 0.05 for the adjusted $p$-value and 0.5 for the log fold-change. Dashed line represents a $p$-value cutoff of 0.05. Gene sets are colored by category: GO biological process (BP, blue), GO cellular component (CC, yellow), and GO molecular function (MF, brown); c. Heatmap of most significant differential gene expression levels for each cell type; d. Heatmap of protein levels of 10 cell-surface markers for each cell type.

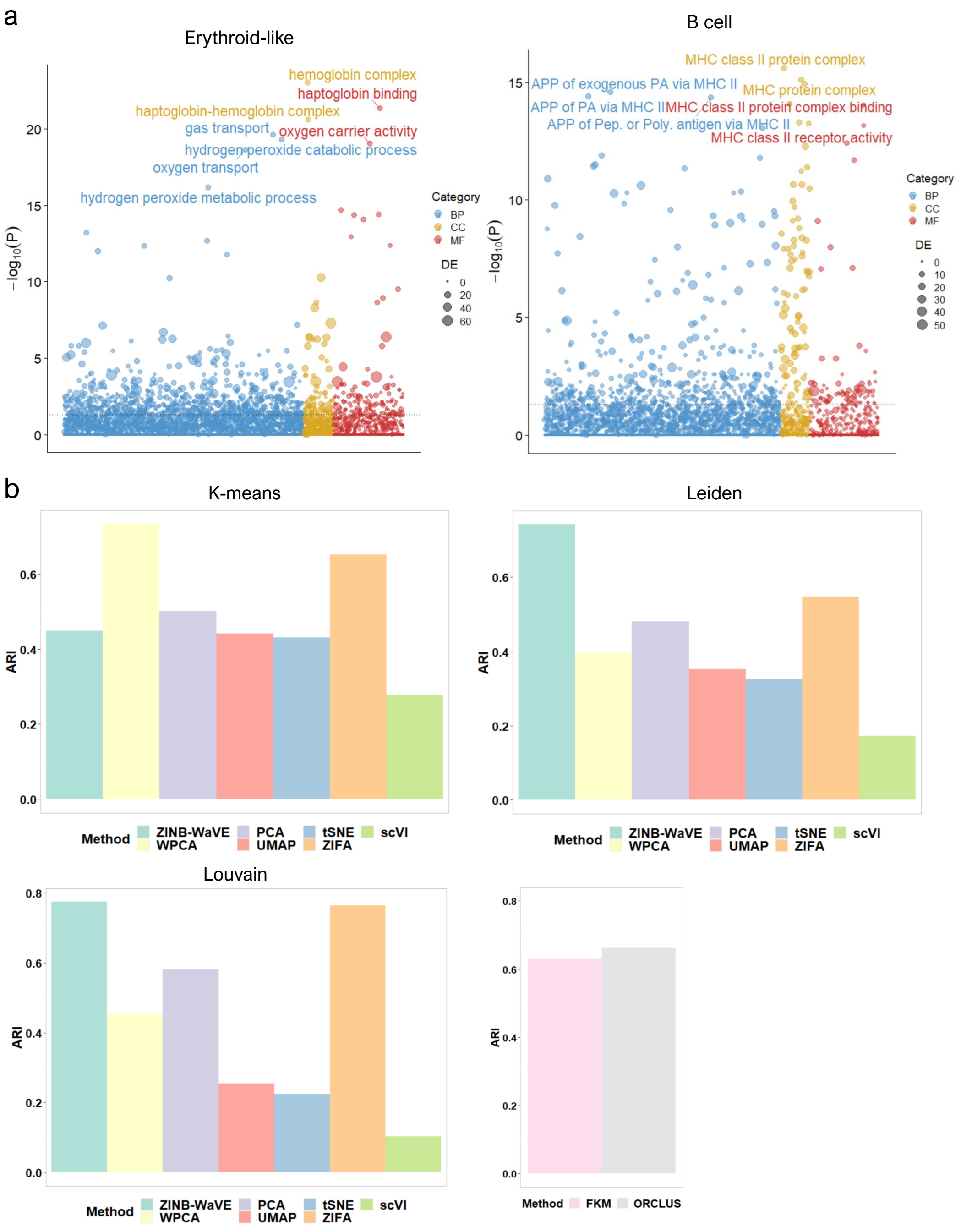

Figure S6: Results for cord blood mononuclear cell data: a. Bubble plot of $-\log10(p-\text{values})$ for pathway enrichment analysis of 326 and 67 DE genes of erythroid-like cells and B cells, respectively, obtained by applying *BPSC* with two cutoffs: 0.05 for the adjusted $p$-value and 0.5 for the log fold-change. Dashed line represents a $p$-value cutoff of 0.05. Gene sets are colored by category: GO biological process (BP, blue), GO cellular component (CC, yellow), and GO molecular function (MF, brown); b. Barplot of ARIs for three clustering methods, $k$-means, Leiden, and Louvain, based on seven dimension-reduction methods and two simultaneous dimension-reduction and clustering methods FKM and ORCLUS.

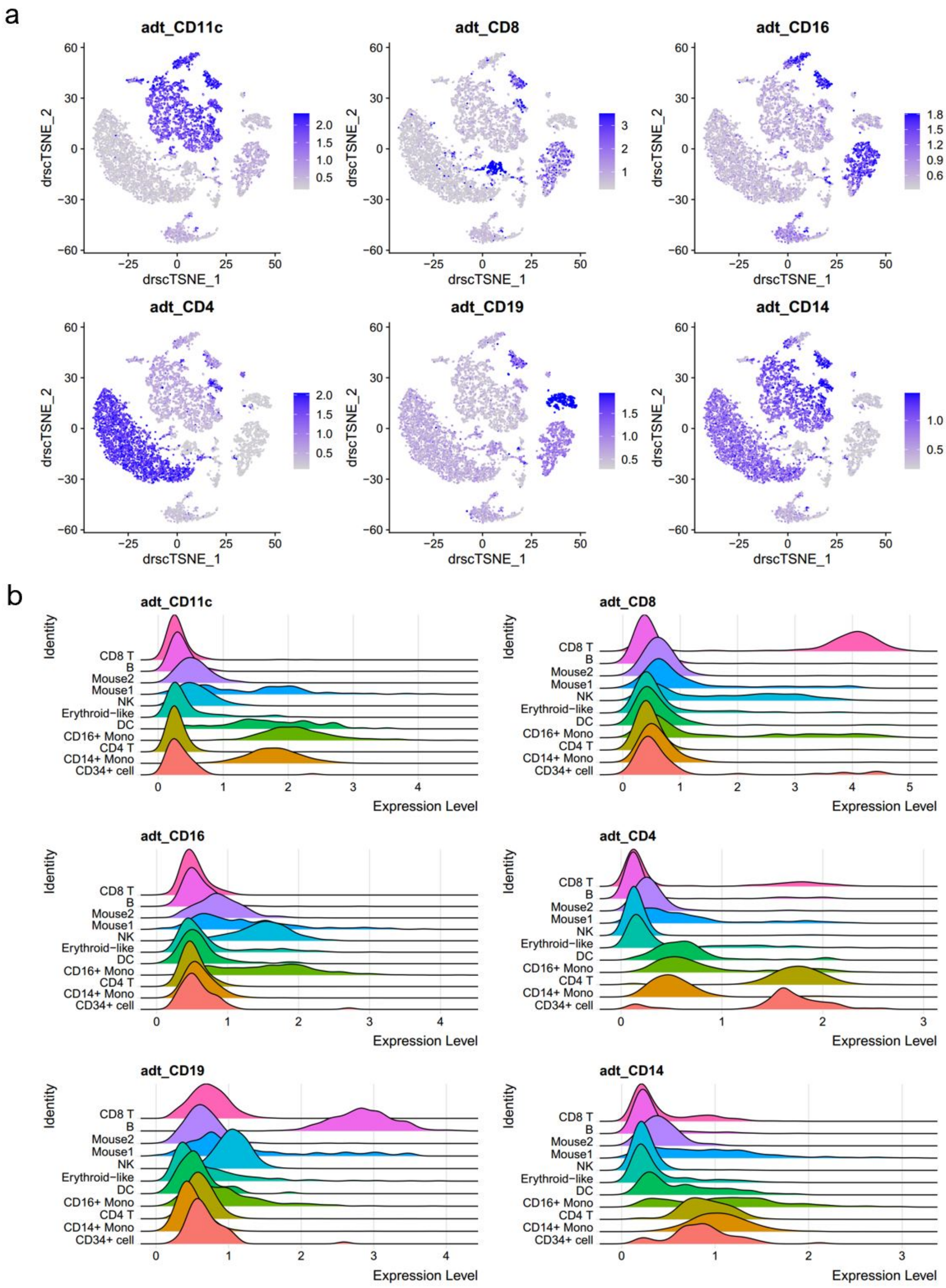

Figure S7: Protein levels of the first six cell-surface markers in cord blood mononuclear cells dataset: a. Scatter plots of tSNE based on the extracted 25-dimensional features from DR-SC colored by the protein levels of six cell-surface markers: CD11c, CD8, CD16, CD4, CD19, and CD14; b. Ridge plots of protein levels of the same six cell-surface markers.

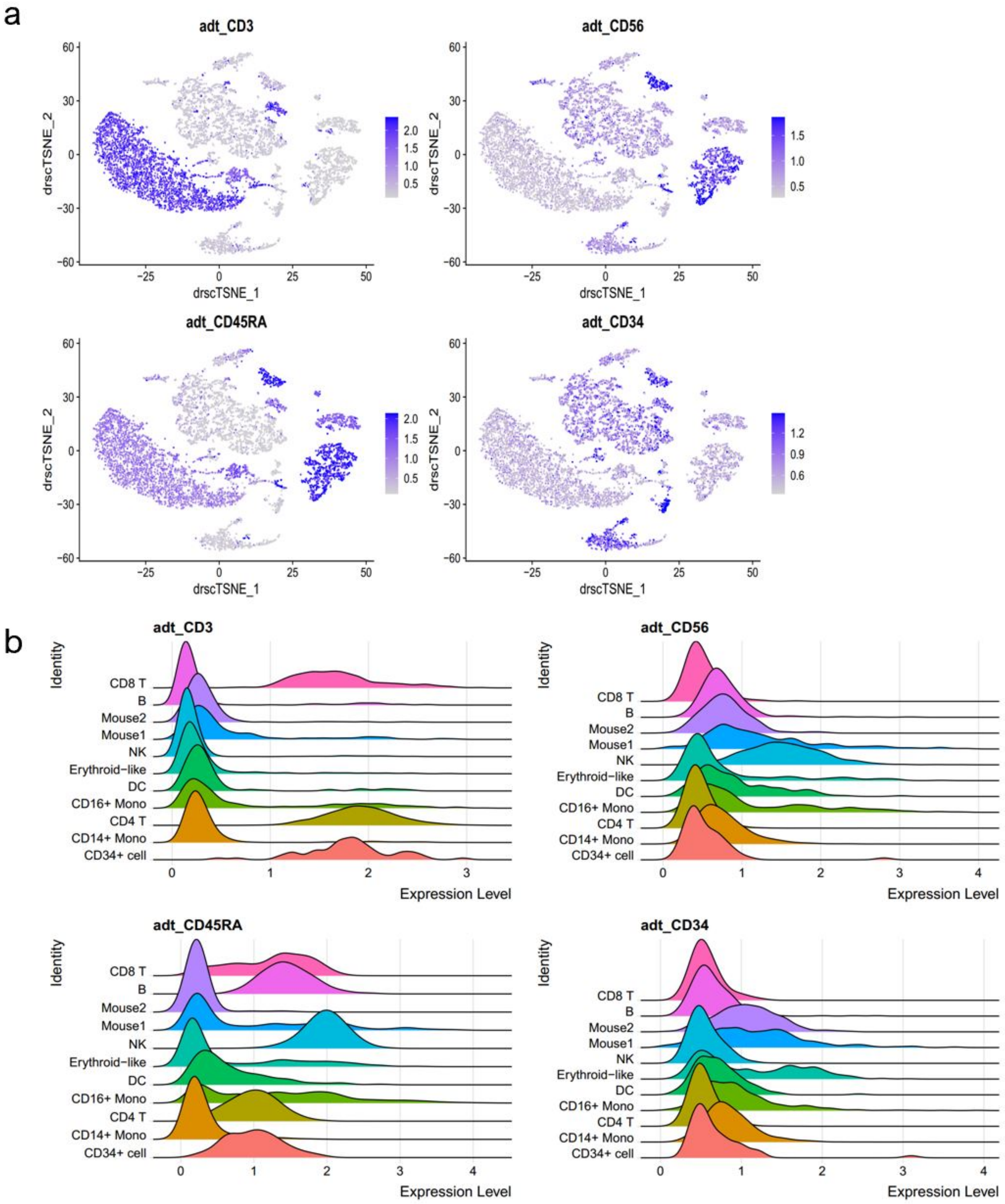

Figure S8: Protein levels of the last four cell-surface markers in cord blood mononuclear cells dataset: a. Scatter plots of tSNE based on the extracted 25-dimensional features from DR-SC colored by the protein levels of the four cell-surface markers: CD3, CD56, CD45RA, and CD34; b. Ridge plots of protein levels of the same four cell-surface markers.

### 2 Supplementary Figures in Main Text

#### 2.1 Additional Simulation Results

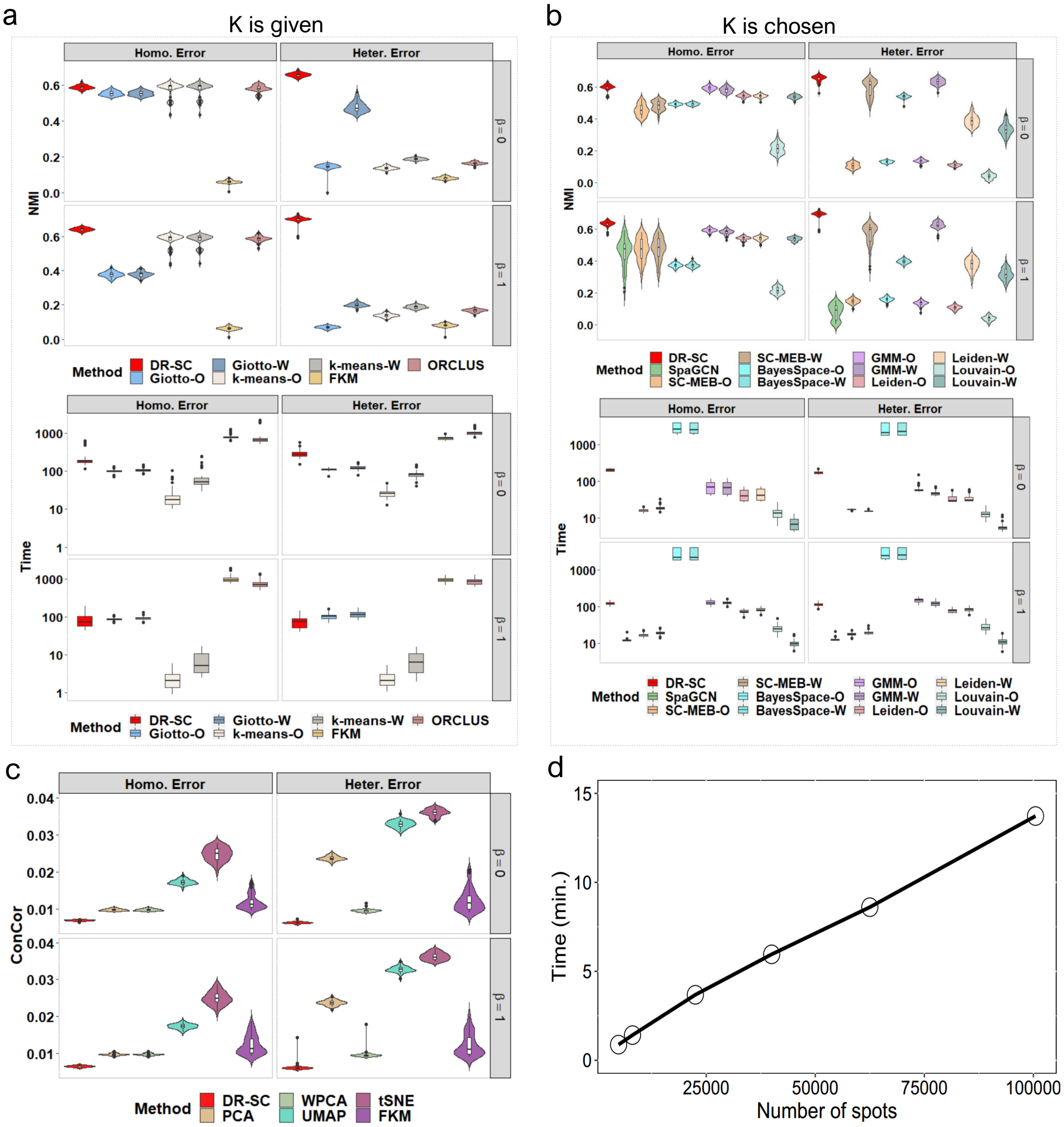

Figure S9: Comparison of 18 clustering methods and six dimension-reduction methods in the Simulation 1: a. Violin plot of NMIs and boxplot of running time for DR-SC and the other six methods that cannot choose the number of clusters in a data-driven way, where running time is shown in seconds and the y-axis is log-scale; b. Violin plot of NMIs and boxplot of running time for DR-SC and the other 11 methods that can automatically choose the number of clusters; c. Violin plot of the conditional correlations $y_i$ and $\mathbf{x}_i$ given the extracted latent features from different methods; d. Line plot between running time and number of spots by running 30 iterations per dataset and using 6.6G memory on a machine with 2.10GHz Intel(R) Xeon(R) Gold 6230 CPU.

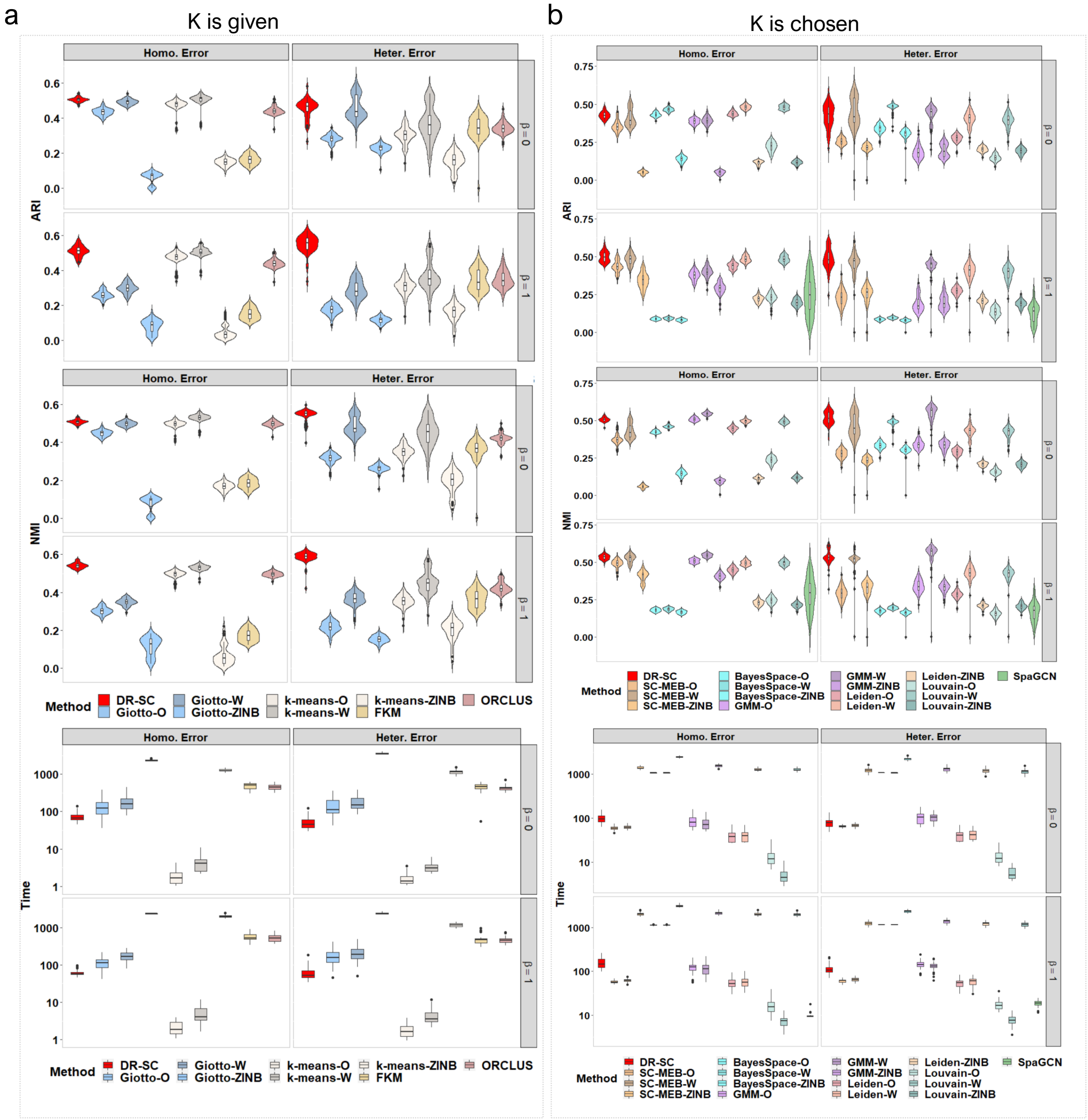

Figure S10: Comparison of 25 clustering methods in Simulation 2: a. Violin plots of ARIs and NMIs and boxplot of running time for DR-SC and the other eight methods that cannot choose the number of clusters in a data-driven way, where running time is shown in seconds and the y-axis is log-scale; b. Violin plots of ARIs and NMIs and boxplot of running time for DR-SC and the other 16 methods that can automatically choose the number of clusters.

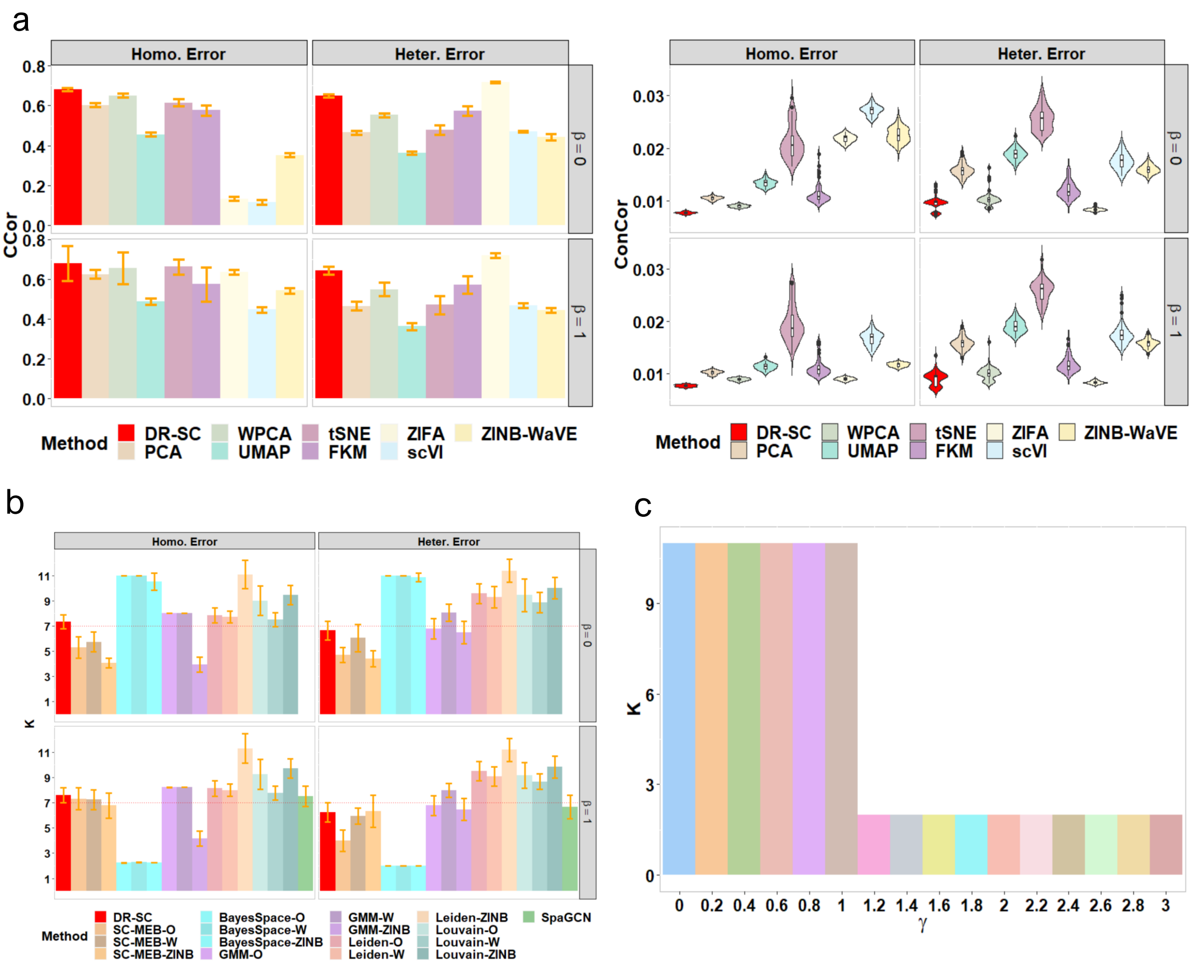

Figure S11: Comparison of 17 clustering methods and nine dimension-reduction methods in Simulation: a. Bar plot of the average canonical correlation coefficients for the true latent features and the estimated ones from different methods over 50 runs, and violin plot of the conditional correlation coefficients $y_i$ and $\mathbf{x}_i$ given the extracted latent features from DR-SC and the other eight different methods in Simulation 2; b. Bar plot of average estimated number of clusters over 50 runs from DR-SC and other 16 methods in Simulation 2, where the red dashed line denotes the true number of clusters; c. Bar plot of average number of clusters chosen by BayesSpace over 10 runs varied with the smoothing parameters in *BayesSpace* R function for the spatial case in Simulation 1.

#### 2.2 Additional results for human dorsolateral prefrontal cortex datasets

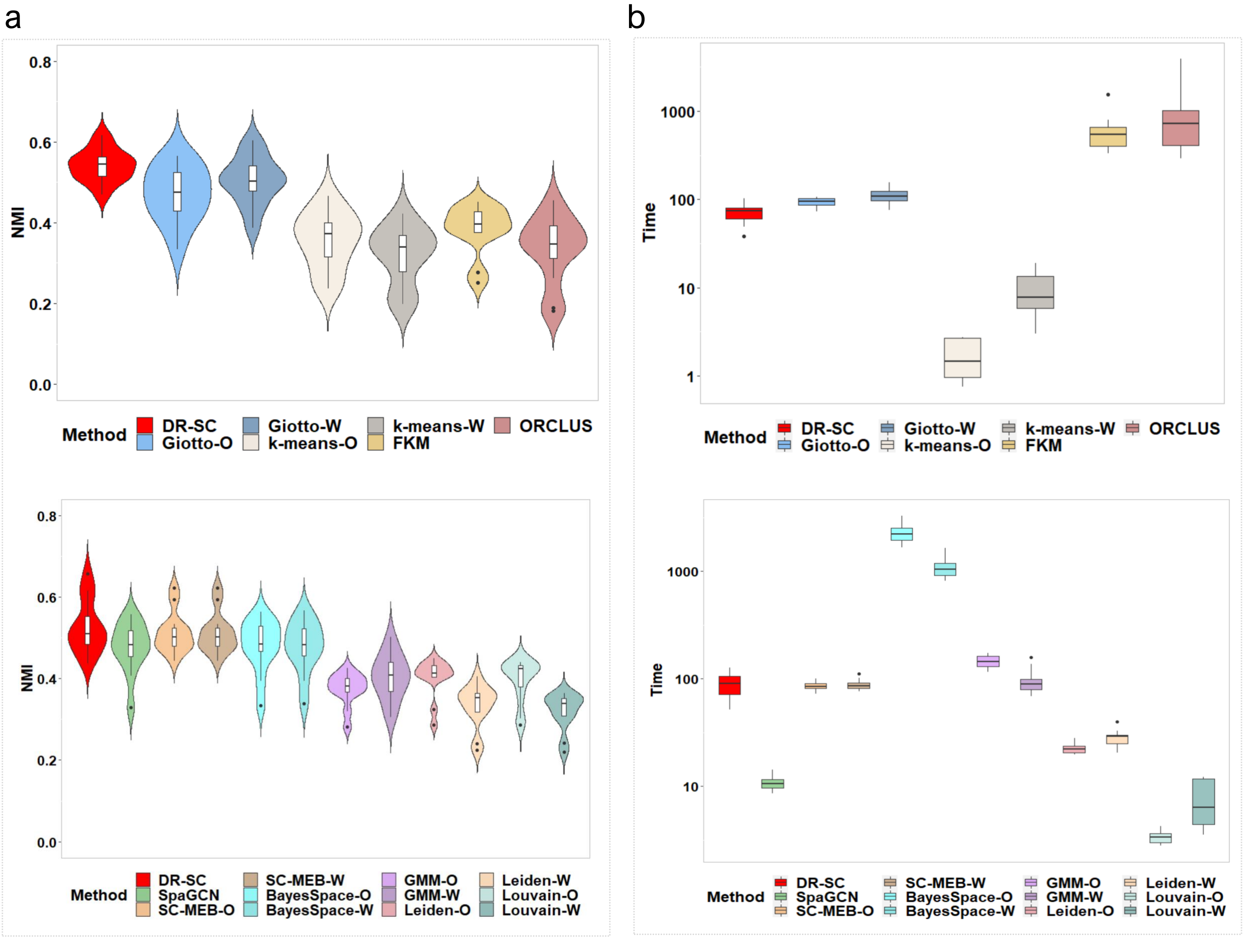

Figure S12: Clustering performance for human dorsolateral prefrontal cortex dataset: a. Violin plot of ARIs for seven clustering methods given the true number of clusters and 12 clustering methods in which the number of clusters is chosen in a method-specific way across 12 tissue samples; b. Boxplot of running time (unit: seconds) for seven clustering methods given the true number of clusters and 12 clustering methods that choose the number of clusters in a method-specific way across 12 tissue samples.

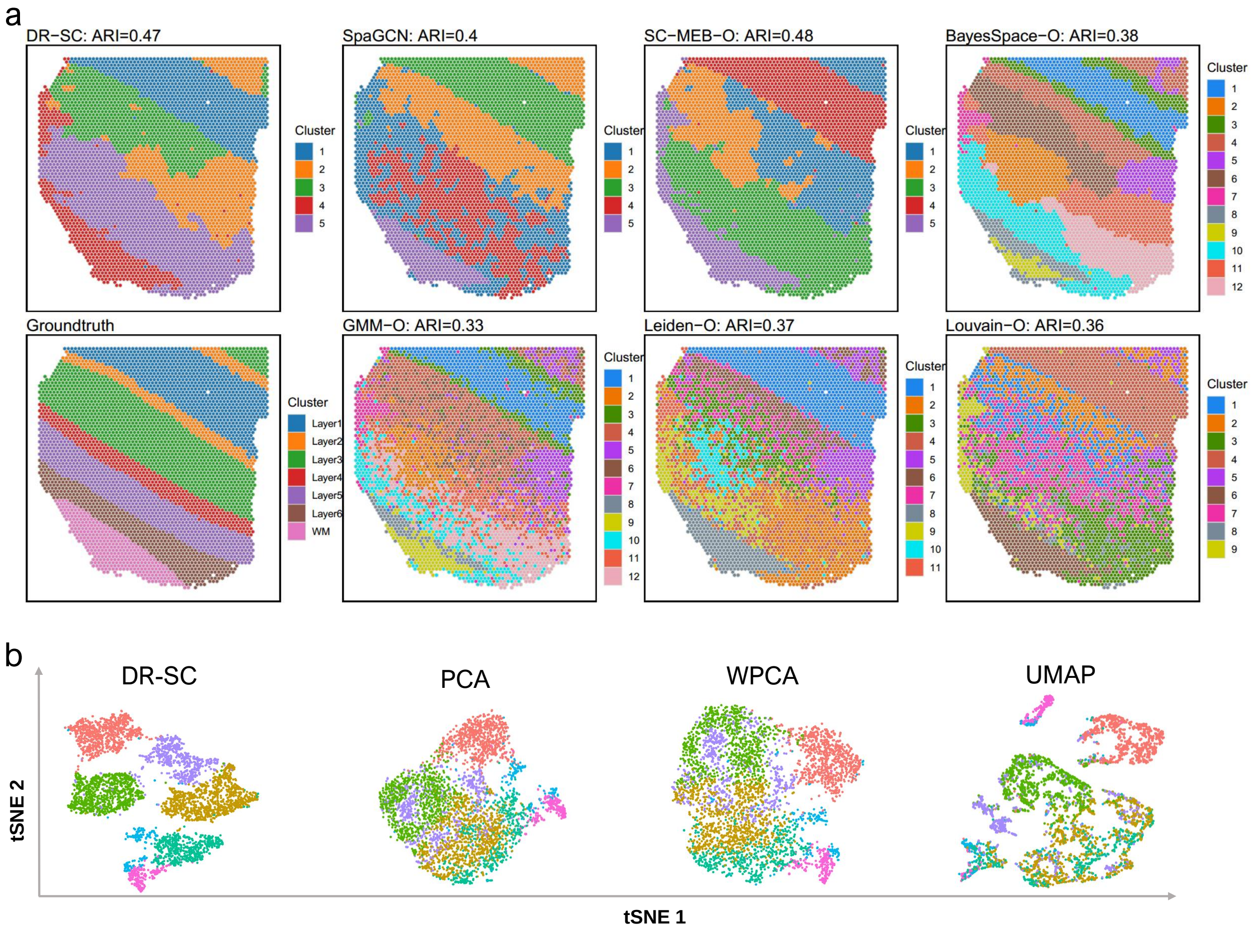

Figure S13: Application of DR-SC in the clustering and visualization of dataset ID151507 from human postmortem DLPFC tissue sections, in which the cell types had been manually annotated. a. Spatial heat map of clusters from different clustering methods based on PCA that can choose the number of clusters. b. Visualization of the cluster labels from DR-SC based on two-dimensional tSNE embeddings from four different DR methods: DR-SC, PCA, WPCA, and UMAP.

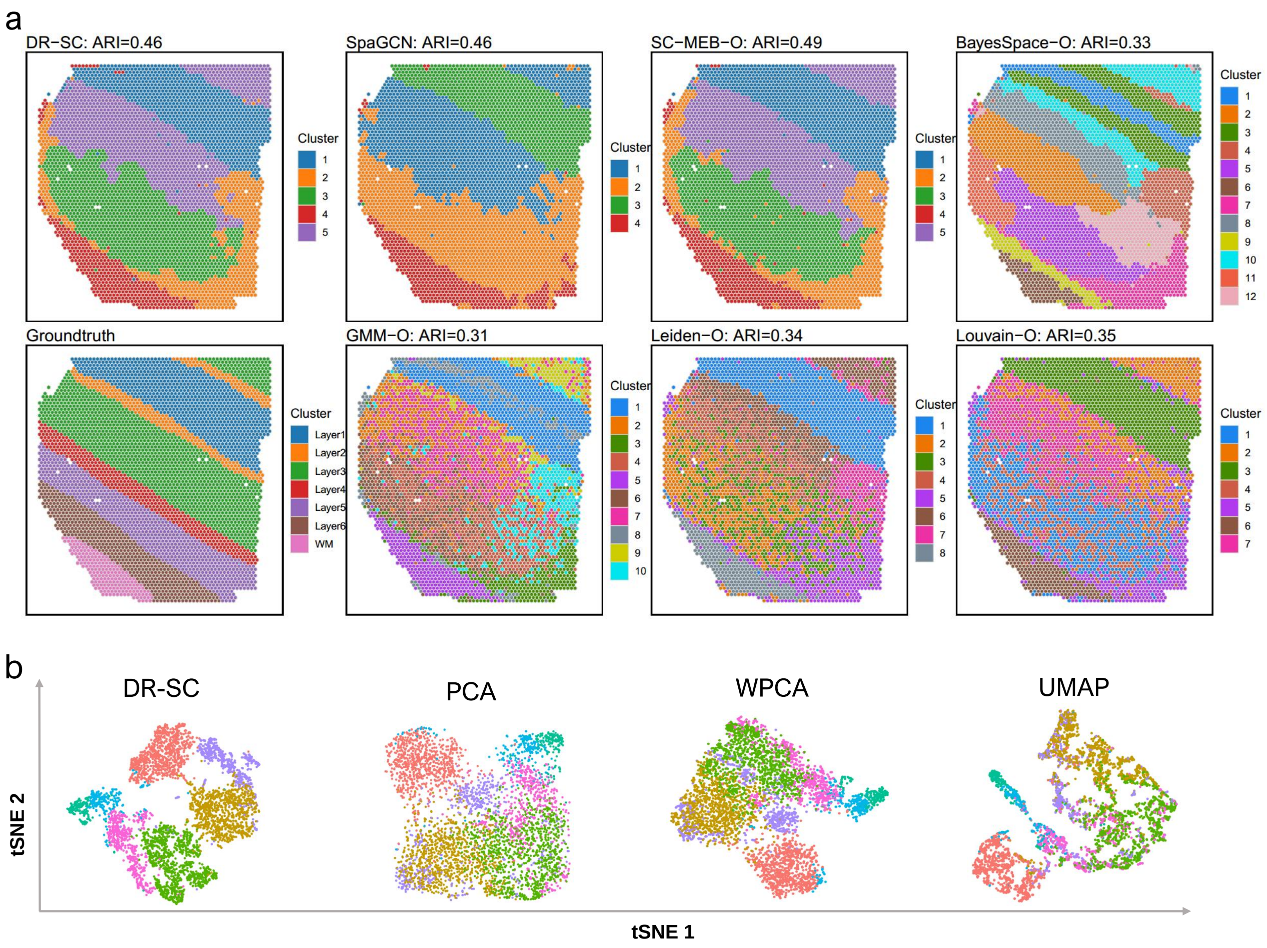

Figure S14: Application of DR-SC to clustering and visualization of dataset ID151508 from human postmortem DLPFC tissue sections, in which the cell types had been manually annotated. a. Spatial heat map of clusters from different clustering methods based on PCA that can choose the number of clusters. b. Visualization of the cluster labels from DR-SC based on two-dimensional tSNE embeddings from four different DR methods: DR-SC, PCA, WPCA, and UMAP.

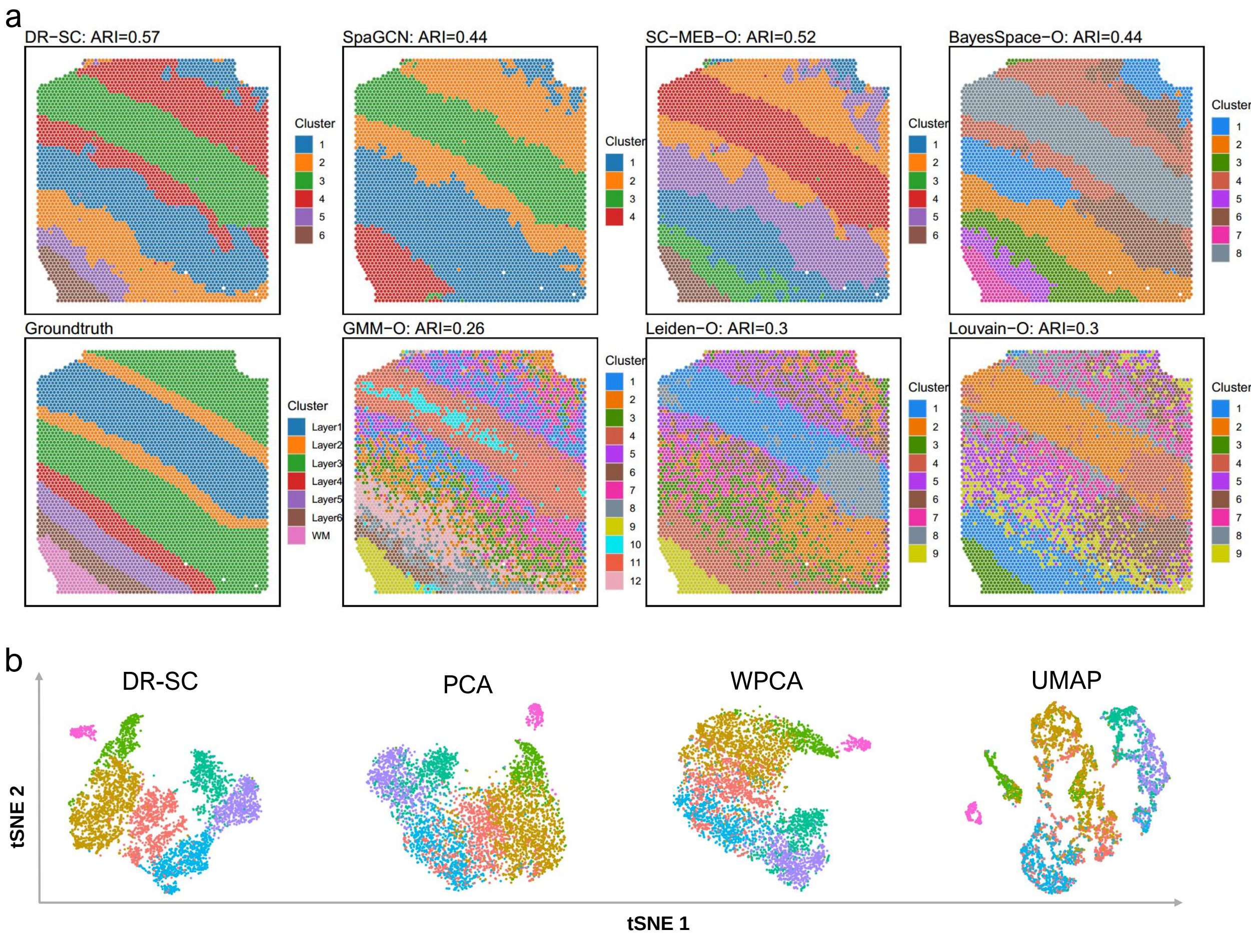

Figure S15: Application of DR-SC to clustering and visualization of dataset ID151509 from human postmortem DLPFC tissue sections, in which the cell types had been manually annotated. a. Spatial heat map of clusters from different clustering methods based on PCA that can choose the number of clusters. b. Visualization of the cluster labels from DR-SC based on two-dimensional tSNE embeddings from four different DR methods: DR-SC, PCA, WPCA, and UMAP.

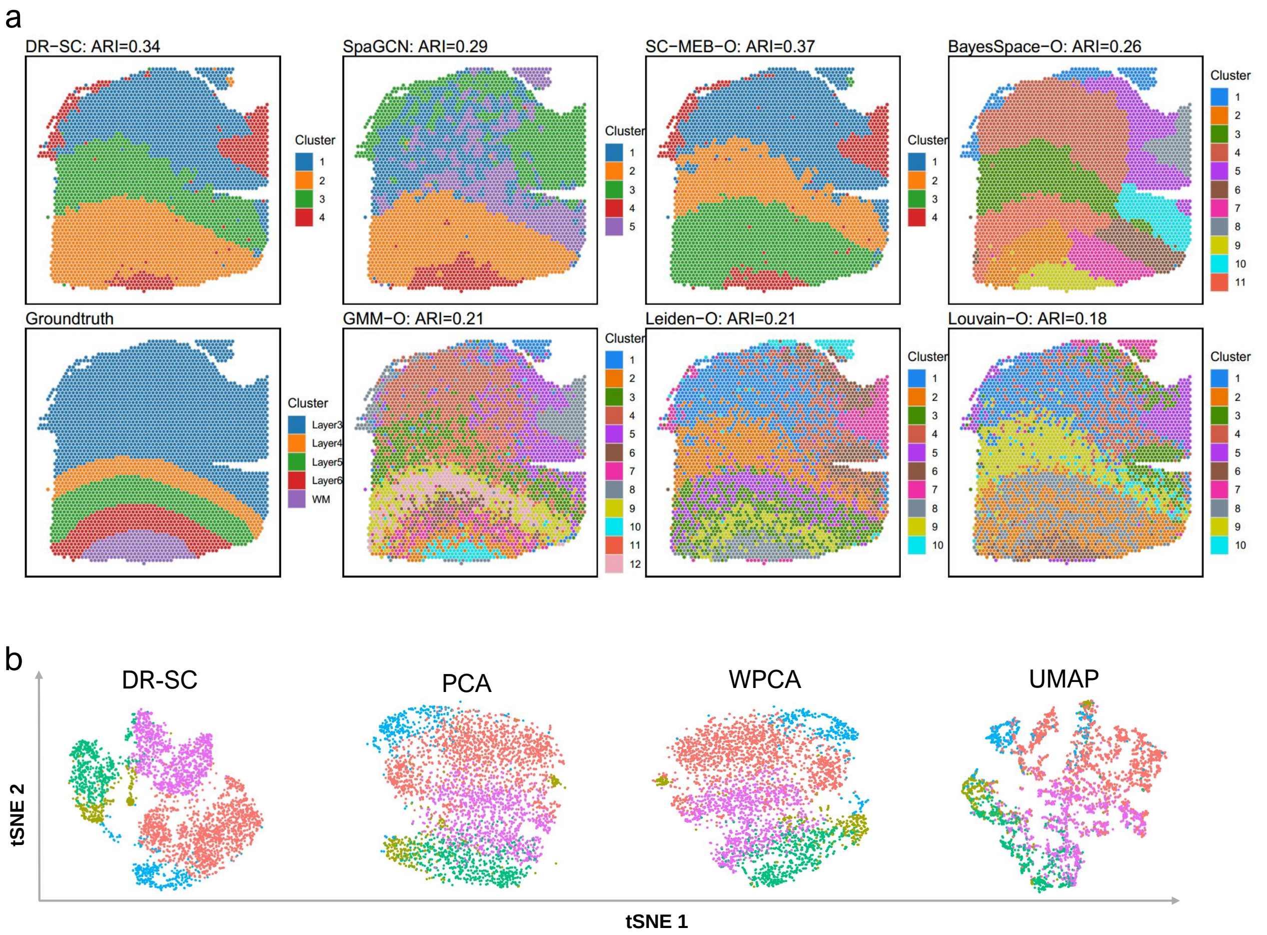

Figure S16: Application of DR-SC in clustering and visualization of dataset ID151669 from human postmortem DLPFC tissue sections, in which the cell types had been manually annotated. a. Spatial heat map of clusters from different clustering methods based on PCA that can choose the number of clusters. b. Visualization of the cluster labels from DR-SC based on two-dimensional tSNE embeddings from four different DR methods: DR-SC, PCA, WPCA, and UMAP.

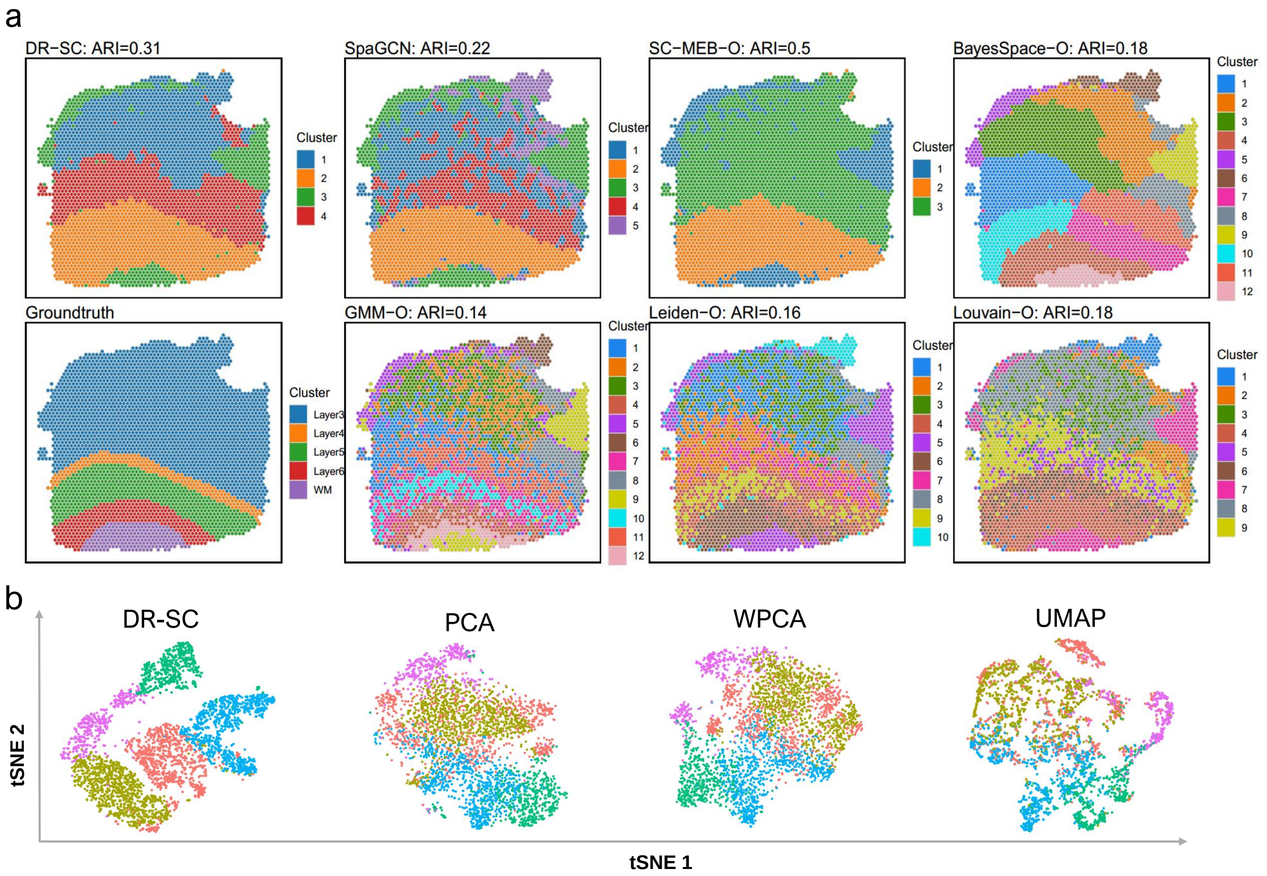

Figure S17: Application of DR-SC in clustering and visualization of dataset ID151670 from human postmortem DLPFC tissue sections, in which the cell types had been manually annotated. a. Spatial heat map of clusters from different clustering methods based on PCA that can choose the number of clusters. b. Visualization of the cluster labels from DR-SC based on two-dimensional tSNE embeddings from four different DR methods: DR-SC, PCA, WPCA, and UMAP.

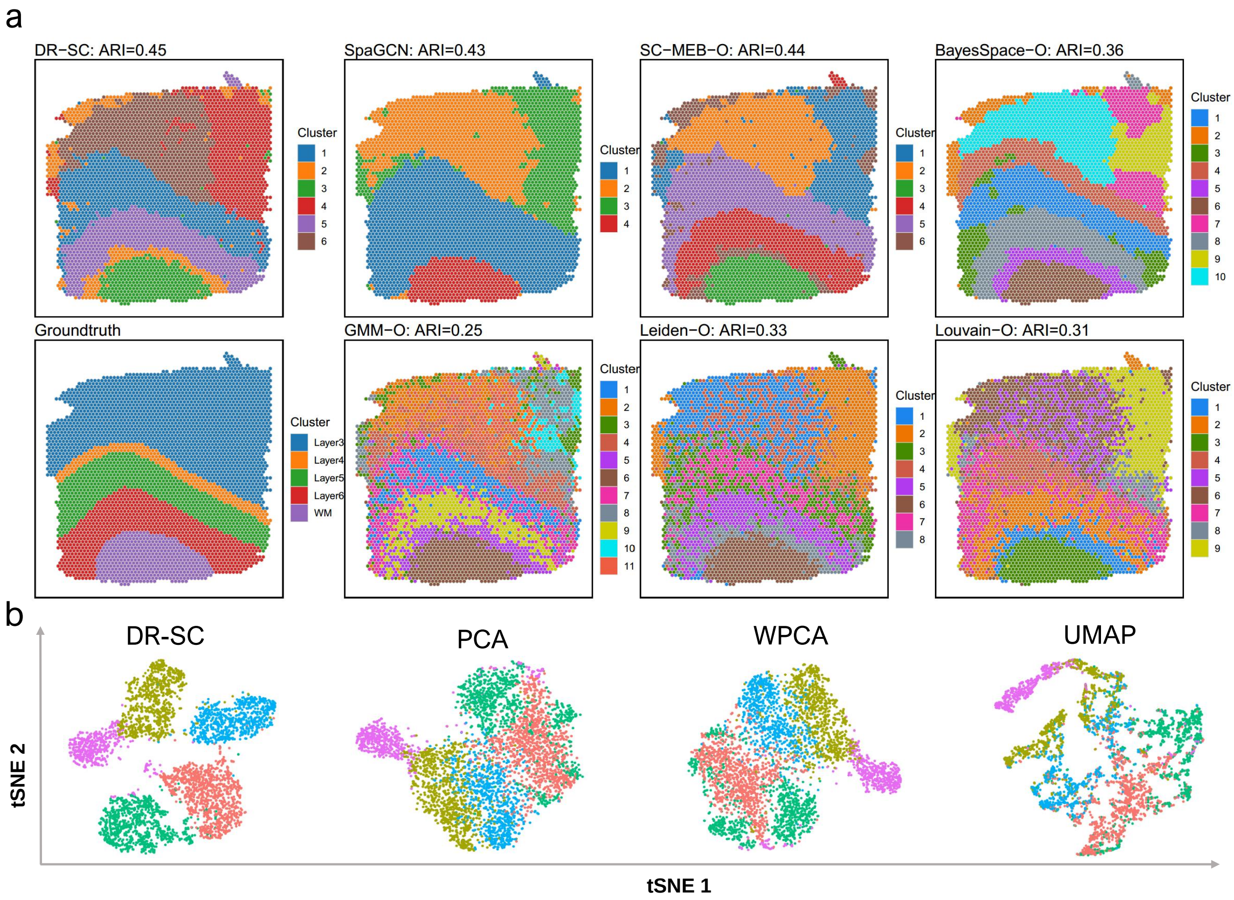

Figure S18: Application of DR-SC to clustering and visualization of dataset ID151671 from human postmortem DLPFC tissue sections, in which the cell types had been manually annotated. a. Spatial heat map of clusters from different clustering methods based on PCA that can choose the number of clusters. b. Visualization of the cluster labels from DR-SC based on two-dimensional tSNE embeddings from four different DR methods: DR-SC, PCA, WPCA, and UMAP.

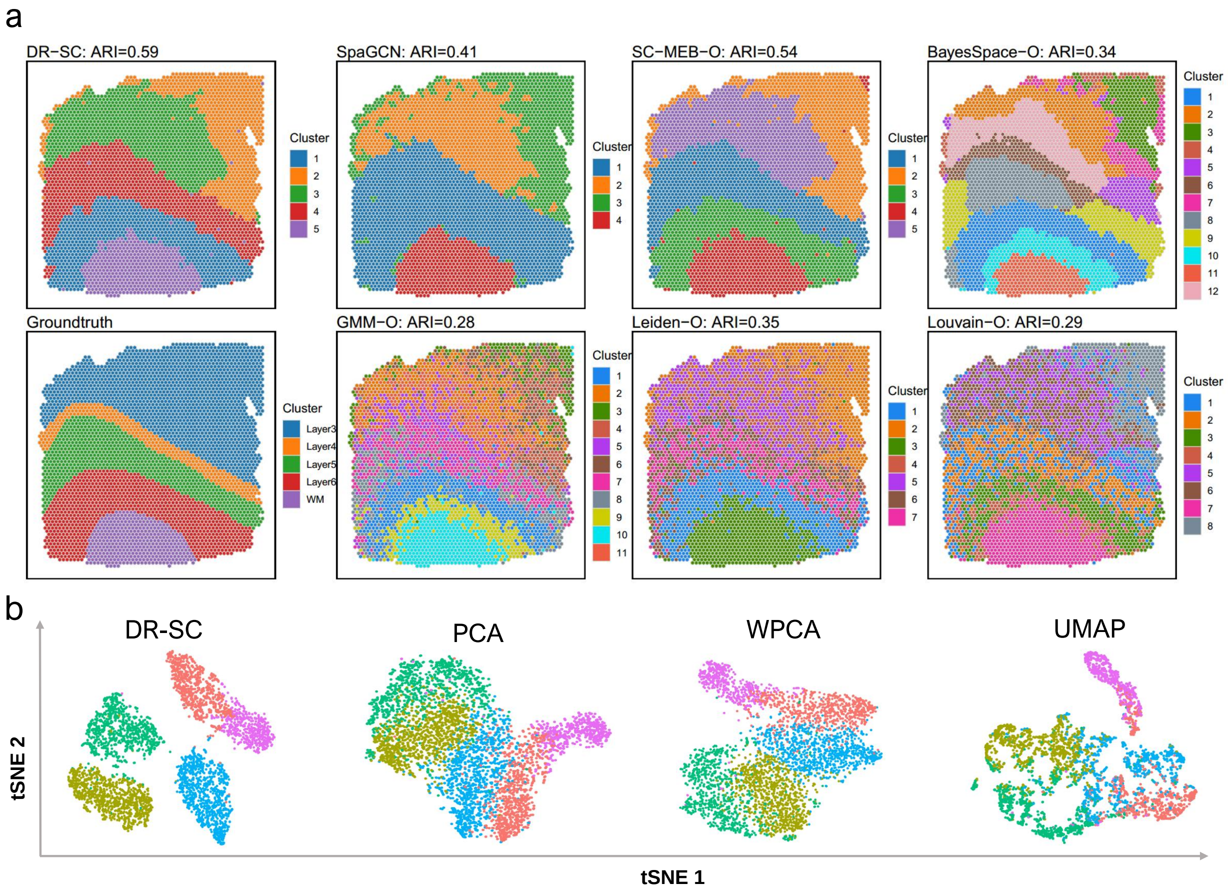

Figure S19: Application of DR-SC to clustering and visualization of dataset ID151672 from human postmortem DLPFC tissue sections, in which the cell types had been manually annotated. a. Spatial heat map of clusters from different clustering methods based on PCA that can choose the number of clusters. b. Visualization of the cluster labels from DR-SC based on two-dimensional tSNE embeddings from four different DR methods: DR-SC, PCA, WPCA, and UMAP.

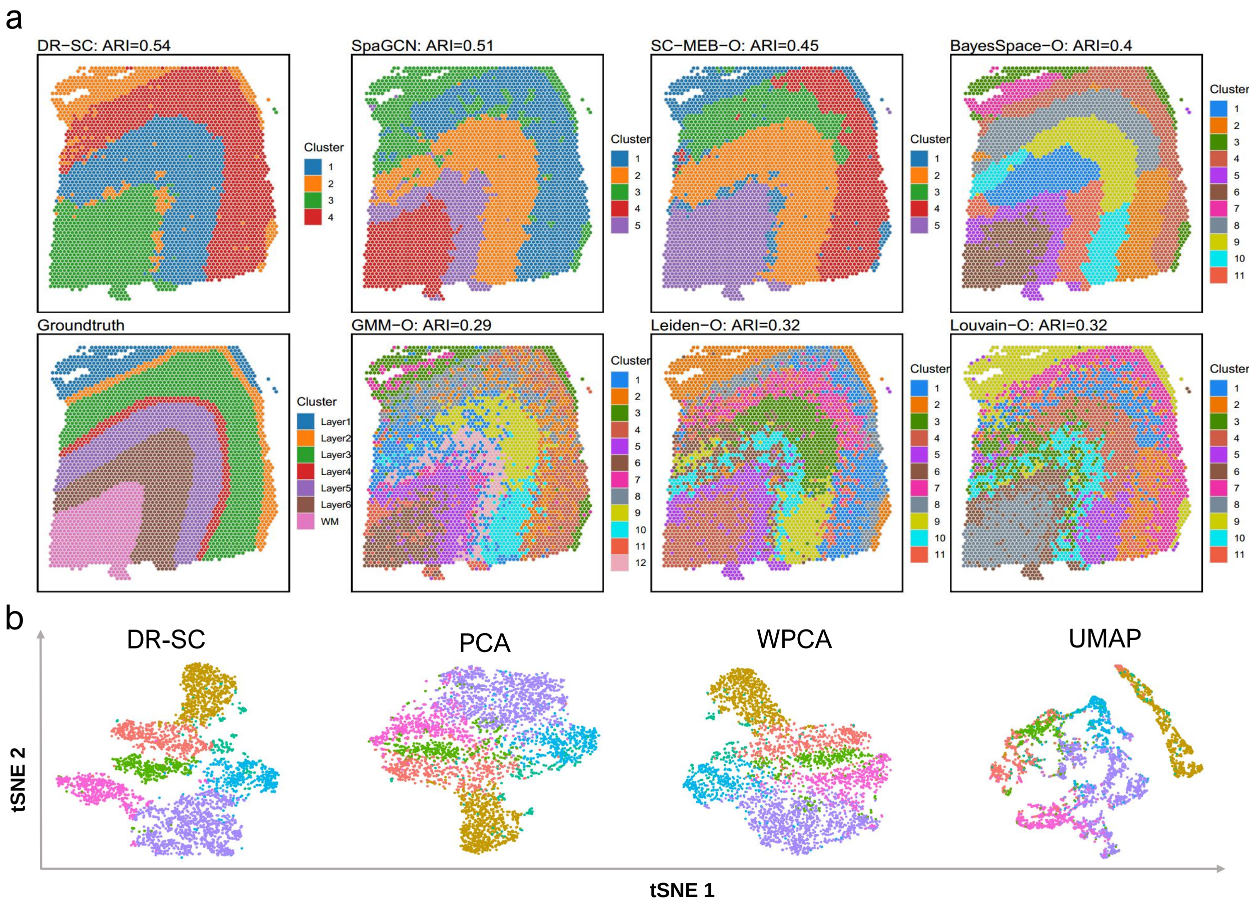

Figure S20: Application of DR-SC to clustering and visualization of dataset ID151673 from human postmortem DLPFC tissue sections, in which the cell types had been manually annotated. a. Spatial heat map of clusters from different clustering methods based on PCA that can choose the number of clusters. b. Visualization of the cluster labels from DR-SC based on two-dimensional tSNE embeddings from four different DR methods: DR-SC, PCA, WPCA, and UMAP.

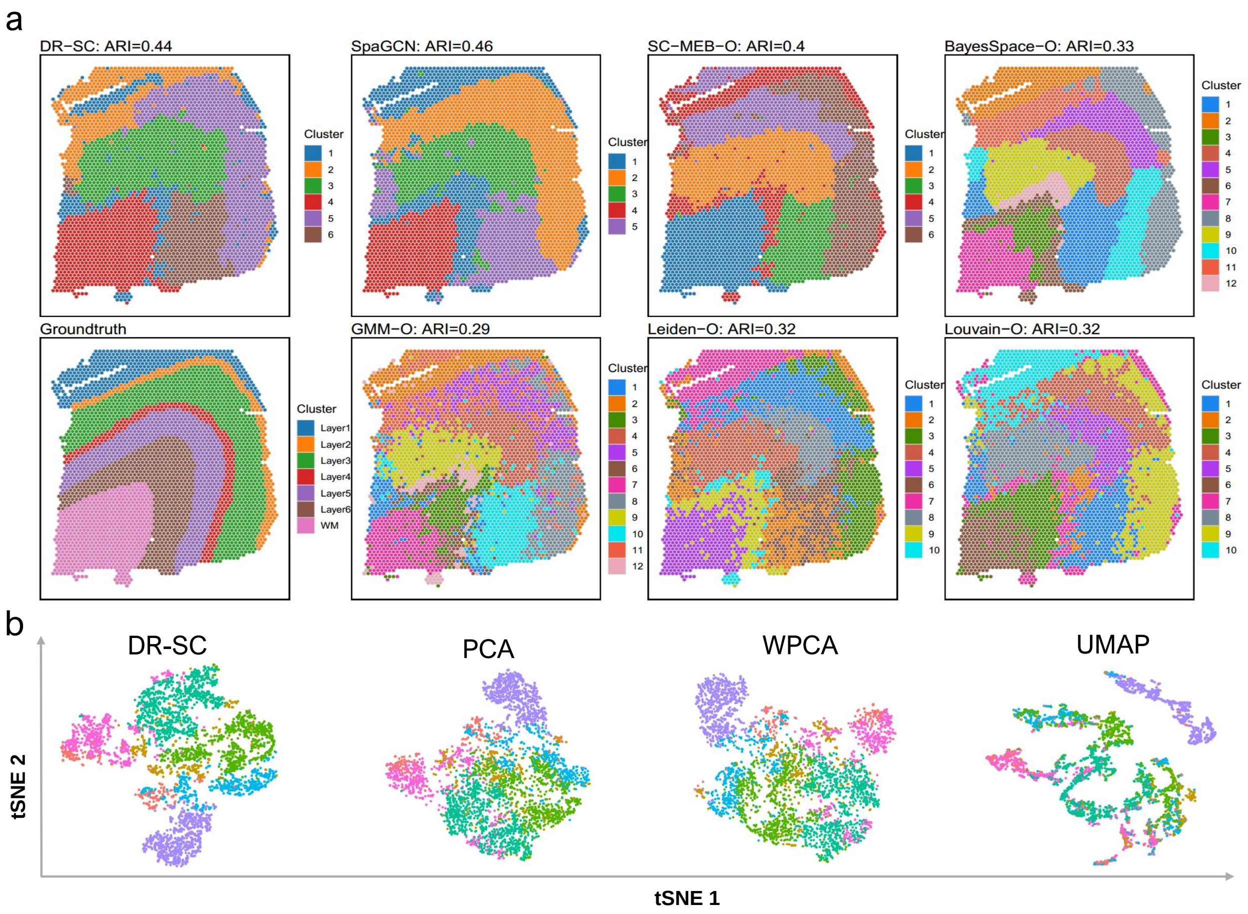

Figure S21: Application of DR-SC to clustering and visualization of dataset ID151674 from human postmortem DLPFC tissue sections, in which the cell types had been manually annotated. a. Spatial heat map of clusters from different clustering methods based on PCA that can choose the number of clusters. b. Visualization of the cluster labels from DR-SC based on two-dimensional tSNE embeddings from four different DR methods: DR-SC, PCA, WPCA, and UMAP.

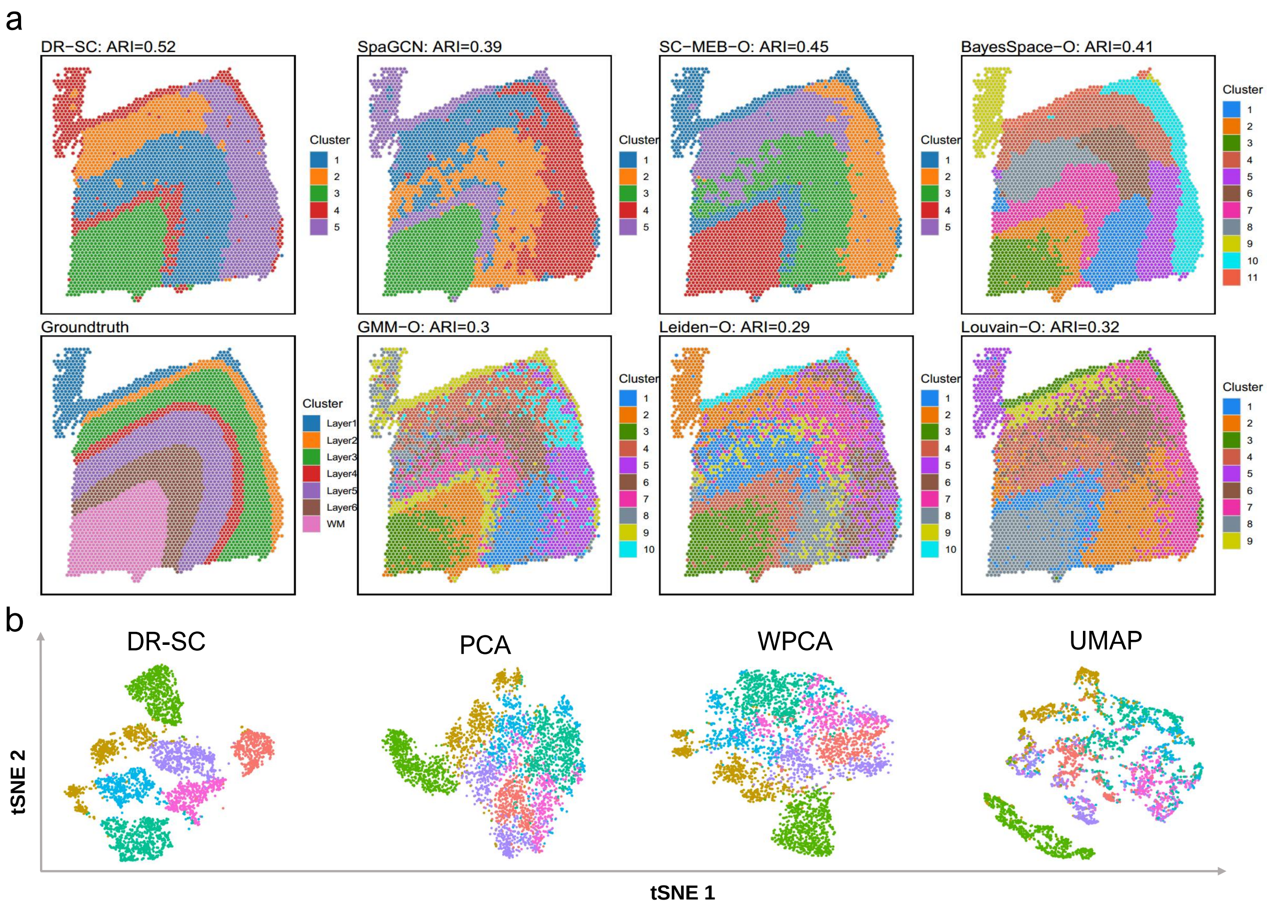

Figure S22: Application of DR-SC to clustering and visualization of dataset ID1511675 from human postmortem DLPFC tissue sections, in which the cell types had been manually annotated. a. Spatial heat map of clusters from different clustering methods based on PCA that can choose the number of clusters. b. Visualization of the cluster labels from DR-SC based on two-dimensional tSNE embeddings from four different DR methods: DR-SC, PCA, WPCA, and UMAP.

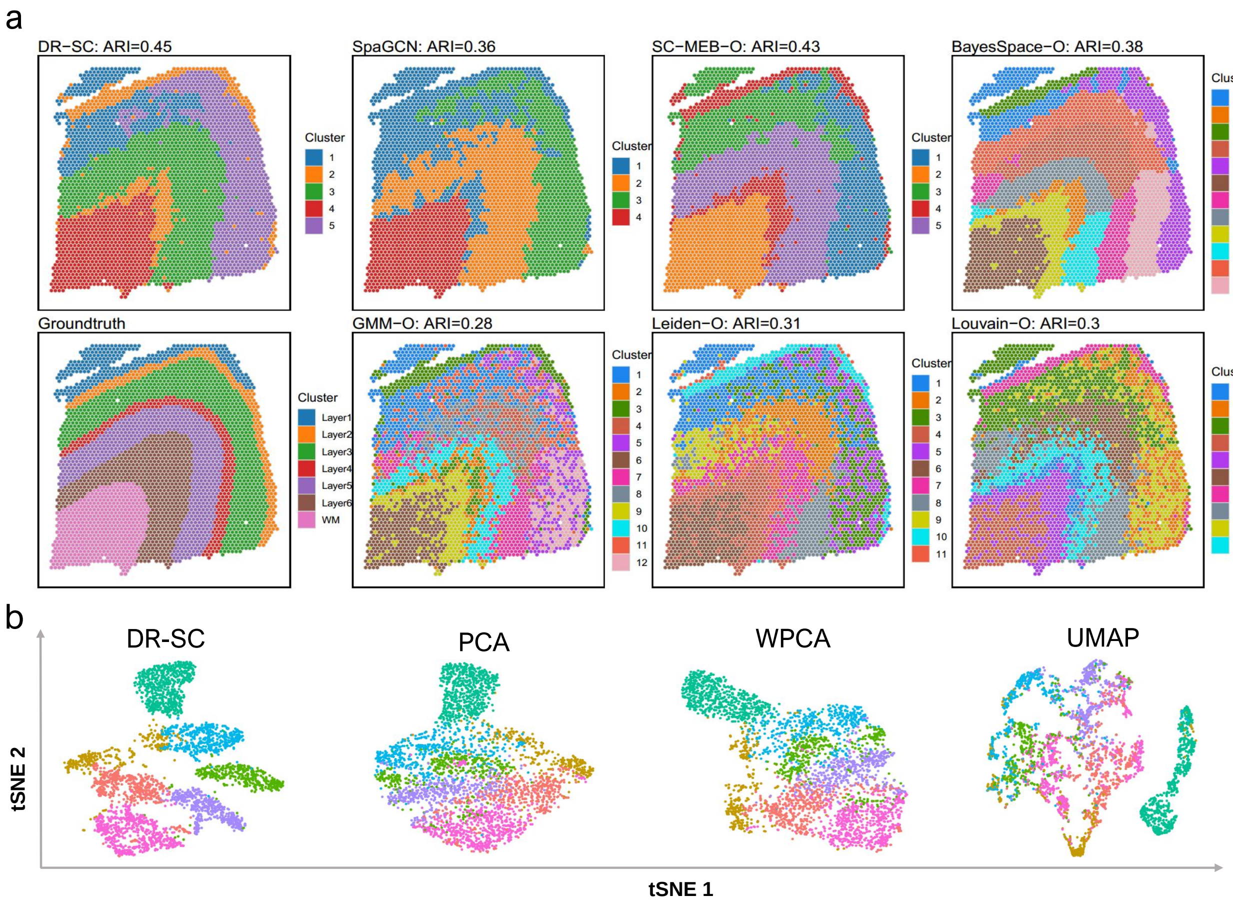

Figure S23: Application of DR-SC to clustering and visualization of dataset ID151676 from human postmortem DLPFC tissue sections, in which the cell types had been manually annotated. a. Spatial heat map of clusters from different clustering methods based on PCA that can choose the number of clusters. b. Visualization of the cluster labels from DR-SC based on two-dimensional tSNE embeddings from four different DR methods: DR-SC, PCA, WPCA, and UMAP.

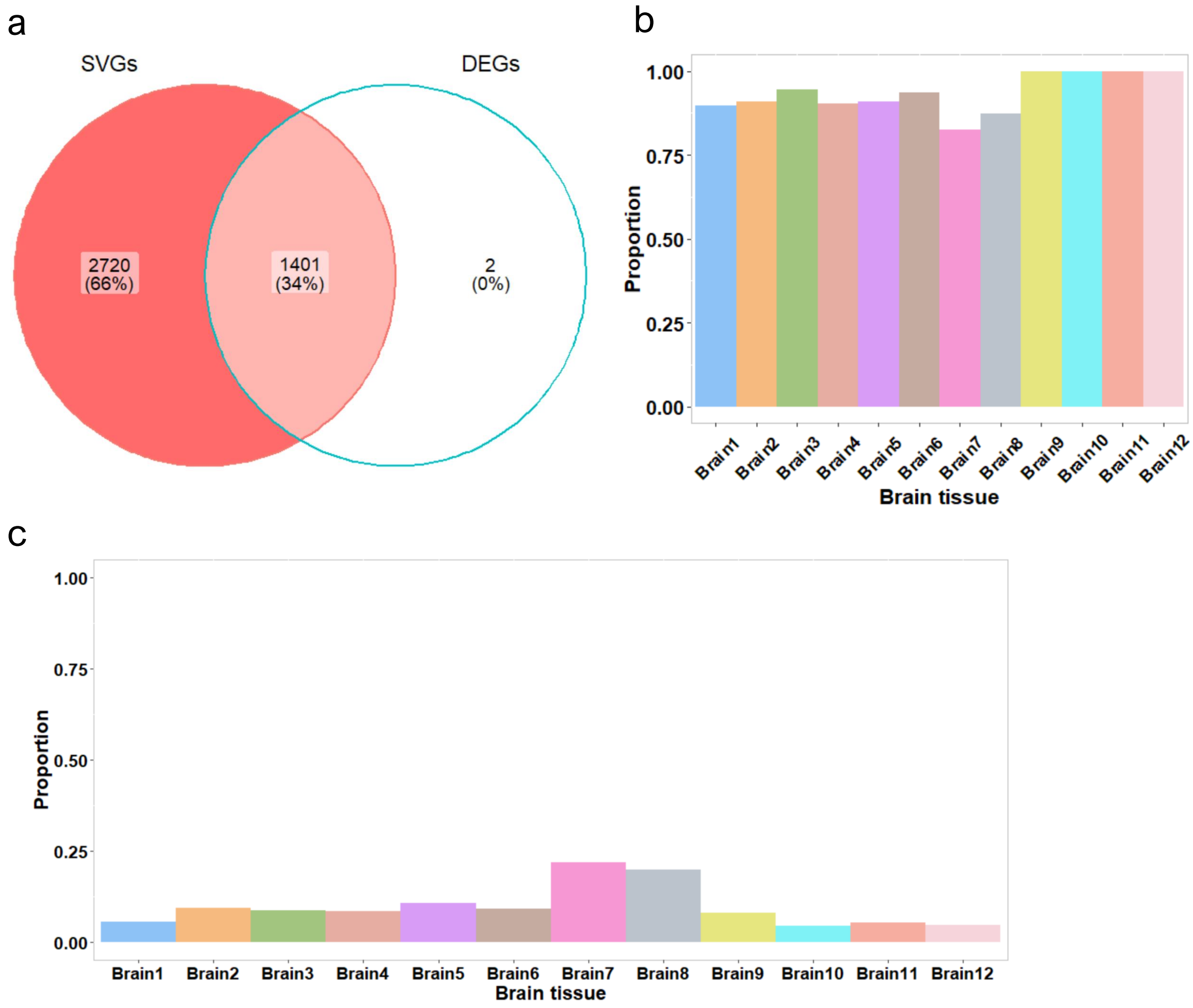

Figure S24: Comparison of SVGs with DEGs from DLPFC datasets. a. Vennplot of overlapping SVGs without adjusting for cell-type-relevant covariates and DEGs across 12 DLPFC tissue sections; b. Bar plot of overlapping proportions for DE genes by comparison with SVGs without controlling PCs in each DLPFC section; c. Bar plot of overlapping proportion for DE genes by comparison with SVGs while controlling PCs in each DLPFC section.

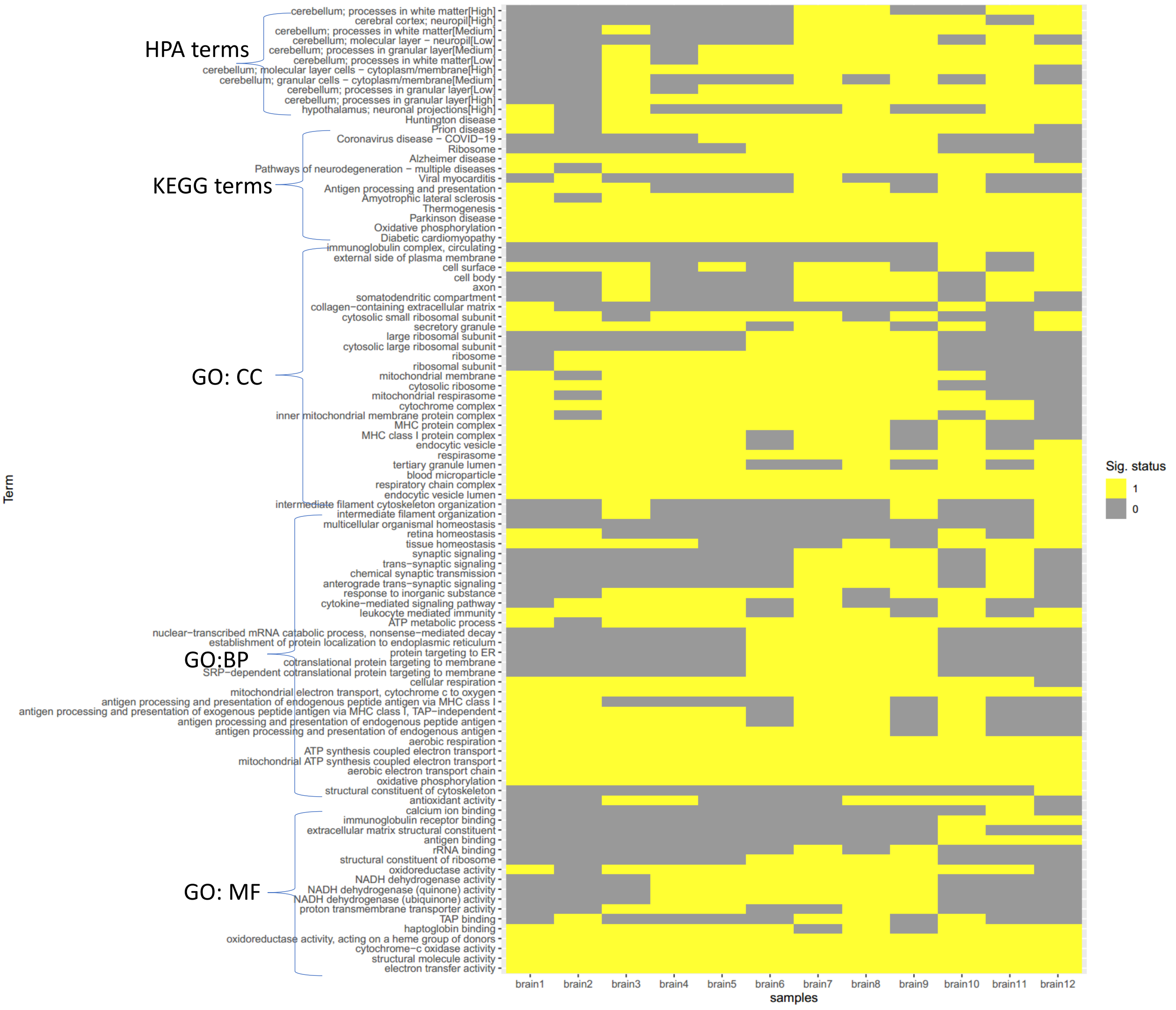

Figure S25: Top 5 enriched pathways across 12 human dorsolateral prefrontal cortex datasets, in which we identified genes with spatial expression variation for each dataset at an FDR of 1% by controlling for cell-type-related covariates, i.e., estimated latent features from DR-SC, followed by gene set enrichment analysis was performed on theses genes. X-axis shows each sample numbered from brain 1 to brain 12. Y-axis denotes the top 5 pathways for each dataset. The significant pathways (Sig. status) for each dataset are in yellow.

#### 2.3 Additional results for mouse olfactory bulb dataset

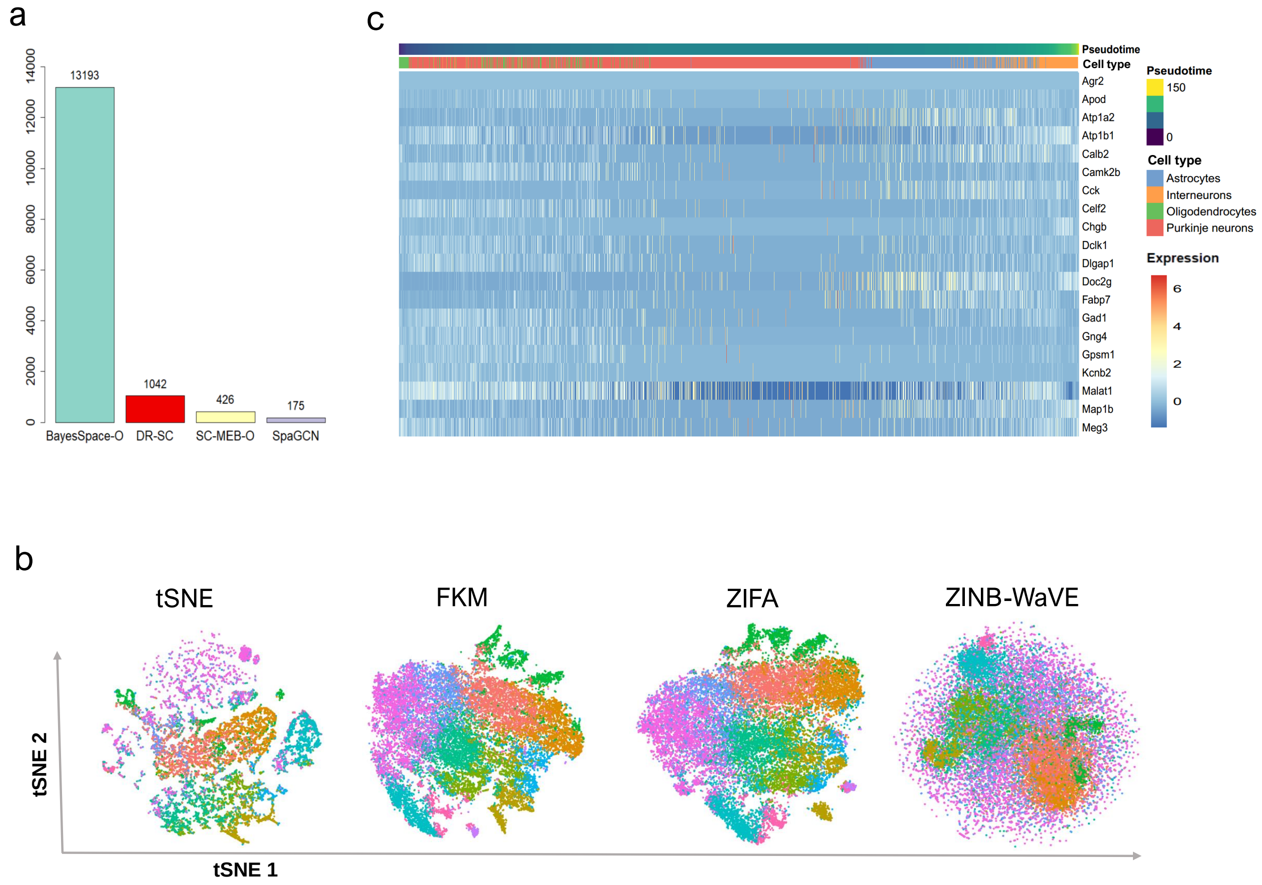

Figure S26: Analysis of mouse olfactory bulb data. a. Bar plot of running time (seconds) for DR-SC and three spatial-clustering methods, SpaGCN, SC-MEB, and BayesSpace, based on PCA. b. Visualization of cluster labels from DR-SC based on two-dimensional tSNE embeddings from four DR methods, tSNE, FKM, ZIFA, and ZINB-WaVE. c. Heatmap of top 20 significantly changed genes expression levels with respect to the Slingshot pseudotime. Each column represents a spot that is mapped to this path and is ordered by its pseudotime value. Each row denotes the most significant gene expression change.

#### 2.4 Additional results for mouse E15 neocortex dataset

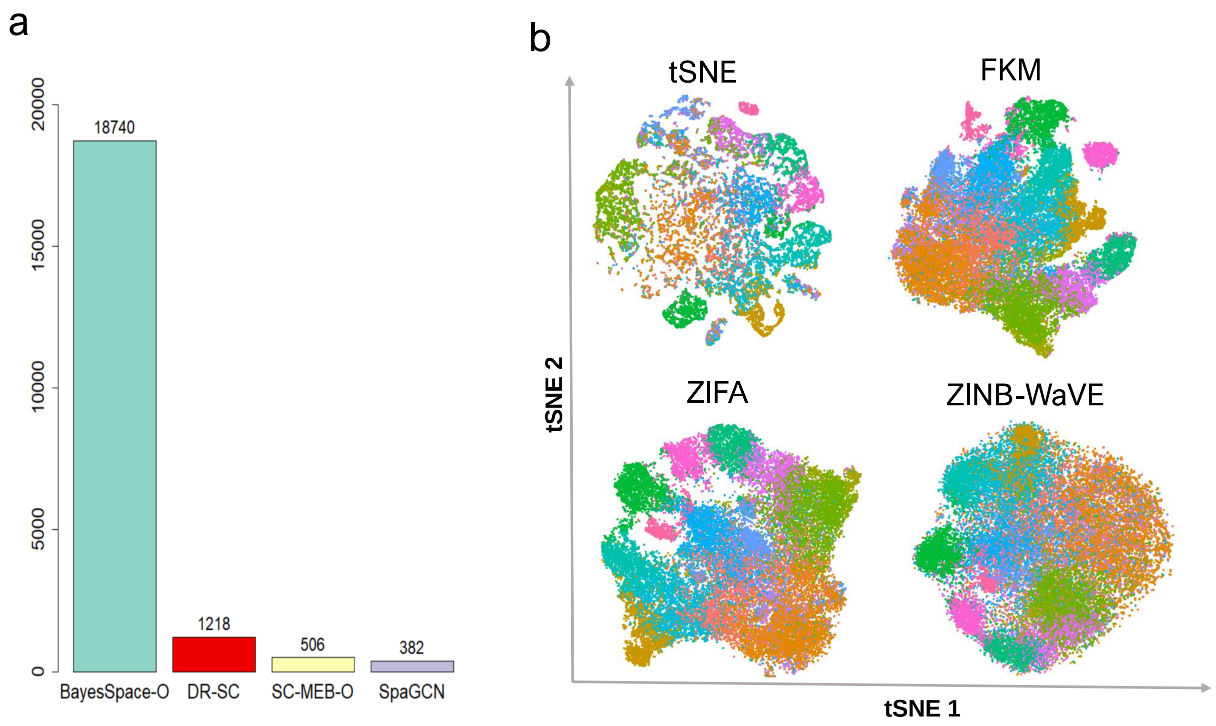

Figure S27: Analysis of mouse E15 neocortex data. a. Bar plot of running time (seconds) for DR-SC and three spatial-clustering methods, SpaGCN, SC-MEB, and BayesSpace, based on PCA. b. Visualization of cluster labels from DR-SC based on two-dimensional tSNE embeddings from four DR methods, tSNE, FKM, ZIFA and ZINB-WaVE.

#### 2.5 Additional results for mouse embryo dataset

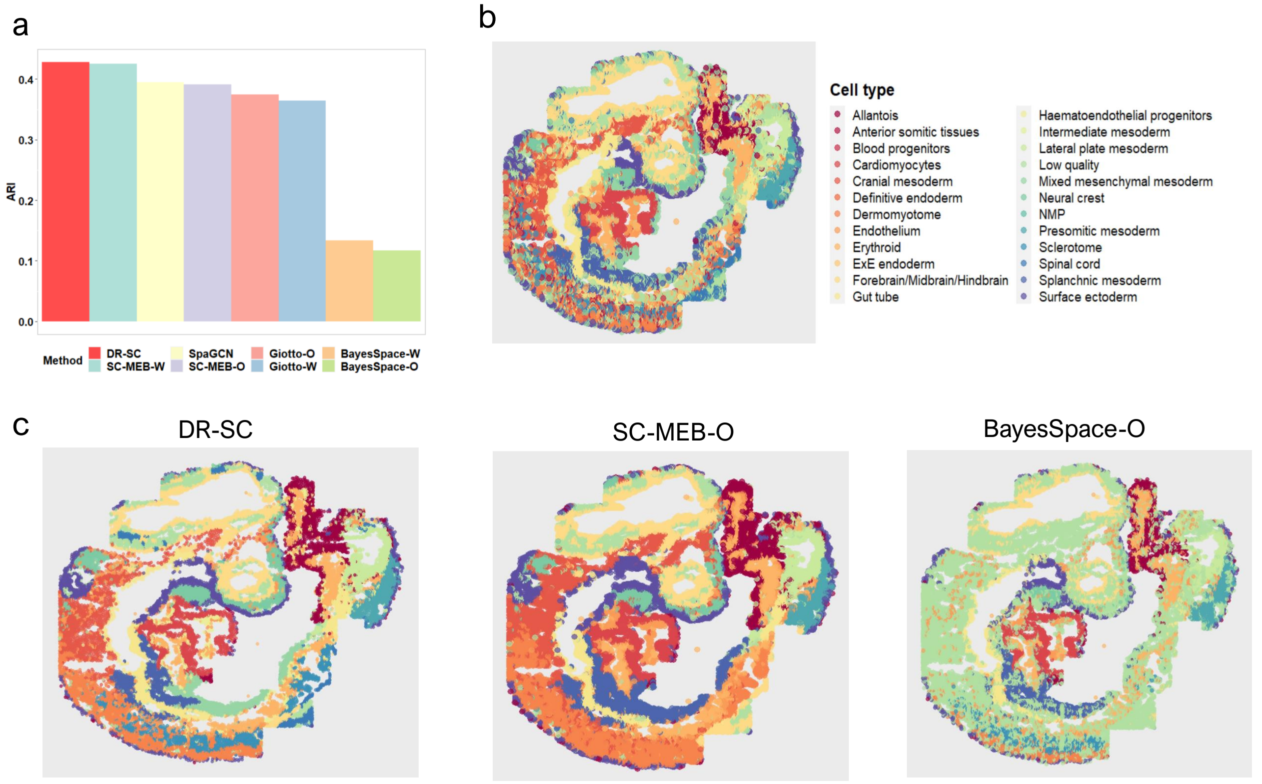

Figure S28: Comparison of clustering performance in mouse embryo dataset: a. Bar plot of ARIs for different spatial-clustering methods, DR-SC, SpaGCN, and tandem analysis based on PCA and WPCA with SC-MEB, Giotto, and BayesSpace; b. Spatial heatmap of annotated cell types; c. Spatial heatmap of cluster assignments from DR-SC, SC-MEB-O, and BayesSpace-O.
