## Supplementary Text for "Joint dimension reduction and clustering analysis for single-cell RNA-seq and spatial transcriptomics data"

**2 DR-SC facilitates lineage analysis 10**

**3 Cord blood mononuclear cells data 10**

### 1 DR-SC method

We first introduce the details of the spatial model implementation, in which the Potts model is used to model the class labels, $y_1, \cdots, y_n$, and to promote spatial smoothness within spot neighborhoods. Then, we present the details of the non-spatial model, in which the multinomial distribution is used to model the class labels, $y_1, \cdots, y_n$, and to capture the mixture proportion of each cluster. As a minor amount of revision to the spatial version produces the non-spatial version, we first present the spatial version.

#### 1.1 Pseudo full/observed log-likelihood

First, the following notations were made. Denote $[n] = \{1, \cdots, n\}$, $[n] \setminus i = \{j \leq n, j \neq i\}$ and for a set $G \subset [n]$ and a random variable $y_i$ representing the cluster label, $\mathbf{y}_G = \{y_i, i \in G\}$. Let $P(\cdot)$ and $P(\cdot|\cdot)$ be the density function and conditional density function of corresponding random variables, respectively.

To estimate the parameters of interest, we propose an expectation-maximization (EM) algorithm based on iterative conditional mode (ICM), named ICM-EM, to maximize the lower-bound function of the pseudo observed log-likelihood. Denote $\boldsymbol{\theta} = (W, \Lambda, \mu_k, \Sigma_k, k = 1, \cdots, K, \beta)$ for all parameters in the model. According to the model settings, the full likelihood can be written as

$$P(\mathbf{X}, \mathbf{Z}, \mathbf{y}) = \Pi_{i=1}^n P(\mathbf{x}_i|\mathbf{z}_i) P(\mathbf{z}_i|y_i) P(\mathbf{y}). \tag{1}$$

And the full loglikelihood can be written as

$$\ln P(\mathbf{X}, \mathbf{Z}, \mathbf{y}; \boldsymbol{\theta}) = \sum_{i=1}^n \{\ln P(\mathbf{x}_i|\mathbf{z}_i; \boldsymbol{\theta}) + \ln P(\mathbf{z}_i|y_i; \boldsymbol{\theta})\} + \ln P(\mathbf{y}; \boldsymbol{\theta}). \tag{2}$$

Note that $\mathrm{E}\{\ln P(\mathbf{y}; \boldsymbol{\theta})|\mathbf{X}\}$ in the standard EM algorithm is not tractable [1, 2], due to the spatial dependence among $\mathbf{y}$.

Following [1], suppose we have a prediction of $\mathbf{y}$, $\hat{\mathbf{y}}$, then the pseudo-likelihood of $\mathbf{y}$ is defined as

$$\tilde{P}(\mathbf{y}; \beta) = \Pi_i P(y_i|\mathbf{y}_{N_i} = \hat{\mathbf{y}}_{N_i}), \tag{3}$$

where $\mathbf{y}_{N_i}$ is the cluster label for neighbors of spot $i$. Eqn. (3) means the cluster label of each spot is only dependent on the cluster labels of its neighborhood, so the pseudo-likelihood simplifies the spatial dependence structure and makes the computation tractable. Using the joint distribution $P(\mathbf{y})$ of (3) in the main text, we showed that the conditional distribution of $y_i$ given $\mathbf{y}_{N_i}$ has the following explicit form,

$$P(y_i|\mathbf{y}_{N_i}) = C_i(\beta, \mathbf{y}_{N_i})^{-1} \exp\left\{-\sum_{i' \in N_i} \beta(1 - \delta(y_i, y_{i'}))\right\}, \tag{4}$$

where $C_i(\beta, \mathbf{y}_{N_i})$ is a normalization constant with respect to $\beta$ and $\mathbf{y}_{N_i}$. The derivation details for Eqn. (4) are given in a later section.

The pseudo-likelihood (3) takes the multiplicative form of the conditional distribution of $y_i$ given $\mathbf{y}_{[n]\backslash i} = \hat{\mathbf{y}}_{[n]\backslash i}$ for each spot $i$, making the joint distribution of $\mathbf{y}$ decomposable. This pseudo-likelihood (3) is the key to making the computation tractable. In the next subsection, we discuss how to obtain $\hat{\mathbf{y}}$ iteratively. Combing Eqns. (1) and (3), we obtain the following pseudo-complete-data likelihood

$$\tilde{P}(\mathbf{y}, \mathbf{X}, \mathbf{Z}; \boldsymbol{\theta}) = \Pi_i \left\{P(\mathbf{x}_i|\mathbf{z}_i; \boldsymbol{\theta}) P(\mathbf{z}_i|y_i; \boldsymbol{\theta}) P(y_i|\mathbf{y}_{N_i} = \hat{\mathbf{y}}_{N_i}; \boldsymbol{\theta})\right\}. \tag{5}$$

Furthermore, by integrating out $\mathbf{Z}$ and $\mathbf{y}$, we obtain the pseudo observed log-likelihood given by

$$\ln \tilde{P}(\mathbf{X}; \boldsymbol{\theta}, \hat{\mathbf{y}}) = \sum_i \ln \sum_k P(\mathbf{x}_i|y_i = k; \boldsymbol{\theta}) P(y_i = k|\mathbf{y}_{N_i} = \hat{\mathbf{y}}_{N_i}; \boldsymbol{\theta}). \tag{6}$$

Then, to estimate the model parameters, an EM algorithm is derived to maximize the lower-bound function of the pseudo observed log-likelihood.

#### 1.2 ICM-EM algorithm

To make inferences with the proposed hierarchical model, we designed an efficient EM algorithm based on the iterative conditional mode (ICM) [1, 2], named ICM-EM. This algorithm alternates between performing an ICM step, which predicts a class label $\mathbf{y}$ by maximizing its posterior, and an EM step, which computes the expectation of the log-likelihood and the model parameters $\hat{\boldsymbol{\theta}}$ by maximizing the expected log-likelihood.

##### 1.2.1 ICM step

Given the model parameters, we can derive the label predictions by maximizing the posterior distribution of $\mathbf{y}$, $P(\mathbf{y}|\mathbf{X}; \boldsymbol{\theta})$. In the Potts model, the joint distribution of $P(\mathbf{X}, \mathbf{y})$ is difficult to evaluate; thus, we apply an iterative conditional mode (ICM [1]) method to iteratively estimate $y_i$. First, note that

$$P(y_i|\mathbf{X}, \mathbf{y}_{[n]\backslash i} = \hat{\mathbf{y}}_{[n]\backslash i}) \propto P(\mathbf{x}_i|y_i) P(y_i|\mathbf{y}_{N_i} = \hat{\mathbf{y}}_{N_i}),$$

implying

$$\hat{y}_i = \arg\max_{y_i} P(\mathbf{x}_i|y_i) P(\mathbf{y}_i|\mathbf{y}_{N_i} = \hat{\mathbf{y}}_{N_i}). \tag{7}$$

In Eqn (7), $P(\mathbf{x}_i|y_i)$ is given an explicit form by integrating out $\mathbf{z}_i$ from $P(\mathbf{x}_i, \mathbf{z}_i|y_i)$, which is given by

$$P(\mathbf{x}_i|y_i = k) = (2\pi)^{-p/2} |S_k|^{1/2} \exp\left\{-\frac{1}{2}(\mathbf{x}_i - W\mu_k)^T S_k (\mathbf{x}_i - W\mu_k)\right\}, \tag{8}$$

where $S_k = \Lambda^{-1} - \Lambda^{-1}WC_k^{-1}W^T\Lambda^{-1}$ and $C_k = W^T\Lambda^{-1}W + \Sigma_k^{-1}$.

Then, given model parameters $\boldsymbol{\theta}$, solving $\hat{y}_i$ from Eqn (7) is equivalent to minimizing the energy function

$$\mathcal{E}_i(y_i) = \sum_k \delta(y_i, k)\left\{-\frac{1}{2}\ln|S_k| + \frac{1}{2}(\mathbf{x}_i - W\mu_k)^T S_k(\mathbf{x}_i - W\mu_k) + \beta \sum_{i' \in N_i}\{1 - \delta(\hat{y}_{i'}, k)\}\right\}.$$

That is,

$$\hat{y}_i \;=\; \arg\min_{y_i \in \{k : k \leq K\}} \mathcal{E}_i(y_i). \tag{9}$$

We define the responsibility that component $k$ takes for explaining the observation $\mathbf{x}_i$ as

$$R_{ik} = P(y_i = k|\mathbf{x}_i, \mathbf{y}_{[n]\setminus i} = \hat{\mathbf{y}}_{[n]\setminus i}) = \frac{P(\mathbf{x}_i|y_i = k; \boldsymbol{\theta})P(y_i = k|\mathbf{y}_{N_i} = \hat{\mathbf{y}}_{N_i})}{\sum_{k'} P(\mathbf{x}_i|y_i = k'; \boldsymbol{\theta})P(y_i = k'|\mathbf{y}_{N_i} = \hat{\mathbf{y}}_{N_i})}, \tag{10}$$

which is also the pseudo posterior probability of $y_i$ given $\mathbf{x}_i$. From this equation (10), we see how the spatial information is used. The term $P(y_i = k|\mathbf{y}_{N_i} = \hat{\mathbf{y}}_{N_i})$ captures the spatial dependence using the neighborhood information of $y_i$ to improve clustering. In the case that $y_i$ is not spatially dependent on its neighbors, the term $P(y_i = k|\mathbf{y}_{N_i} = \hat{\mathbf{y}}_{N_i})$ degenerates to the mixture proportion $P(y_i = k)$ in mixture models. In this case, $R_{ik} = \frac{P(\mathbf{x}_i|y_i=k;\boldsymbol{\theta})P(y_i=k)}{\sum_{k'} P(\mathbf{x}_i|y_i=k';\boldsymbol{\theta})P(y_i=k')}$, and the conventional EM algorithm can be used because there is no requirement for predicting $\mathbf{y}_{N_i}$. We further investigate this point in the implementation details of the non-spatial model.

##### 1.2.2 E-step

In this subsection, we derive the lower-bound function of the pseudo observed log-likelihood, called Q-function (with respect to $\boldsymbol{\theta}$). Given that the model parameters are estimated in the $t$th iteration $\boldsymbol{\theta}^{(t)}$, we have

$$\begin{aligned}
& \ln \tilde{P}(\mathbf{X}; \boldsymbol{\theta}, \hat{\mathbf{y}}) \\
=\; & \ln \int_{\mathbf{y}}\int_{\mathbf{Z}} \tilde{P}(\mathbf{y}, \mathbf{X}, \mathbf{Z}; \boldsymbol{\theta})d\mathbf{y}d\mathbf{Z} \\
=\; & \sum_i \ln \int_{y_i}\int_{\mathbf{z}_i} P(\mathbf{x}_i|\mathbf{z}_i; \boldsymbol{\theta})P(\mathbf{z}_i|y_i; \boldsymbol{\theta})P(y_i|\mathbf{y}_{N_i} = \hat{\mathbf{y}}_{N_i}; \boldsymbol{\theta})dy_i d_{\mathbf{z}_i} \\
=\; & \sum_i \ln \int_{y_i}\int_{\mathbf{z}_i} P(y_i, \mathbf{z}_i|\mathbf{x}_i, \mathbf{y}_{N_i} = \hat{\mathbf{y}}_{N_i}; \boldsymbol{\theta}^{(t)})\frac{P(\mathbf{x}_i|\mathbf{z}_i; \boldsymbol{\theta})P(\mathbf{z}_i|y_i; \boldsymbol{\theta})P(y_i|\mathbf{y}_{N_i} = \hat{\mathbf{y}}_{N_i}; \boldsymbol{\theta})}{P(y_i, \mathbf{z}_i|\mathbf{x}_i, \mathbf{y}_{N_i} = \hat{\mathbf{y}}_{N_i}; \boldsymbol{\theta}^{(t)})}dy_i d_{\mathbf{z}_i} \\
\geq\; & \sum_i \mathrm{E}_{\boldsymbol{\theta}^{(t)}}\left\{\ln \frac{P(\mathbf{x}_i|\mathbf{z}_i; \boldsymbol{\theta})P(\mathbf{z}_i|y_i; \boldsymbol{\theta})P(y_i|\mathbf{y}_{N_i} = \hat{\mathbf{y}}_{N_i}; \boldsymbol{\theta})}{P(y_i, \mathbf{z}_i|\mathbf{x}_i, \mathbf{y}_{N_i} = \hat{\mathbf{y}}_{N_i}; \boldsymbol{\theta}^{(t)})}\right\} \\
=\; & Q(\boldsymbol{\theta}; \boldsymbol{\theta}^{(t)}, \hat{\mathbf{y}}) - Q_1(\boldsymbol{\theta}^{(t)}, \hat{\mathbf{y}}),
\end{aligned}$$

where expectation $E_{\boldsymbol{\theta}^{(t)}}$ is taken with respect to $(y_i, \mathbf{z}_i)$ given $\mathbf{x}_i, \mathbf{y}_{N_i} = \hat{\mathbf{y}}_{N_i}$ and $\boldsymbol{\theta}^{(t)}$, and the inequality follows from the Jensen's inequality and the last equality holds if and only if $\boldsymbol{\theta} = \boldsymbol{\theta}^{(t)}$, where

$$Q(\boldsymbol{\theta}; \boldsymbol{\theta}^{(t)}, \hat{\mathbf{y}}) = \sum_i E_{\boldsymbol{\theta}^{(t)}} \ln\{P(\mathbf{x}_i|\mathbf{z}_i; \boldsymbol{\theta})P(\mathbf{z}_i|y_i; \boldsymbol{\theta})P(y_i|\mathbf{y}_{N_i} = \hat{\mathbf{y}}_{N_i}; \boldsymbol{\theta})\}$$

and

$$Q_1(\boldsymbol{\theta}^{(t)}, \hat{\mathbf{y}}) = \sum_i E_{\boldsymbol{\theta}^{(t)}} \ln P(y_i, \mathbf{z}_i | \mathbf{x}_i, \mathbf{y}_{N_i} = \hat{\mathbf{y}}_{N_i}; \boldsymbol{\theta}^{(t)}).$$

Next, we derive the specific form of $P(y_i, \mathbf{z}_i | \mathbf{x}_i, \mathbf{y}_{N_i} = \hat{\mathbf{y}}_{N_i}; \boldsymbol{\theta}^{(t)})$ for inducing the explicit expression of $Q(\boldsymbol{\theta}; \boldsymbol{\theta}^{(t)}, \hat{\mathbf{y}})$. Using the fact that $P(\mathbf{z}_i|\mathbf{x}_i, y_i = k) = \frac{P(\mathbf{z}_i, \mathbf{x}_i | y_i = k)}{P(\mathbf{x}_i | y_i = k)}$, we have

$$P(\mathbf{z}_i|\mathbf{x}_i, y_i = k) = (2\pi)^{-q/2} |C_k|^{1/2} \exp\left\{ -\frac{1}{2}(\mathbf{z}_i - C_k^{-1}\mathbf{v}_{ik})^T C_k (\mathbf{z}_i - C_k^{-1}\mathbf{v}_{ik}) \right\},$$

where $\mathbf{v}_{ik} = W^T \Lambda^{-1} \mathbf{x}_i + \Sigma_k^{-1} \mu_k$. Furthermore,

$$P(y_i, \mathbf{z}_i | \mathbf{x}_i, \mathbf{y}_{N_i} = \hat{\mathbf{y}}_{N_i}) = P(y_i | \mathbf{x}_i, \mathbf{y}_{N_i} = \hat{\mathbf{y}}_{N_i}) P(\mathbf{z}_i | \mathbf{x}_i, y_i) = R_{ik} P(\mathbf{z}_i | \mathbf{x}_i, y_i). \tag{11}$$

With the help of the explicit expression of $P(\mathbf{y}_i, \mathbf{z}_i | \mathbf{x}_i, \mathbf{y}_{N_i} = \hat{\mathbf{y}}_{N_i})$ in (11), we can derive the explicit form of $Q(\boldsymbol{\theta}; \boldsymbol{\theta}^{(t)}, \hat{\mathbf{y}})$, which is given by

$$\begin{aligned} Q(\boldsymbol{\theta}; \boldsymbol{\theta}^{(t)}, \hat{\mathbf{y}}) &= \sum_i \sum_k R_{ik}^{(t)} \int P(\mathbf{z}_i | \mathbf{x}_i, y_i = k; \boldsymbol{\theta}^{(t)}) \left\{ \ln P(\mathbf{x}_i | \mathbf{z}_i; \boldsymbol{\theta}) + \ln P(\mathbf{z}_i | y_i = k; \boldsymbol{\theta}) \right\} d\mathbf{z}_i \\ &+ \sum_i \sum_k R_{ik}^{(t)} \ln P(y_i = k | \mathbf{y}_{N_i} = \hat{\mathbf{y}}_{N_i}; \beta) \\ &= \sum_i (I_{i1} + I_{i2} + I_{i3}) + const, \end{aligned}$$

where $R_{ik}^{(t)}$ is the value of $R_{ik}$ when $\boldsymbol{\theta} = \boldsymbol{\theta}^{(t)}$, const is a constant term independent of parameters, and

$$I_{i1} = \sum_k R_{ik}^{(t)} \left\{ -\frac{1}{2} \ln |\Lambda_r| - \frac{1}{2} \left( \mathbf{x}_i^T \Lambda^{-1} \mathbf{x}_i + tr(W^T \Lambda^{-1} W \langle \mathbf{z}_i \mathbf{z}_i^T \rangle_k^{(t)}) - 2\mathbf{x}_i^T \Lambda^{-1} W \langle \mathbf{z}_i \rangle_k^{(t)} \right) \right\},$$

where $\langle \mathbf{z}_i \rangle_k^{(t)} = C_k^{(t),-1} \{ W^{(t),T} \Lambda^{(t),-1} \mathbf{x}_i + \Sigma_k^{(t),-1} \mu_k^{(t)} \}$, $\langle \mathbf{z}_i \mathbf{z}_i^T \rangle_k^{(t)} = C_k^{(t),-1} + \langle \mathbf{z}_i \rangle_k^{(t)} \langle \mathbf{z}_i \rangle_k^{(t),T}$,

$$I_{i2} = \sum_k R_{ik}^{(t)} \left\{ -\frac{1}{2} \ln |\Sigma_k| - \frac{1}{2} tr(\Sigma_k^{-1} \langle \mathbf{z}_i \mathbf{z}_i^T \rangle_k^{(t)}) + \mu_k^T \Sigma_k^{-1} \langle \mathbf{z}_i \rangle_k^{(t)} - \frac{1}{2} \mu_k^T \Sigma_k^{-1} \mu_k \right\},$$

and

$$I_{i3} = -\sum_k R_{ik}^{(t)} \ln C_i(\beta, \hat{\mathbf{y}}_{N_i}) - \beta \sum_k R_{ik}^{(t)} \sum_{i' \in N_i} \{ 1 - \delta(k, \hat{y}_{i'}) \}.$$

##### 1.2.3 M-step

Taking derivatives of $Q$ function with respect to each parameter in $\boldsymbol{\theta}$, we have

$$\mu_k = \frac{\sum_i R_{ik}(\boldsymbol{\theta}^{(t)}) \langle \mathbf{z}_i \rangle_k^{(t)}}{\sum_i R_{ik}(\boldsymbol{\theta}^{(t)})}, \tag{12}$$

$$\Sigma_k = \frac{\sum_i R_{ik}(\boldsymbol{\theta}^{(t)}) \left\{ \langle \mathbf{z}_i \mathbf{z}_i^T \rangle_k^{(t)} - \langle \mathbf{z}_i \rangle_k^{(t)} \mu_k^T - \mu_k \langle \mathbf{z}_i \rangle_k^{(t),T} + \mu_k \mu_k^T \right\}}{\sum_i R_{ik}(\boldsymbol{\theta}^{(t)})}, \tag{13}$$

$$W = \left\{ \sum_i \sum_k R_{ik}(\boldsymbol{\theta}^{(t)}) \mathbf{x}_i \langle \mathbf{z}_i \rangle_k^{(t),T} \right\} \left\{ \sum_i \sum_k R_{ik}(\boldsymbol{\theta}^{(t)}) \langle \mathbf{z}_i \mathbf{z}_i^T \rangle_k^{(t)} \right\}^{-1}, \tag{14}$$

$$\lambda_j = \frac{1}{n} \sum_i \sum_k R_{ik} \left( s_{1ijk}^2 + s_{2jk}^2 \right), \tag{15}$$

where $s_{1ijk}^2 = (x_{ij} - \mathbf{w}_j \langle \mathbf{z}_i \rangle_k^{(t)})^2, s_{2jk}^2 = \mathbf{w}_j C_k^{(t),-1} \mathbf{w}_j^T$ and $\mathbf{w}_j$ is the $j$-th row of $W$.

Because the derivative of $Q$ with respect to $\beta$ is complicated due to the complexity of the partition function $C_i(\beta, \hat{\mathbf{y}}_{N_i})$, we use a grid-search strategy to update $\beta$ by using the fact

$$\beta = \arg\max_{\beta} \sum_i I_{i3}.$$

Given a grid of $\beta$, such as $\{\beta_1, \cdots, \beta_S\}$, we can evaluate the value of $\sum_i I_{i3}$ for each $\beta_s, s \leq S$, then we choose the value that maximizes $\sum_i I_{i3}$.

We use the pseudo posterior expectation of $\mathbf{z}_i$ given $\mathbf{x}_i$ with predicted $\hat{\mathbf{y}}_{N_i}$ to estimate $\mathbf{z}_i$, that is

$$\begin{aligned}
& \mathrm{E}(\mathbf{z}_i | \mathbf{x}_i; \hat{\mathbf{y}}_{N_i}) \\
= & \int \mathbf{z}_i \left\{ \sum_k P(y_i = k, \mathbf{z}_i | \mathbf{x}_i, ; \hat{\mathbf{y}}_{N_i}) \right\} d\mathbf{z}_i \\
= & \sum_k \{ R_{ik} \langle \mathbf{z}_i \rangle_k \},
\end{aligned} \tag{16}$$

where $\langle \mathbf{z}_i \rangle_k = \int \mathbf{z}_i P(\mathbf{z}_i | \mathbf{x}_i, y_i = k) d\mathbf{z}_i = C_k^{-1} \mathbf{v}_{ik}$. It is clear that the dimension reduction for estimating $\mathbf{z}_i$ also uses the inferred class label information provided by $R_{ik}$. In the simulation study, we discovered that this information can improve the dimension-reduction performance.

We summarize the proposed algorithm in Algorithms 1 and 2. Algorithm 1 describes the detailed implementation of the ICM algorithm used in Algorithm 2 for predicting $\mathbf{y}$. Algorithm 2 presents details of ICM-EM algorithm. Following the suggestion of [3], we set the initial value of $\beta$ to 1.5. To get the initital values of other paramters, we perform PCA on $\mathbf{X}$ with the number of PCs $q$, then obtain loading matrix $L_1$ and score matrix $L_2$, and perform Gaussian mixture model estimation on the score matrix $L_2$, then we obtain the mean component $\tilde{\mu}_k$, covariance component $\tilde{\Sigma}_k$ and cluster labels $\tilde{\mathbf{y}}$. Finally, we initialize $\hat{\mathbf{y}}^{(0)} = \tilde{\mathbf{y}}, W^{(0)} = L_1, \mu_k^{(0)} = \tilde{\mu}_k, \Sigma_k^{(0)} = \tilde{\Sigma}_k, k \leq K$ and $\lambda_j^{(0)} = \frac{1}{n} \sum_{i=1}^n (x_{ij} - L_{2,i.} L_{1,j.}^T)^2$, where $L_{2,i.}$ is the $i$-th row of $L_2$ and $L_{1,j.}$ is the $j$-th row of $L_1$.

#### 1.3 Derivation of $P(y_i | \mathbf{y}_{N_i})$

First, $P(\mathbf{y})$ has a decomposed form,

$$P(\mathbf{y}) = \phi_1(y_i, \mathbf{y}_{N_i}) \phi_2(\mathbf{y}_{N_i}, \mathbf{y}_{[n] \backslash (i \cup N_i)}),$$

**Algorithm 1** ICM algorithm used in ICM-EM algorihtm

**Input:** $\mathbf{X}$, $\hat{\mathbf{y}}^{(t-1)}, \boldsymbol{\theta}^{(t-1)}$, $\mathcal{S} = \{s_i\}_{i=1}^n$, maximum iterations of ICM $maxIter\_ICM$, relative tolerance of difference of total energy $eps\_Eng$.

**Output:** $\hat{\mathbf{y}}^{(t)}$.

**for** each $l \in 1, \cdots, maxIter\_ICM$ **do**
  **for** each $i \in 1, \cdots, n$ **do**
    Update $\mathbf{y}_i^{(t-1),l}$ based on equation (9);
  **end for**
  Evaluate the total energy, $Eng(l)$, by (**??**).
  **if** $|Eng(l) - Eng(l-1)|/|Eng(l-1)| < eps\_Eng$ **then**
    break;
  **end if**
**end for**
$\hat{\mathbf{y}}^{(t)} = \hat{\mathbf{y}}^{(t-1),l}$;
**return** $\hat{\mathbf{y}}^{(t)}$

**Algorithm 2** The proposed ICM-EM algorithm for DR-SC

**Input:** $\mathbf{X}$, $\mathcal{S} = \{s_i\}_{i=1}^n$, $q$, $K$, grid points of $\beta$, $beta\_grid$, maximum iterations of EM $maxIter$, relative tolerance of pseudo loglikelihood $epsLogLike$, maximum iterations of ICM $maxIter\_ICM$, relative tolerance of total energy of ICM $eps\_Eng$.

**Output:** $\hat{\mathbf{y}}, \hat{\mathbf{Z}}, \hat{W}, \hat{\Lambda}, \hat{\beta}, \hat{\mu}_k, \hat{\Sigma}_k, k \leq K$

Initialize $\hat{\mathbf{y}}^{(0)}, W^{(0)}, \mu_k^{(0)}, \Sigma_k^{(0)}, k \leq K, \beta^{(0)}$ and $\Lambda^{(0)}$.
**for** each $t \in 1, \cdots, maxIter$ **do**
  Update $\hat{\mathbf{y}}^{(t)}$ based on function $ICM(\mathbf{X}, \hat{\mathbf{y}}^{(t-1)}, \boldsymbol{\theta}^{(t-1)}, \mathcal{S}, maxIter\_ICM, eps\_Eng)$;
  Update $\beta^{(t)}$ based on grid search on $beta\_grid$;
  Update $\mu_k^{(t)}$ based on Equation (12);
  Update $\Sigma_k^{(t)}$ based on Equation (13);
  Update $W^{(t)}$ based on Equation (14);
  Update $\Lambda^{(t)}$ based on Equation (15);
  Evaluate the pseudo observational loglikelihood ,$LogLike(t)$,by (6).
  **if** $|LogLike(t) - LogLike(t-1)|/|LogLike(t-1)| < epsLogLike$ **then**
    break;
  **end if**
**end for**
Evaluate $\hat{\mathbf{Z}}$ based on Equation (16) by replacing $R_{ik}$ and $\langle \mathbf{z}_i \rangle_k$ with $R_{ik}^{(t)}$ and $\langle \mathbf{z}_i \rangle_k^{(t)}$, respectively.
**return** $\hat{\mathbf{y}}, \hat{\mathbf{Z}}, \hat{W}, \hat{\Lambda}, \hat{\beta}, \hat{\mu}_k, \hat{\Sigma}_k, k \leq K$

where $\ln \phi_1(y_i, \mathbf{y}_{N_i}) = -\sum_{i' \in N_i} \beta(1 - \delta(y_i, y_{i'}))$, $\ln \phi_2(\mathbf{y}_{N_i}, \mathbf{y}_{[n]\setminus(i \cup N_i)}) = \ln P(\mathbf{y}) - \ln \phi_1(y_i, \mathbf{y}_{N_i})$. Then we can easily drive the conditional distribution given by

$$
\begin{aligned}
P(y_i|\mathbf{y}_{N_i}) &= \frac{P(y_i, \mathbf{y}_{N_i})}{P(\mathbf{y}_{N_i})} \\
&= \frac{\int P(\mathbf{y}) d\mathbf{y}_{[n]\setminus(i\cup N_i)}}{\int P(\mathbf{y}) dy_i d\mathbf{y}_{[n]\setminus(i\cup N_i)}} \\
&= \frac{\int \phi_1(y_i, \mathbf{y}_{N_i}) \phi_2(\mathbf{y}_{N_i}, \mathbf{y}_{[n]\setminus(i\cup N_i)}) d\mathbf{y}_{[n]\setminus(i\cup N_i)}}{\int \phi_1(y_i, \mathbf{y}_{N_i}) \phi_2(\mathbf{y}_{N_i}, \mathbf{y}_{[n]\setminus(i\cup N_i)}) dy_i d\mathbf{y}_{[n]\setminus(i\cup N_i)}} \\
&= \frac{\phi_1(y_i, \mathbf{y}_{N_i}) g(\mathbf{y}_{N_i})}{\int \phi_1(y_i, \mathbf{y}_{N_i}) dy_i g(\mathbf{y}_{N_i})} \\
&= \frac{\phi_1(y_i, \mathbf{y}_{N_i})}{\int \phi_1(y_i, \mathbf{y}_{N_i}) dy_i} \\
&= C_i(\beta, \mathbf{y}_{N_i})^{-1} \exp\left\{ -\sum_{i' \in N_i} \beta(1 - \delta(y_i, y_{i'})) \right\},
\end{aligned}
$$

where $g(\mathbf{y}_{N_i}) = \int \phi_2(\mathbf{y}_{N_i}, \mathbf{y}_{[n]\setminus(i\cup N_i)}) d\mathbf{y}_{[n]\setminus(i\cup N_i)}$ in the fourth equality and $C_i(\beta, \mathbf{y}_{N_i}) = \int \phi_1(y_i, \mathbf{y}_{N_i}) dy_i$ in the last equality.

#### 1.4 Implementation of non-spatial model

In the absence of spatial information, we assume a multinomial distribution for $y_i$, where each $y_i$ is independent and identically distributed, i.e. $P(y_i = k) = \pi_k$. Then the joint distribution of $\mathbf{y}$ is decomposable, which leads to the posterior expectation of the full-data log-likelihood (3) having a closed form. We only need to make small changes to the E-step and M-step for spatial implementation. In the E-step, the responsibility that component $k$ takes for explaining the observation $\mathbf{x}_i$ is changed to

$$
R_{ik} = P(y_i = k|\mathbf{x}_i) = \frac{P(\mathbf{x}_i|y_i = k; \boldsymbol{\theta}) P(y_i = k)}{\sum_{k'} P(\mathbf{x}_i|y_i = k'; \boldsymbol{\theta}) P(y_i = k')}.
$$

In the M-step, the update of smoothing parameter $\beta$ is changed to the update of mixture proportion $(\pi_k, k \leq K)$ while the updating formulae of the remaining parameters are unchanged. The formula for updating $\pi_k$ is given by

$$
\pi_k = \frac{\sum_i R_{ik}}{n}, k \leq K.
$$

Once the above two changes are made in E-step and M-step, the conventional EM algorithm can be implemented in the non-spatial model.

#### 1.5 MBIC criteria for determining $K$

Modified BIC criteria [4, 5] are used to determine the number of clusters $K$, which is given by

$$
MBIC(K) = -2 \ln P(\mathbf{X}; \widehat{\boldsymbol{\theta}}(K)) + C_n df(K) \ln n, \tag{17}
$$

where $df(K) = 1 + p(q+1) + K(q + \frac{q(q+1)}{2})$ for the spatial model, $df(K) = K + p(q+1) + K(q + \frac{q(q+1)}{2})$ for the non-spatial model, and $C_n$ is a positive constant that can depend on $n$ and $p$. When $C_n = 1$, the modified BIC reduces to the traditional BIC [6]. Following the lead of [5], we take the same strategy and let $C_n = c\ln(\ln(p+n))$, where $c$ is a positive constant default as 1. This modified BIC is proposed for use in high-dimensional data settings to select tuning parameters, and is more suitable than conventional BIC criteria for our proposed model involving high-dimensional gene expression. As the observed log-likelihood $\ln P(\mathbf{X}; \widehat{\boldsymbol{\theta}}(K))$ is intractable in Enq. (17), it is approximated by the pseudo observed log-likelihood $\ln \tilde{P}(\mathbf{X}; \widehat{\boldsymbol{\theta}}(K), \hat{\mathbf{y}}(K))$ in Enq. (6).

#### 1.6 Details of evaluation metrics

In the simulation, the adjusted Rand index (ARI) and normalized mutual information (NMI) were used to measure the similarity between the estimated partition and the true one. CCor was used to measure the similarity of the extracted latent features and the true one. Concor was used to assess the remaining associations between gene expression and cell type by excluding the effects of the extracted latent features. In the analysis of the CBMC, DLPFC and mouse embryo datasets, manual annotations based on additional experiments and computational results were available. ARI and NMI were used to measure the similarity between the labels from the estimated partition and the manually annotated clusters.

To compare the selection of cluster number, we compared the consistency of the chosen cluster number with the true one. Finally, for computational speed, we recorded the computational time for each method.

Finally, we introduced the metrics used for the trajectory inference for the 16 benchmark datasets. Following information in the trajectory inference-related literature [7, 8], we used Spearman correlation and Kendall correlation to measure the correlation between the true trajectory time $\mathbf{t} = (t_1, \cdots, t_n)$ and the inferred pseudotime $\hat{\mathbf{t}} = (\hat{t}_1, \cdots, \hat{t}_n)$ as both of these metrics are rank-based methods and focus on the order of time. The Spearman correlation $\rho$ and the Kendall correlation $\tau$ are calculated as follows:

$$\begin{aligned}
\rho &= \frac{cov(rg_{\mathbf{t}}, rg_{\hat{\mathbf{t}}})}{\sigma_{rg_{\mathbf{t}}}\sigma_{rg_{\hat{\mathbf{t}}}}}, \\
\tau &= \frac{1}{n(n-1)}\sum_{i \neq j} sgn(t_i - t_j)sgn(\hat{t}_i - \hat{t}_j),
\end{aligned}$$

where $rg_{\mathbf{t}}$ and $rg_{\hat{\mathbf{t}}}$ are the rankings of cells/spots, $cov(rg_{\mathbf{t}}, rg_{\hat{\mathbf{t}}})$ is the covariance of rank variables, $\sigma_{rg_{\mathbf{t}}}$ and $\sigma_{rg_{\hat{\mathbf{t}}}}$ are the standard deviations of the rank variables, and $sgn(t_i - t_j)$ is the sign of $t_i - t_j$. It is worth noting that although the true lineage is linear without any bifurcation or multifurcation patterns, the inferred result may contain multiple ending points for the datasets used. Thus, for each estimated lineage, we calculated the corresponding correlations coefficients ($\rho$ and $\tau$) for one trajectory at a time, then took the average or maximum of the coefficients over all lineages as the final metrics. Therefore, there are four metrics, the mean/maximum Spearman correlations and the mean/maximum Kendall correlations. In the Slingshot [7], the authors only used the maximum Kendall correlation to measure the performance of the inferred trajectory but they also pointed out that the maximum version produces a bias in favor of methods that identify many, potentially spurious lineages. This is why we also used the average version.

### 2 DR-SC facilitates lineage analysis

We used the 16 scRNA-seq datasets with known lineage information (Supplementary Table S3) as the baseline to show that the embeddings and cell labels estimated by DR-SC facilitate the inference for cell lineage. The known lineage information in all datasets is linear, with no bifurcation or multifurcation patterns. We inferred cell lineage based on reduced-dimensionality space estimated by both DR-SC and eight other dimension reduction methods. In all methods except for tSNE, we varied the number of low-dimensional components from 5, 10 to 15 [9] to examine their influence on the downstream lineage analysis while the number of low-dimensional components for tSNE was fixed at three [10]. With the estimated embeddings from DR-SC and the other eight methods, we applied Slingshot [7] to perform the downstream trajectory inference since it was shown to provide superior performance to other trajectory inference methods in terms of accuracy, scalability, stability and usability [11]. In the trajectory inference, cell labels were required for the identification of lineages and branching events. For joint methods, i.e., DR-SC and FKM, the estimated features and cell labels were simultaneously identified, while FKM used the number of clusters from DR-SC due to its limitation in determining $K$. For all other methods, we applied GMM to perform clustering analysis using the corresponding estimated embeddings. Supplementary Fig. S1 shows the mean and maximum Kendall's rank correlation coefficients between the inferred and true pseudotime variables among all 16 datasets. We observed that Kendall's rank correlation coefficients were quite stable for DR-SC with various numbers of low-dimensional components, but they fluctuated much more for all the other methods. Supplementary Fig. S2 shows the mean and maximum Spearman's rank correlation coefficients between the inferred and true pseudotime variables among all 16 datasets, and a similar pattern could be observed. Supplementary Fig. S3 shows the number of lineages inferred from the low-dimensional representations estimated from different methods. We found that only analysis using embeddings from DR-SC recovered the underlying true number of cell lineages while the analysis using embeddings from all other methods tended to over-estimate the number of cell lineages.

### 3 Cord blood mononuclear cells data

CITE-seq technology combines the highly multiplexed antibody-based detection of protein markers with transcriptome profiling at the single-cell resolution [12]. In this analysis, we applied DR-SC and other methods to identify and visualize cell clusters using only single-cell transcriptome profiles from a CITE-seq dataset for cord blood mononuclear cells (CBMCs). Since cell-surface proteins are routinely used as markers to classify immune cells [13], we ran clustering analysis based on cell-surface markers only and took this as the benchmark. The clusters identified using cell-surface markers were further refined using the RNA expression data for those cells, e.g., in distinguishing dendritic cells (DC) from CD14+ monocytes (CD14+ Mono) [12]. Details of clustering analysis using cell-surface markers are given in the Materials and Methods.

Using only single-cell transcriptome profiles in this dataset, we first applied DR-SC and the other eight dimension reduction methods to extract 25-dimensional embeddings [12] and further applied tSNE to reduce these 25-dimensional embeddings to a two-dimensional representation for each method. DR-SC obtained the class labels by simultaneously conducting dimension reduction and clustering, while the other dimension-reduction methods estimated the class labels using a Gaussian mixture model (GMM). Supplementary Fig. S4a and Fig. S5a show the tSNE plots of the two-dimensional representation obtained from DR-SC and the other eight methods. We found that the tSNE plot based on DR-SC separated clusters more clearly compared to other methods.

Taking the clustering results obtained using only cell-surface markers as ground truth (see Methods), we evaluated the ARI for the different clustering methods, including the tandem analysis methods, such as GMM, $k$-means, Leiden and Louvain, and joint methods, such as FKM and ORCLUS. Supplementary Fig. S4b shows the clustering performance of DR-SC and GMM using low-dimensional features obtained from the other seven dimension reduction methods. DR-SC achieved a higher ARI value than GMM using other dimension reduction methods. The clustering performances of $k$-means, Leiden and Louvain in the tandem analysis and FKM and ORCLUS in the joint analysis are shown in Supplementary Fig. S6b. Among all methods considered, DR-SC achieved the highest ARI value. To further visualize the separability of the clustering using transcriptome profiles, we showed both the heatmap and ridge plot of the protein levels on RNA clusters from DR-SC (Supplementary Fig. S7 and S8). The separability of cell types using protein levels was consistent with the tSNE plot for embeddings estimated by DR-SC (Supplementary Fig. S4a).

Next, we performed differential gene expression (DGE) analysis for nine human cell types identified by DR-SC. Supplementary Fig. S6c&d show the heatmaps of the normalized expression of genes identified in the DGE analysis and all 10 surface proteins, respectively. These results provided an indication of the separability of the inferred cell types using DR-SC. In the DGE analysis, we identified 14, 171, 34, 87, 105, 326, 102, 67 and 16 differentially expressed genes in CD34+ cell, CD14+ monocytes (CD14+ Mono), CD4 T, CD 16+ monocytes (CD16+ Mono), dendritic cell (DC), erythroid-like and erythroid precursor cells (erythroid-like), natural killer (NK), B cells, and CD8 T cells, respectively, with a false discovery rate (FDR) of less than 0.05 and a log-fold change greater than 0.5. Details of all identified differentially expressed genes are provided in Supplementary Table S4. Then, we performed functional enrichment analysis of these differentially expressed genes. A total of 885 and 188 respective Gene Ontology (GO) terms were enriched for CD14+ Mono and NK cells with adjusted $p$-values of less than 0.05. Supplementary Fig. S4c shows the bubble plots of $-\log 10(p-\text{values})$ for these two cell types, in which immune-related pathways were enriched for both CD14+ Mono and NK cells. Both CD14+ Mono and NK cells play crucial roles in the processes of infectious diseases [14, 15]. The results of the functional enrichment analysis for CD16+ Mono, DC cells, erythroid-like cells and B cells are provided in Supplementary Fig. S5b and S6a. The most significantly enriched pathways for DC and CD16+ Mono cells included immune system process. DC cells are best known as antigen-presenting cells that initiate and regulate immune responses [16, 17], while it has been shown that a subset of CD16+ Mono exhibits phenotypical and functional DC cells characteristics with various abilities to stimulate CD4+ T cells in the immune system [18]. The top significantly enriched pathways for erythroid-like cells included hemoglobin complex and oxygen/gas transport processes.
